# Genome-wide Association Study of Multisite Chronic Pain in UK Biobank

**DOI:** 10.1101/502807

**Authors:** Keira J.A. Johnston, Mark J. Adams, Barbara I. Nicholl, Joey Ward, Rona J Strawbridge, Amy Ferguson, Andrew McIntosh, Mark E.S. Bailey, Daniel J. Smith

## Abstract

Chronic pain is highly prevalent worldwide, contributing a significant socioeconomic and public health burden. Several aspects of chronic pain, for example back pain and a severity-related phenotype, chronic pain grade, have been shown to be complex, heritable traits with a polygenic component. Additional pain-related phenotypes capturing aspects of an individual’s overall sensitivity to experiencing and reporting chronic pain have also been suggested. We have here made use of a measure of the number of sites of chronic pain in individuals within the general UK population. This measure, termed Multisite Chronic Pain (MCP), is also a complex trait, but its genetic architecture has not previously been investigated. To address this, a large-scale genome-wide association study (GWAS) of MCP was carried out in ~380,000 UK Biobank participants to identify associated genetic variants. Findings were consistent with MCP having a significant polygenic component with a SNP heritability of 10.2%, and 76 independent lead single nucleotide polymorphisms (SNPs) at 39 risk loci were identified. Additional gene-level association analyses identified neurogenesis, synaptic plasticity, nervous system development, cell-cycle progression and apoptosis genes as being enriched for genetic association with MCP. Genetic correlations were observed between MCP and a range of psychiatric, autoimmune and anthropometric traits including major depressive disorder (MDD), asthma and BMI. Furthermore, in Mendelian randomisation (MR) analyses a bi-directional causal relationship was observed between MCP and MDD. A polygenic risk score (PRS) for MCP was found to significantly predict chronic widespread pain (pain all over the body), indicating the existence of genetic variants contributing to both of these pain phenotypes. These findings support the proposition that chronic pain involves a strong nervous system component and have implications for our understanding of the physiology of chronic pain and for the development of novel treatment strategies.

## Introduction

Chronic pain, conventionally defined as pain lasting longer than 3 months, has high global prevalence (~30%) (Elzahaf *et al.*, 2012), imposes a significant socioeconomic burden, and contributes to excess mortality (Hocking *et al.*, 2012; Vos *et al.*, 2015). It is often associated with both specific and non-specific medical conditions such as cancers, HIV/AIDS, fibromyalgia and musculoskeletal conditions (Merskey and Bogduk, 1994; Greene, 2010; Vellucci, 2012), and can be classified according to different grading systems, such as the Von Korff chronic pain grade (Von Korff *et al.*, 1992). Several aspects of chronic pain, such as chronic pain grade and back pain, have been studied from the genetic point of view, and several have been shown to be complex traits with moderate heritability (Hocking *et al.*, 2012; McIntosh *et al.*, 2016). In part due to the heterogeneity of pain assessment and pain experience, there are very few large-scale genetic studies of chronic pain and no genome-wide significant genetic variants have yet been identified (Mogil, 2012; Zorina-Lichtenwalter *et al.*, 2016).

Chronic pain and chronic pain disorders are often comorbid with psychiatric and neurodevelopmental disorders (Gureje *et al.*, 2008). The immune and nervous systems play a central joint role in chronic pain development (Pinho-Ribeiro, Verri and Chiu, 2017; Kwiatkowski and Mika, 2018). Similarly, obesity and chronic pain are often comorbid, with lifestyle factors such as MDD and sleep disturbance also impacting on chronic pain (Okifuji and Hare, 2015; Paley and Johnson, 2016). Sleep changes and loss of circadian rhythm is common in those with chronic pain (Alföldi, Wiklund and Gerdle, 2014). Chronic pain is also a common component of many neurological diseases (Borsook, 2012).

The relationship between injury and other peripheral insult, consequent acute pain and the subsequent development of chronic pain has not been fully explained. Not everyone who undergoes major surgery or is badly injured will develop chronic pain, for example (Denk, McMahon and Tracey, 2014), and the degree of joint damage in osteoarthritis is not related to chronic pain severity (Trouvin and Perrot, 2018). Conversely, Complex Regional Pain Syndrome (CRPS) can be incited by minor peripheral insult such as insertion of a needle (reviewed by Denk, McMahon and Tracey, 2014).

Structural and functional changes in the brain and spinal cord are associated with the development and maintenance of chronic pain, and affective brain regions are involved in chronic pain perception (this is in contrast to acute pain and even to prolonged acute pain experience) (Hashmi *et al.*, 2013; Mansour *et al.*, 2013; Baliki *et al.*, 2014; Baliki and Apkarian, 2015; Bliss *et al.*, 2016). It is also unlikely that there are legitimate cut-off points or thresholds for localised and widespread chronic pain, with pain instead existing on a “continuum of widespreadness” (Kamaleri *et al.*, 2008). It may, therefore, be more valuable and powerful to examine measures of chronic pain as complex neuropathological traits in themselves, rather than genome-wide study of disorders with chronic pain as a main feature, injuries and events that tend to incite chronic pain, or specific bodily locations.

## Methods

To investigate the underlying genetic architecture of chronic pain, we carried out a GWAS of Multisite Chronic Pain (MCP), a derived chronic pain phenotype, in 387, 649 UK Biobank participants (Table 1). UK Biobank is a general-population cohort of roughly 0.5 million participants aged 40-79 recruited across the UK from 2006-2010. Details on phenotyping, follow-up and genotyping have been described in detail elsewhere (Sudlow *et al.*, 2015).

**Table 1:** Demographics of those included in BOLT-LMM GWAS of MCP.

| chronic pain sites | male (N) | female (N) | male (%) | female (%) | age (mean) | total (N) | total (%) |
| --- | --- | --- | --- | --- | --- | --- | --- |
| 0 | 105474 | 113148 | 48.2 | 51.8 | 56.71 | 218622 | 56.40 |
| 1 | 42734 | 49984 | 46.1 | 53.9 | 57.03 | 92718 | 23.92 |
| 2 | 18612 | 26000 | 41.7 | 58.3 | 57.29 | 44612 | 11.51 |
| 3 | 7771 | 12376 | 38.6 | 61.4 | 57.65 | 20147 | 5.20 |
| 4 | 2970 | 5319 | 35.8 | 64.2 | 57.48 | 8289 | 2.14 |
| 5 | 780 | 1723 | 31.2 | 68.8 | 56.53 | 2503 | 0.65 |
| 6 | 181 | 471 | 27.8 | 72.2 | 56.20 | 652 | 0.17 |
| 7 | 34 | 72 | 32.1 | 67.9 | 56.17 | 106 | 0.03 |
| total | 178556 | 209093 | NA | NA | 56.91 | 387649 | NA |

**Table 2:**
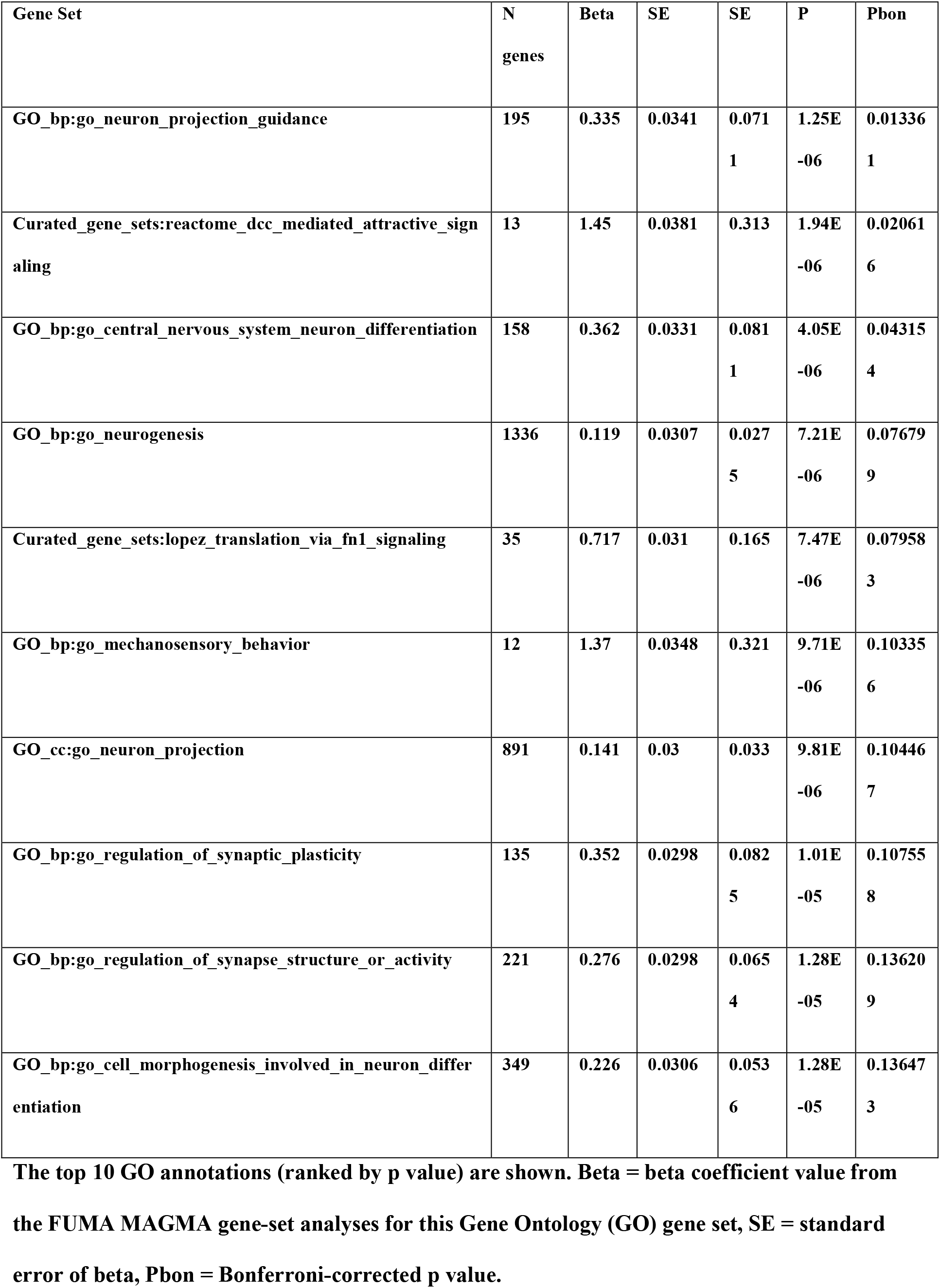
GO Annotations.

### BOLT-LMM GWAS

During the baseline investigations, UK Biobank participants were asked via a touchscreen questionnaire about “pain types experienced in the last month” (field ID 6159), with possible answers: ‘None of the above’; ‘Prefer not to answer’; pain at seven different body sites (head, face, neck/shoulder, back, stomach/abdomen, hip, knee); or ‘all over the body’. The seven individual body-site pain options were not mutually exclusive and participants could choose as many as they felt appropriate. Where patients reported recent pain at one or more body sites, or all over the body, they were additionally asked (category ID 100048) whether this pain had lasted for 3 months or longer. Those who chose ‘all over the body’ could not also select from the seven individual body sites.

Multisite Chronic Pain (MCP) was defined as the sum of body sites at which chronic pain (at least 3 months duration) was recorded: 0 to 7 sites. Those who answered that they had chronic pain ‘all over the body’ were excluded from the GWAS as there is some evidence that this phenotype relating to widespread pain can be substantially different from more localised chronic pain (Nicholl *et al.*, 2014) and should not, therefore, be considered a logical extension of the multisite scale. 10,000 randomly-selected individuals reporting no chronic pain were excluded from the GWAS to use as controls in subsequent PRS analyses.

SNPs with an imputation quality score of less than 0.3, Minor Allele Frequency (MAF), < 0.01 and Hardy-Weinberg equilibrium (HWE) test p < 10^−6^ were removed from the analyses. Participants whose self-reported sex did not match their genetically-determined sex, those who had putative sex-chromosome aneuploidy, those considered outliers due to missing heterozygosity, those with more than 10% missing genetic data and those who were not of self-reported white British ancestry were excluded from analyses.

An autosomal GWAS was run using BOLT-LMM (Loh *et al.*, 2015), with the outcome variable, MCP, modelled as a linear quantitative trait under an infinitesimal model, and the model adjusted for age, sex and chip (genotyping array). The summary statistics from the GWAS output were analysed using FUMA (Watanabe *et al.*, 2017), which implements a number of the functions from MAGMA (gene-based association testing, gene-set analyses) tissue expression (GTeX) analyses (Aguet *et al.*, 2017) and Gene Ontology annotation (de Leeuw *et al.*, 2015), and ANNOVAR annotation (Wang, Li and Hakonarson, 2010) was used to characterise lead SNPs further. LocusZoom (Pruim *et al.*, 2010) was used to plot regions around independently (of FUMA)-identified lead SNPs (N = 47) (Supplementary Information).

#### Genetic Correlation Analyses

Genetic correlations between MCP and 22 complex traits selected in the basis of prior association evidence were calculated using linkage disequilibrium score regression (LDSR) analyses (Bulik-Sullivan *et al.*, 2015), either implemented using the ‘ldsc’ package (Bulik-Sullivan *et al.*, 2015) and downloaded publicly-available summary statistics or using summary statistics from in-house analyses or carried out using LD Hub (Zheng *et al.*, 2017). LD Hub datasets from the categories Psychiatric, Personality, Autoimmune and Neurological were selected and datasets with the attached warning note ‘Caution: using this data may yield less robust results due to minor departure from LD structure’ were excluded from the analyses. Where multiple GWAS datasets were available for the same trait, the one with the largest sample size and/or European ancestry was retained with priority given to European ancestry.

#### Mendelian Randomisation of MCP and Major Depressive Disorder

Mendelian randomisation analysis was carried out with MR-RAPS (MR-Robust Adjusted Profile Score; (Zhao *et al.*, 2018) using the R package ‘mr-raps’. This method is appropriate when doing MR analysis of phenotypes that are moderately genetically correlated and likely to share some pleiotropic risk loci. Summary statistics from the most recent MDD GWAS meta-analysis (Wray *et al.*, 2018), with UK Biobank and 23andMe results removed, were harmonised with MCP GWAS summary statistics following guidelines by Hartwig et al as closely as possible with the available data (Hartwig *et al.*, 2016), and also following Zhao et al to allow MR-RAPS analyses (Zhao *et al.*, 2018). Bi-allelic SNPs shared between the two datasets were identified and harmonised (by ‘flipping’) with respect to the strand used to designate alleles. (Hartwig *et al.*, 2016). Reciprocal MR analysis was carried out using subsets of SNPs associated with each of the exposure traits (MCP and MDD) at p < 10^−5^. was ensured that the effect allele was trait-increasing in the exposure trait, and that the effect allele matched between the exposure and the outcome. These selected subsets of variants were then LD-pruned at a threshold of r^2^ < 0.01 using command-line PLINK using ‘indep-pairwise’ with a 50-SNP window and sliding window of 5 SNPs (Purcell *et al.*, 2007). This resulted in a set of 200 instruments for MCP as the exposure, and a set of 99 instruments for MDD as the exposure.

#### PRS Prediction of Chronic Widespread Pain

Those who reported chronic pain all over the body were excluded from the MCP GWAS analyses above. This is because chronic pain all over the body, taken as a proxy for chronic widespread pain (CWP), may be a different clinical syndrome from more localised chronic pain, and does not necessarily directly reflect chronic pain at 7 bodily sites. To investigate the relationship between CWP and MCP, a PRS approach was taken.

A polygenic risk score (PRS) was constructed for MCP in individuals who reported chronic pain all over the body (n = 6, 815; these individuals had all been excluded from the MCP GWAS), and in controls (n = 10, 000 individuals reporting no chronic pain at any site, also excluded from the MCP GWAS). The PRS was calculated using SNPs associated with MCP at p < 0.01, weighting by MCP GWAS effect size (GWAS β) for each SNP. A standardised PRS (Z-scores) was used in all analyses, constructed by dividing the calculated PRS by its standard deviation across all samples. The ability of the standardised PRS to predict chronic widespread pain status was investigated in logistic regression models adjusted for age, sex, genotyping array and the first 8 genetic principal components.

## Results

### BOLT-LMM GWAS

Genetic loci influencing MCP level, and thus number of chronic pain sites reported, were identified using a GWAS approach. No evidence was found for inflation of the test statistics due to hidden population stratification (λ_GC_ = 1.26; after adjustment for sample size λ_GC_1000 = 1.001). LDSR analysis was consistent with a polygenic contribution to MCP (1.0249 (SE = 0.0274); Figure 1) and yielded a SNP heritability estimate of 10.2%. BOLT-LMM gave a similar SNP heritability estimate (pseudo-h^2^ = 10.3%). In total, 1, 748 SNPs associated with MCP level at genome-wide significance (p < 5 x 10^−8^) were identified. Conditional analysis of the association signals at each locus revealed 76 independent genome-wide significant lead SNPs (Supplementary Figures across 39 risk loci (Table 4)

**Figure 1:**
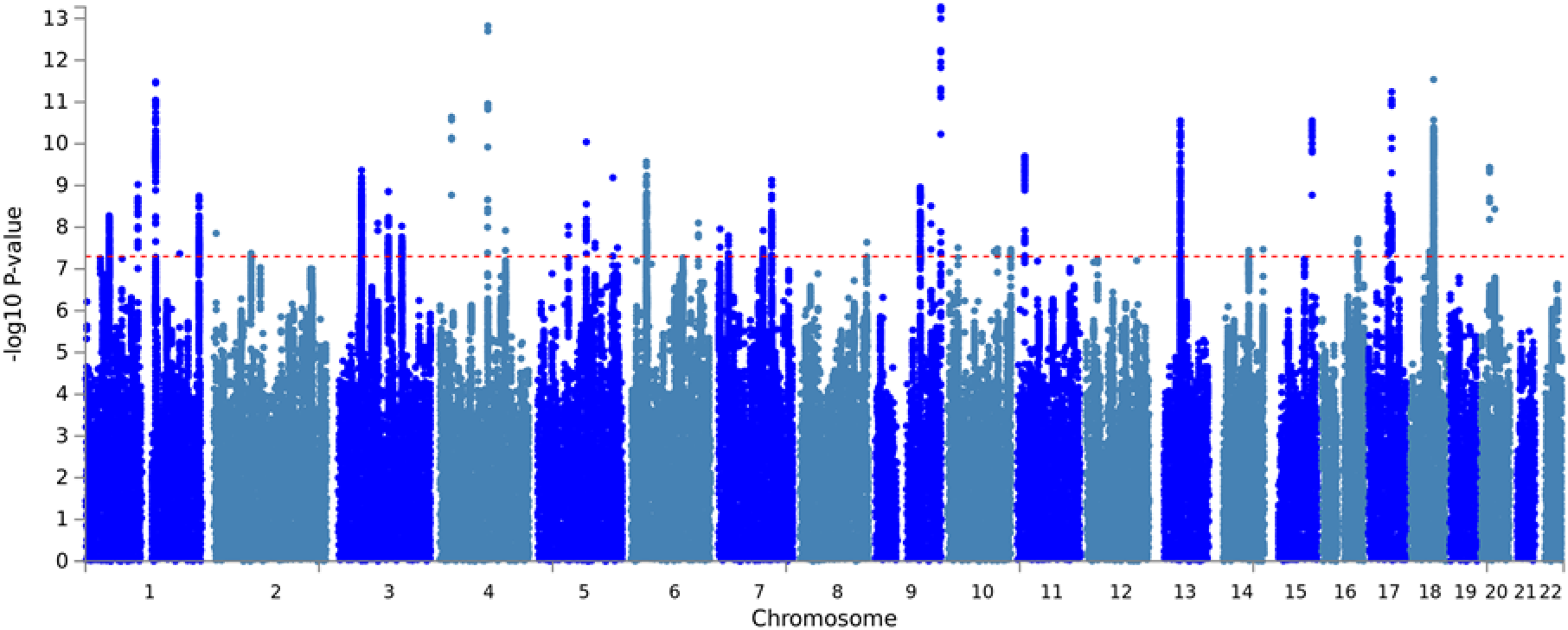
Manhattan Plot for MCP GWAS.

**Figure 2:**
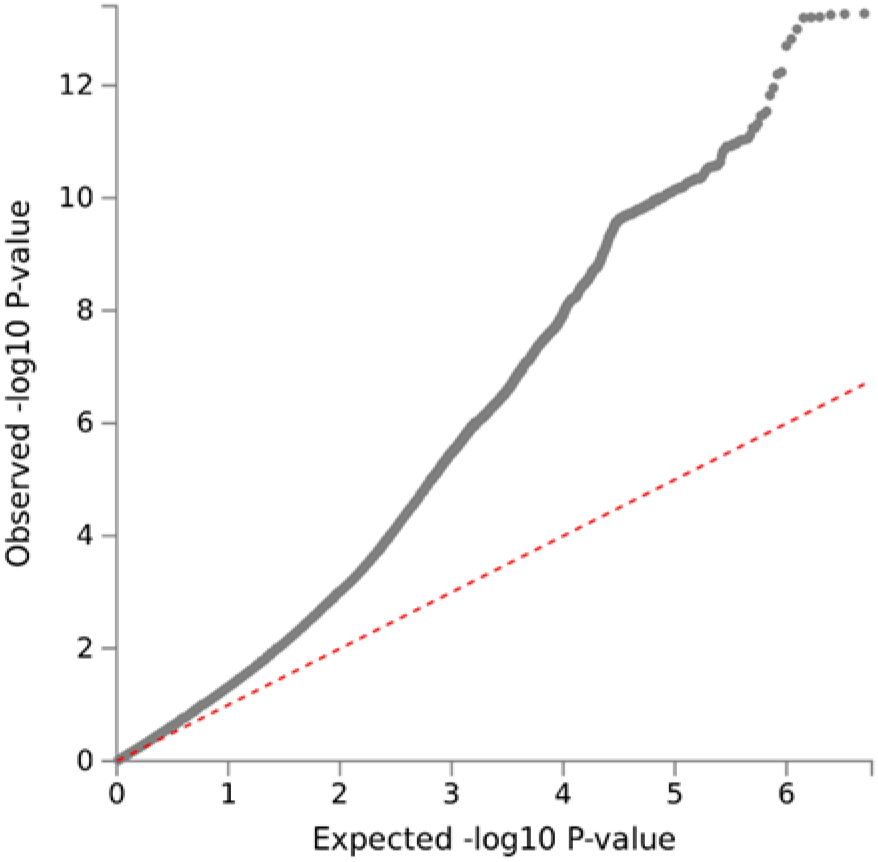
MCP GWAS QQ Plot.

**Figure 3:**
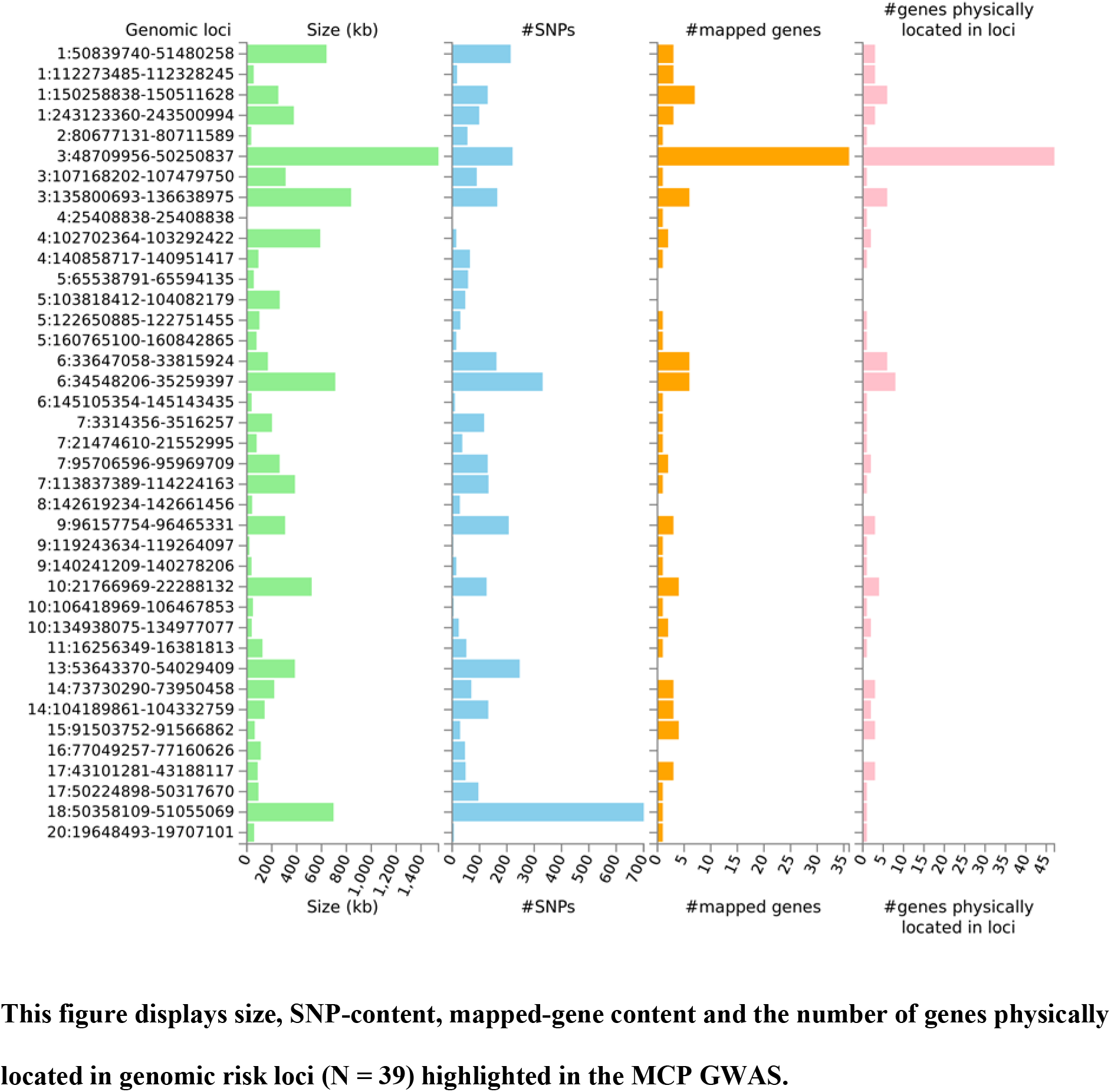
Summary of findings at the identified loci.

GWAS results for individual SNPs were integrated in gene-level association tests (MAGMA gene-based test), which revealed 143 genes significantly associated with MCP (Figure 4).

**Figure 4:**
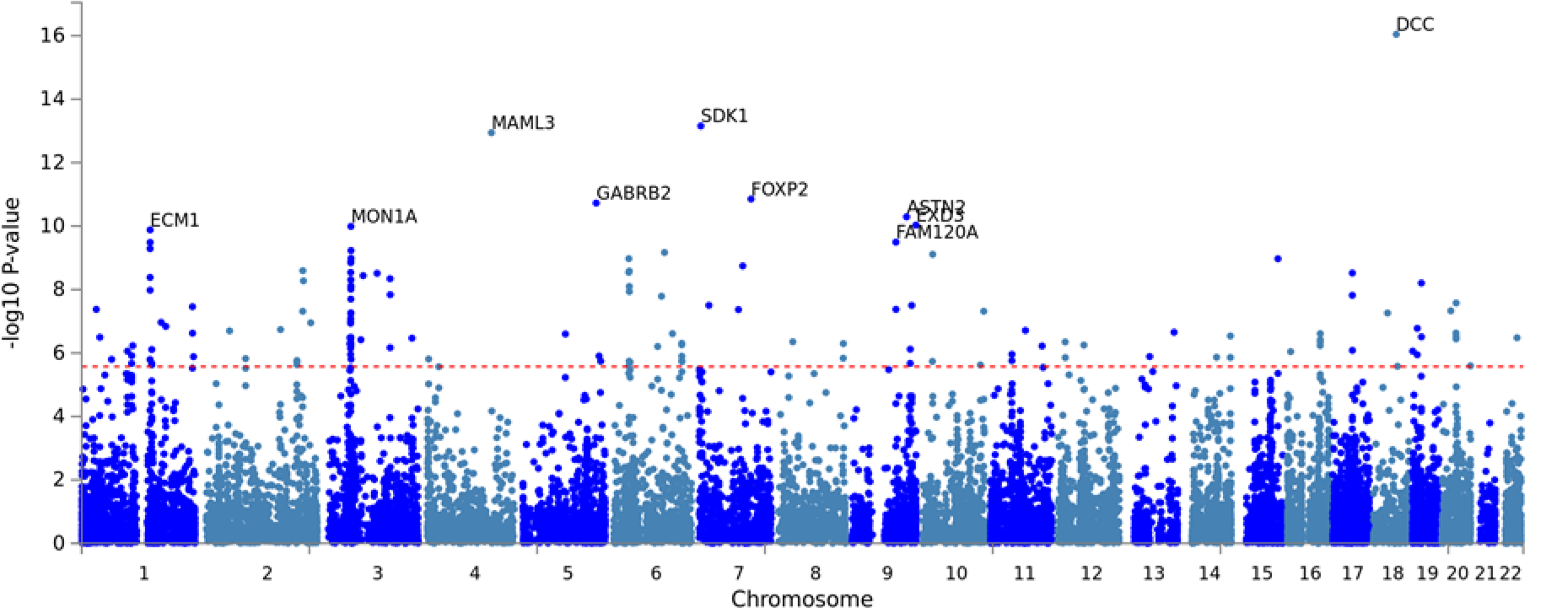
Gene-Based Test (MAGMA) Manhattan Plot. Results of the MAGMA gene-based test results implemented via FUMA are shown, with the top 10 most-significant gene associations labelled.

Gene Ontology (GO) annotations involved neurogenesis and synaptic plasticity, with significant category annotations (Bonferroni-corrected p < 0.05) being DCC-mediated attractive signalling, neuron projection guidance and central nervous system neuron differentiation.

**Figure 5:**
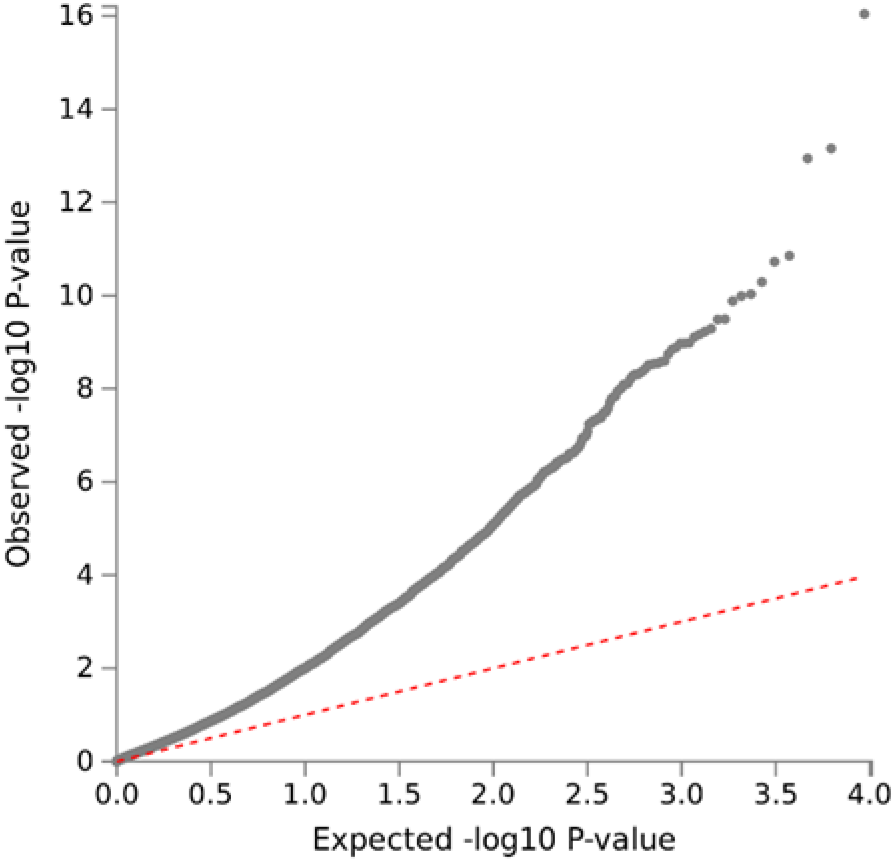
Gene-Based Test QQ Plot.

Tissue expression analyses showed biased expression in the brain (Fig. 6a), particularly in the cortex and cerebellum (Fig. 6b).

**Figure 6a:**
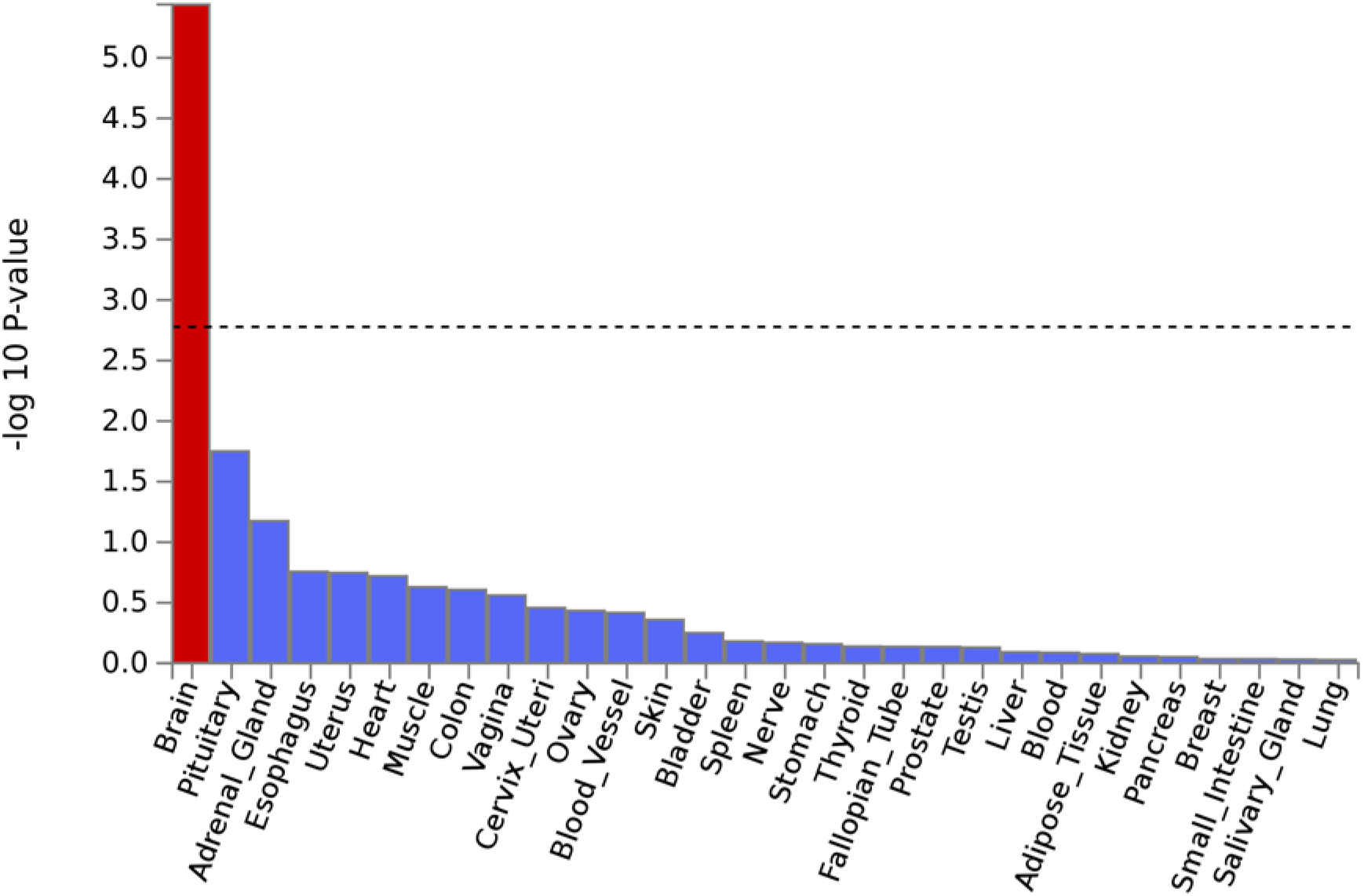
GteX Output – General Tissues.

**Figure 6b:**
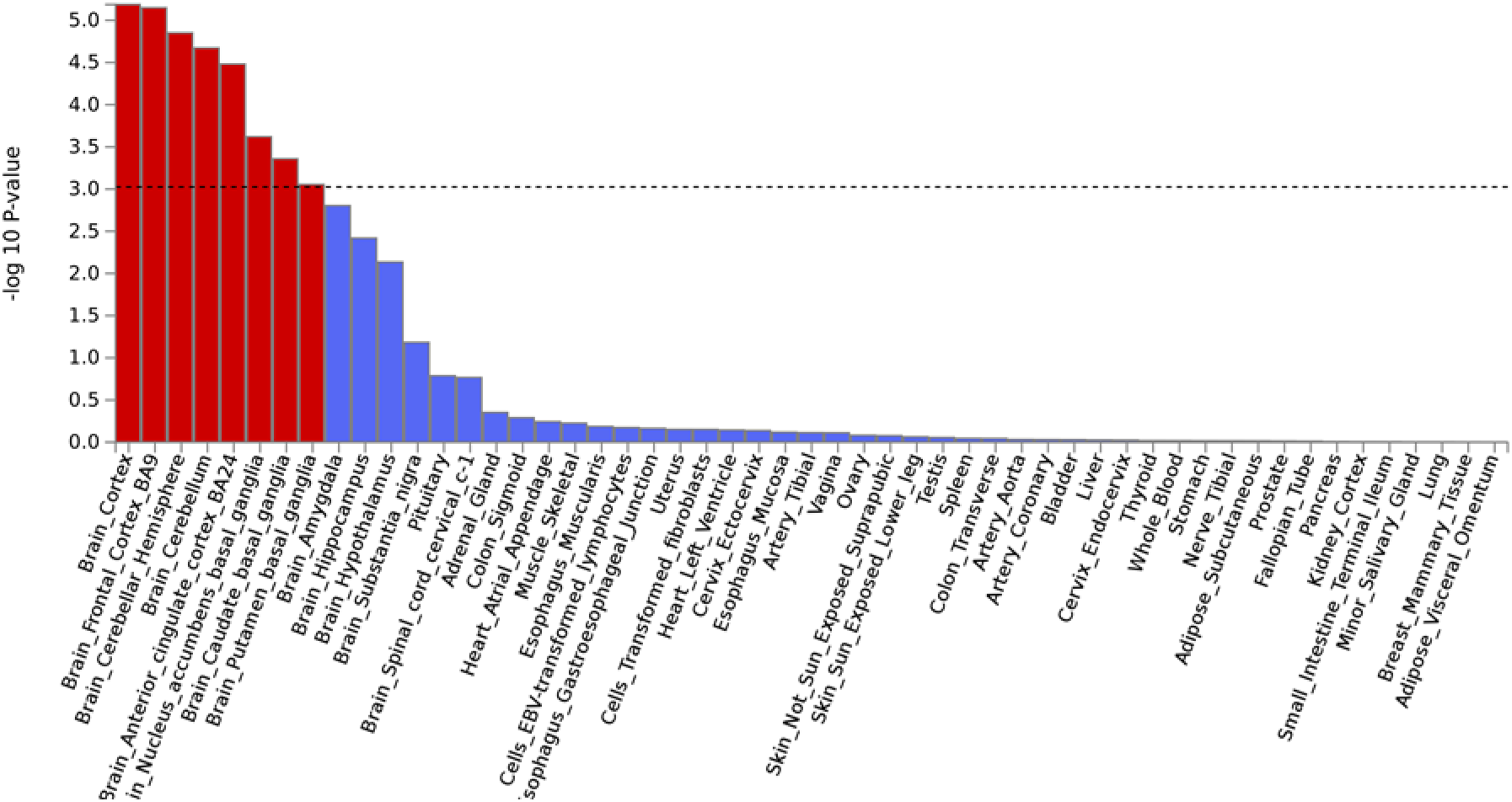
GteX Output – Detailed Tissues.

39 genomic risk loci were found for MCP (Table 3), Genomic risk loci as defined by FUMA via independent lead SNPs and the maximum distance between their LD block (Watanabe *et al.*, 2017), and were found on chromosomes 1-11, 13-18 and chromosome 20. 35 genes of interest were chosen (Table 4, Supplementary Information).

**Table 3:** Genomic Risk Loci.

| GenomicLocus | rsID | chr | pos |
| --- | --- | --- | --- |
| 1 | rs10888692 | 1 | 50991473 |
| 2 | rs197422 | 1 | 112317512 |
| 3 | rs59898460 | 1 | 150493004 |
| 4 | rs12071912 | 1 | 243241614 |
| 5 | rs4852567 | 2 | 80703379 |
| 6 | rs7628207 | 3 | 49754970 |
| 7 | rs28428925 | 3 | 107294634 |
| 8 | rs6770476 | 3 | 136073920 |
| 9 | rs34811474 | 4 | 25408838 |
| 10 | rs13135092 | 4 | 103198082 |
| 11 | rs13136239 | 4 | 140908755 |
| 12 | rs6869446 | 5 | 65570607 |
| 13 | rs1976423 | 5 | 104042643 |
| 14 | rs17474406 | 5 | 122732342 |
| 15 | rs1946247 | 5 | 160836620 |
| 16 | rs11751591 | 6 | 33794215 |
| 17 | rs6907508 | 6 | 34592090 |
| 18 | rs6926377 | 6 | 145105354 |
| 19 | rs10259354 | 7 | 3487414 |
| 20 | rs7798894 | 7 | 21552995 |
| 21 | rs6966540 | 7 | 95727967 |
| 22 | rs12537376 | 7 | 114025053 |
| 23 | rs11786084 | 8 | 142651709 |
| 24 | rs10992729 | 9 | 96181075 |
| 25 | rs6478241 | 9 | 119252629 |
| 26 | 9:140251458_G_A | 9 | 140251458 |
| 27 | rs2183271 | 10 | 21957229 |
| 28 | rs11599236 | 10 | 106454672 |
| 29 | rs12765185 | 10 | 134977077 |
| 30 | rs61883178 | 11 | 16317779 |
| 31 | rs1443914 | 13 | 53917230 |
| 32 | rs12435797 | 14 | 73797669 |
| 33 | rs2006281 | 14 | 104327732 |
| 34 | rs2386584 | 15 | 91539572 |
| 35 | rs285026 | 16 | 77100089 |
| 36 | rs11871043 | 17 | 43172849 |
| 37 | rs11079993 | 17 | 50301552 |
| 38 | rs62098013 | 18 | 50863861 |
| 39 | rs2424248 | 20 | 19650324 |

Genomic risk loci (as determined by FUMA). rsID = rsID of locus lead SNP, Chr = chromosome, p = GWAS p value, pos = position (base-pairs).

**Table 4:**
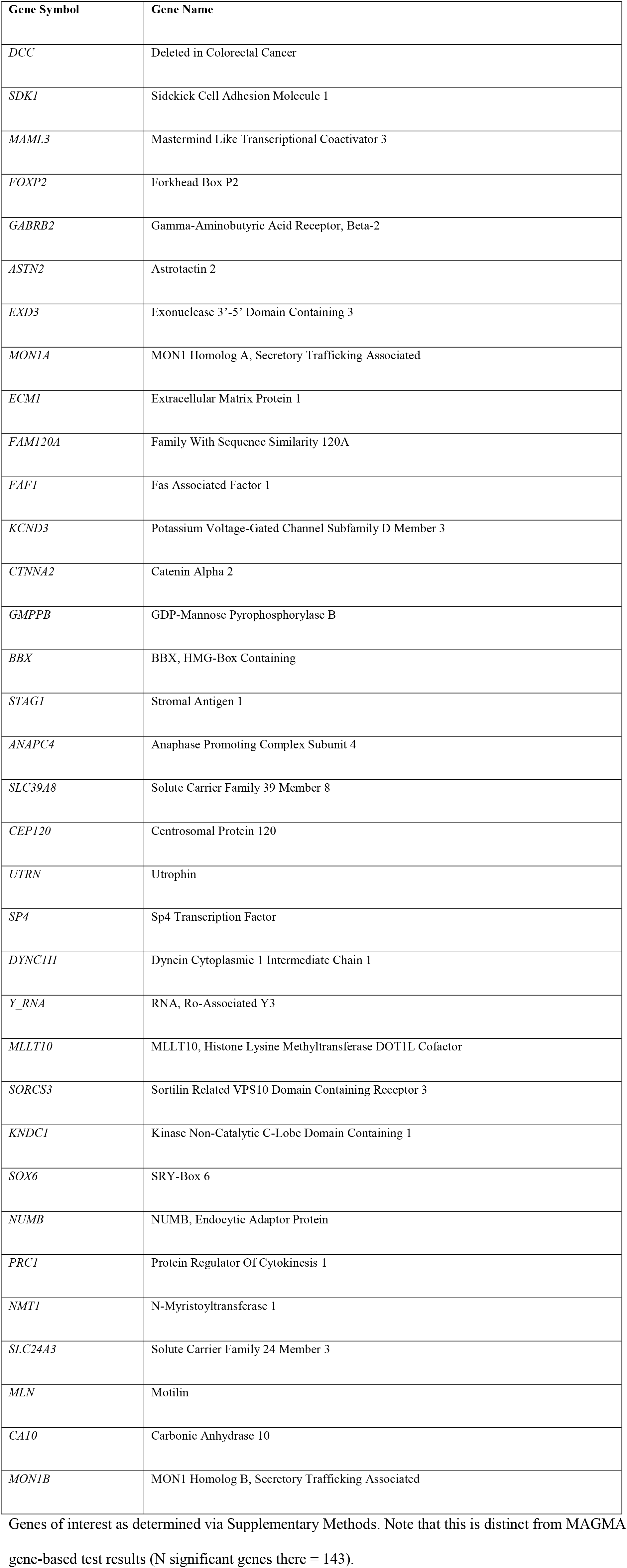
Genes of Interest.

**Table 5:** Genetic Correlations.

| Trait | rg | se | z | h2 | P<sub>h2</sub> (fdr) | source | PMID | Category | p | P (fdr-corrected) |
| --- | --- | --- | --- | --- | --- | --- | --- | --- | --- | --- |
| MDD | 0.53 | 0.03 | 18.92 | 0.077 | 1.25E-47 | PGC | 29700475 | psychiatric | 7.68E-80 | 1.69E-78 |
| Depressive symptoms | 0.59 | 0.03 | 17.16 | 0.047 | 6.87E-29 | ld_hub | 27089181 | psychiatric | 5.63E-66 | 6.19E-65 |
| BMI | 0.31 | 0.02 | 15.69 | 0.138 | 5.42E-59 | GIANT consortium | 25673413 | anthropometric | 1.90E-55 | 1.39E-54 |
| Neuroticism | 0.4 | 0.03 | 11.9 | 0.089 | 3.66E-26 | ld_hub | 27089181 | personality | 1.24E-32 | 6.82E-32 |
| Subjective well being | -0.36 | 0.04 | -8.94 | 0.025 | 2.77E-32 | ld_hub | 27089181 | psychiatric | 3.78E-19 | 1.66E-18 |
| Low Relative Amplitude | -0.3 | 0.05 | -6.37 | 0.053 | 3.03E-13 | In-house analysis | 30120083 | circadian | 1.91E-10 | 7.00E-10 |
| Rheumatoid Arthritis | 0.16 | 0.03 | 4.7 | 0.160 | 7.41E-08 | ld_hub | 24390342 | autoimmune | 2.64E-06 | 8.30E-06 |
| Anxiety (Case-Control) | 0.49 | 0.11 | 4.53 | 0.081 | 0.00405 | PGC | 26754954 | psychiatric | 5.91E-06 | 1.63E-05 |
| Schizophrenia | 0.1 | 0.03 | 4.08 | 0.443 | 6.56E-79 | PGC | 25056061 | psychiatric | 4.50E-05 | 1.10E-04 |
| Asthma | 0.22 | 0.06 | 3.63 | 0.123 | 3.53E-06 | ld_hub | 17611496 | autoimmune | 3.00E-04 | 6.60E-04 |
| PGC cross-disorder analysis | 0.13 | 0.04 | 3.54 | 0.172 | 7.89E-36 | ld_hub | 23453885 | psychiatric | 4.00E-04 | 8.00E-04 |
| PTSD (European Ancestry) | 0.41 | 0.12 | 3.28 | 0.097 | 0.030855 | PGC | 28439101 | psychiatric | 0.001047 | 1.92E-03 |
| Autism spectrum disorder | -0.1 | 0.04 | -2.22 | 0.451 | 9.38E-17 | ld_hub | NA | psychiatric | 0.026 | 0.0443 |
| Primary biliary cirrhosis | 0.1 | 0.04 | 2.17 | 0.376 | 1.11E-08 | ld_hub | 26394269 | autoimmune | 0.03 | 0.047 |
| Anorexia Nervosa | -0.06 | 0.03 | -2.14 | 0.556 | 2.18E-63 | ld_hub | 24514567 | psychiatric | 0.032 | 0.0471 |
| Inflammatory Bowel Disease (European | 0.05 | 0.03 | 1.75 | 0.333 | 9.17E-21 | ld_hub | 26192919 | autoimmune | 0.08 | 0.1101 |
| Ancestry) |  |  |  |  |  |  |  |  |  |  |
| Celiac disease | -0.07 | 0.05 | -1.49 | 0.314 | 2.50E-10 | ld_hub | 20190752 | autoimmune | 0.136 | 0.1756 |
| Crohn's disease | 0.04 | 0.03 | 1.35 | 0.504 | 2.65E-17 | ld_hub | 26192919 | autoimmune | 0.179 | 0.2125 |
| Systemic lupus erythematosus | 0.06 | 0.04 | 1.33 | 0.390 | 9.77E-09 | ld_hub | 26502338 | autoimmune | 0.184 | 0.2125 |
| Ulcerative colitis | 0.04 | 0.04 | 1.08 | 0.257 | 1.19E-14 | ld_hub | 26192919 | autoimmune | 0.281 | 0.3094 |
| Bipolar disorder | -0.02 | 0.04 | -0.66 | 0.436 | 5.51E-29 | ld_hub | 21926972 | psychiatric | 0.509 | 0.5329 |
| Parkinson's disease | 0 | 0.04 | 0.05 | 0.409 | 0.000761 | ld_hub | 19915575 | neurological | 0.961 | 0.9612 |

rg = genetic correlation coefficient value, se = standard error of correlation value, z = z value, h2 =
SNP-heritability value, ph2(fdr) = p value (FDR-corrected) for SNP-heritability, source = source of
GWAS summary statistics, PMID = PubMed ID of associated paper (if applicable), p = p value for
genetic correlation coefficient, p(fdr) = FDR-corrected p value for genetic correlation coefficient.

#### Genetic Correlations

MDD was the psychiatric phenotype most significantly correlated with MCP (r_g_ = 0.53, p_FDR_ = 1.69e-78) and the highest significant correlation coefficient value was for MCP and depressive symptoms (r_g_ = 0.59, p_FDR_ = 6.19e-65). MCP was also positively correlated with neuroticism (r_g_ = 0.40), anxiety (r_g_ = 0.49), schizophrenia (r_g_ = 0.10), cross-disorder psychiatric phenotype (r_g_ = 0.13) and PTSD (r_g_ = 0.41). Significant negative correlations were observed between MCP and subjective well-being (r_g_ = −0.36), ASD (r_g_ = −0.10) and with AN (r_g_ = −0.06). BD was not significantly correlated with MCP (P_FDR_ > 0.05). Rheumatoid arthritis (r_g_ = 0.16) and asthma (r_g_ = 0.22) were significantly positively correlated with MCP, as was primary biliary cholangitis (r_g_ = 0.10). The autoimmune gastrointestinal disorders Ulcerative Colitis and Crohn’s disease were not correlated with MCP (P_FDR_ > 0.05). SLE was not correlated with MCP (P_FDR_ > 0.05). BMI showed significant positive correlation with MCP (r_g_ = 0.31). MCP was negatively correlated with low relative amplitude (r_g_ = −0.30). There was no correlation between Parkinson’s disease and MCP (P_FDR_ > 0.05).

#### Mendelian Randomisation of MCP and Major Depressive Disorder

QQ plots, leave-one out versus t-value plots (Supplementary Figure 1) and Anderson-Darling/Shapiro-Wilk test p values indicated that models without dispersion were best-fitting (Table 6 rows 1-3, p_AD_ > 0.05, p_SW_ > 0.05). Effects of outliers (idiosyncratic pleiotropy) are not ameliorated in models with dispersion despite robust regression (Supplementary Fig 1: D, E, F right-hand panels). The model allowing the greatest amelioration of pleiotropy is one without over-dispersion and with a Tukey loss function (Table 6: row 3, Fig 7: C). This indicates idiosyncratic pleiotropy (pleiotropy in some but not all instruments), i.e. that a subset of instruments may affect MCP through pathways other than via MDD (the exposure). The causal effect of MDD on MCP is positive and significant at beta = 0.019 and p = 0.0006.

**Table 6:** MDD Exposure MR-RAPS Results.

| overdispersion | Loss function | $\beta$ | SE ($\beta$) | P ($\beta$) | P (AD) | P (SW) | $\tau$ | P ($\tau$) | C.F |
| --- | --- | --- | --- | --- | --- | --- | --- | --- | --- |
| FALSE | L2 | 0.0117 | 0.0052 | 0.0241 | 0.9375 | 5.34E-01 | NA | NA | Fig 7. A |
| FALSE | Huber | 0.0153 | 0.0054 | 0.0042 | 0.9285 | 5.23E-01 | NA | NA | Fig 7. B |
| **FALSE** | **Tukey** | **0.0185** | **0.0054** | **0.0006** | **0.9230** | **5.18E-01** | **NA** | **NA** | Fig 7. C |
| TRUE | L2 | -0.0096 | 0.0132 | 0.4671 | 0.0080 | 1.76E-03 | 1.61E-04 | 0.0470 | Fig 7. D |
| TRUE | Huber | -0.0056 | 0.0126 | 0.6556 | 0.0087 | 2.11E-03 | 1.30E-04 | 0.0677 | Fig 7. E |
| TRUE | Tukey | -0.0065 | 0.0137 | 0.6330 | 0.0055 | 9.03E-04 | 1.67E-04 | 0.0627 | Fig 7. F |

MR results for MDD-exposure. B refers to the causal effect, SE ( $\beta$ ) and P ( $\beta$ ) to the standard error and p value of $\beta$ , P (AD) to the Anderson-Darling test of normality p value, P (SW) to the Shapiro-Wilk test of normality p value, tau to the over-dispersion statistic size and P ( $\tau$ ) to the p value. C.F = corresponding QQ plot panel for the model. P ( $\tau$ ) was calculated from the tau estimate and its standard error, according to methods outlined by Altman & Bland (2011). The row of the table corresponding to the regression model found to be best-fitting is in bold.

Models with dispersion are a better fit than those without (Supplementary Fig 2: A, B, C vs D, E, F, Table 7: rows 4-6, p_AD_ > 0.05, p_SW_ > 0.05, pτ << 0.05). This indicates that effectively all instruments are pleiotropic (affecting MDD through pathways other than via MCP). The causal effect of MCP on MDD is positive and significant at beta = 0.16 and p = 0.047.

**Table 7:** MCP Exposure MR-RAPS Results.

| overdispersion | Loss function | $\beta$ | SE ($\beta$) | P ($\beta$) | P (AD) | P (SW) | $\tau$ | P ($\tau$) | C.F |
| --- | --- | --- | --- | --- | --- | --- | --- | --- | --- |
| FALSE | L2 | 0.1714 | 0.0605 | 0.0046 | 0.4256 | 0.0853 | NA | NA | Fig 8. A |
| FALSE | Huber | 0.1815 | 0.0621 | 0.0034 | 0.4247 | 0.0835 | NA | NA | Fig 8. B |
| FALSE | Tukey | 0.2097 | 0.0621 | 0.0007 | 0.4221 | 0.0784 | NA | NA | Fig 8. C |
| TRUE | L2 | 0.1201 | 0.0790 | 0.1286 | 0.8374 | 0.2853 | 9.81E-05 | 2.43E-03 | Fig 8. D |
| TRUE | Huber | 0.1446 | 0.0801 | 0.0712 | 0.8289 | 0.2724 | 9.18E-05 | 5.13E-03 | Fig 8. E |
| **TRUE** | **Tukey** | **0.1578** | **0.0795** | **0.0471** | **0.8236** | **0.2641** | **8.77E-05** | **7.09E-03** | Fig 8. F |

MR results for chronic pain-exposure. B refers to the causal effect, SE ( $\beta$ ) and P ( $\beta$ ) to the standard error and p value of $\beta$ , P (AD) to the Anderson-Darling test of normality p value, P (SW) to the Shapiro-Wilk test of normality p value, tau to the over-dispersion statistic size and P ( $\tau$ ) to the p value. P ( $\tau$ ) was calculated from the tau estimate and its standard error, according to methods outlined by Altman & Bland (2011). The row of the table corresponding to the regression model found to be of best fit is in bold.

Overall, this analysis suggests a bi-directional causal relationship between MCP and MDD, with the causal effect more significant in the direction MDD 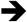 MCP, and with a smaller subset of (horizontally) pleiotropic instruments compared to when MCP is the exposure.

#### PRS Prediction of Chronic Widespread Pain

Increasing MCP PRS value was significantly associated with having chronic pain all over the body (Table 8: p = 1.45e-109), with each per-SD increase in PRS associated with a 63% increase in the odds of having chronic widespread pain.

**Table 8:** PRS Results.

| **Term** | **Estimate** | **SE (Estimate)** | **Z** | **P** | **OR** |
| --- | --- | --- | --- | --- | --- |
| (Intercept) | -61.418 | 2.763 | -22.227 | 1.90E-109 | 2.12E-27 |
| Age | 0.016 | 0.002 | 7.451 | 9.25E-14 | 1.02 |
| Sex | -0.488 | 0.035 | -14.073 | 5.56E-45 | 0.61 |
| PRS | 0.488 | 0.022 | 22.239 | 1.45E-109 | 1.63 |

Regression beta coefficient values (Estimate), odds ratios (OR), and P values. The reference level for 'sex' is set to female, PRS = z-polygenic risk score.

## Discussion

We identified 76 independent genome-wide significant SNPs across 39 loci associated with multisite chronic pain (MCP). The genes of interest had diverse functions, but many were implicated in nervous-system development, neural connectivity and neurogenesis.

### Genes of interest identified in GWAS of MCP

Potentially interesting genes included *DCC* (Deleted in Colorectal Cancer a.k.a. DCC netrin 1 receptor) which encodes DCC, the receptor for the guidance cue netrin 1, which is important for nervous-system development (Manitt *et al.*, 2011). *SDK1* (Sidekick Cell Adhesion molecule 1) is implicated in HIV-related nephropathy in humans (Kaufman *et al.*, 2007) and synaptic connectivity in vertebrates (Yamagata and Sanes, 2008), and *ASTN2* (Astrotactin 2) is involved in glial-guided neuronal migration during development of cortical mammalian brain regions (Wilson *et al.*, 2010).

*MAML3* (Mastermind-Like Transcriptional coactivator 3) is a key component of the Notch signalling pathway (Andersson, Sandberg and Lendahl, 2011; Kitagawa, 2015), which regulates development and maintenance of a range of cell and tissue types in metazoans. During neurogenesis in development the inhibition of Notch signalling by Numb promotes neural differentiation (Ables *et al.*, 2011). Numb is encoded by *NUMB* (Endocytic Adaptor Protein), which was also associated with MCP. In the adult brain Notch signalling has been implicated in CNS plasticity across the lifespan (Ables *et al.*, 2011).

*CTNNA2* (Catenin Alpha 2) encodes a protein involved in cell-cell adhesion (Janssens *et al.*, 2001), found to play a role in synapse morphogenesis and plasticity (Arikkath and Reichardt, 2008; Arikkath *et al.*, 2009). *CEP120* (Centrosomal Protein 120) encodes Cep120, vital for Interkinetic Nuclear Migration (INM) in neural progenitor cells of the cortex (Guerrier and Polleux, 2007). *KNDC1* (Kinase Non-Catalytic C-Lobe Domain Containing 1) encodes v-KIND in mice, linked to neural morphogenesis in the cortex (Hayashi *et al.*, 2017), and KNDC1 in humans, linked to neuronal dendrite development and cell senescence (Ji *et al.*, 2018). *SOX6* (SRY-Box 6) is part of the *Sox* gene family, first characterised in mouse and human testis-determining gene *Sry* (Sinclair *et al.*, 1990) and encoding transcription factors involved in a range of developmental processes (Denny *et al.*, 1992; Cohen-Baraka *et al.*, 2001). *SOX6* may be involved in development of skeletal muscle (Cohen-Baraka *et al.*, 2001), maintenance of brain neural stem cells (Kurtsdotter *et al.*, 2017) and cortical interneuron development (Batista-Brito *et al.*, 2009), and variants in this gene have been associated with bone mineral density in both white and Chinese populations (Yang *et al.*, 2012). *CA10* (Carbonic Anhydrase 10) is predominantly expressed in the CNS, encoding a protein involved in development and maintenance of synapses (Sterky *et al.*, 2017). *DYNC1I1* (Dynein Cytoplasmic 1 Intermediate Chain 1) encodes a subunit of cytoplasmic dynein, a motor protein which plays a role in cargo transport along microtubules, including in the function of neuronal cells (Goldstein and Yang, 2000). *UTRN* (Utrophin) is a homologue of Duchenne Muscular Dystrophy gene (*DMD*), encoding utrophin protein which is localised to the neuromuscular junction (NMJ) (Blake, Tinsley and Davies, 1996). Utrophin has also been implicated in neutrophil activation (Cerecedo *et al.*, 2010), dystrophin-associated-protein (DPC)-like complex formation in the brain (Blake *et al.*, 1999), and is expressed during early foetal brain development in neurons and astrocytes (Sogos *et al.*, 2002).

*FOXP2* encodes a member of the FOX family of transcription factors, which are thought to regulate expression of hundreds of genes in both adult and foetal tissue, including the brain. These transcription factors may play an important role in brain development, neurogenesis, signal transmission and synaptic plasticity (Vernes *et al.*, 2011). *FOXP2* is essential for normal speech and language development (MacDermot *et al.*, 2005). *GABRB2* encodes the beta subunit of the GABA (gamma-aminobutyric acid) A receptor, a multi-subunit chloride channel which mediates fast inhibitory synaptic transmission in the CNS (Jacob, Moss and Jurd, 2008). The protein also acts as a histamine receptor, mediating cellular response to histamines (Saras *et al.*, 2008).

Another group of genes associated with MCP were linked to cell-cycle progression, DNA replication and apoptosis such as *EXD3* (Exonuclease 3’-5’ Domain Containing 3), which encodes a protein involved in maintaining DNA fidelity during replication (‘proof-reading’) (Bębenek and Ziuzia-Graczyk, 2018). *BBX* (HMG-Box Containing protein 2) encodes an HMG (high mobility group) box-containing protein necessary for cell-cycle progression from G1 to S phase (Malarkey and Churchill, 2012). *STAG1* (Cohesin Subunit SA-1) encodes a cohesin-complex component – cohesin ensures sister chromatids are organised together until prometaphase (Losada *et al.*, 2000; Peters and Nishiyama, 2012; Murayama and Uhlmann, 2014). *ANAPC4* (Anaphase Promoting Complex Subunit 4) encodes a protein making up the anaphase promoting complex (APC), an essential ubiquitin ligase for eukaryotic cell-cycle progression (Peters, 2006). *PRC1* (Protein Regulator of Cytokinesis 1) is involved in the regulation of cytokinesis (Shrestha *et al.*, 2012), the final stage of the cell cycle. *Y RNA* (Small Non-Coding RNA, Ro-Associated Y3) encodes a small non-coding Y RNA. These RNAs have been implicated in a wide range of processes, including cell stress response, DNA replication initiation and RNA stability (Kowalski and Krude, 2015). *FAM120A* (Oxidative Stress-Associated Src Activator) encodes an RNA-binding protein which regulated Src-kinase activity during oxidative stress-induced apoptosis (Tanaka *et al.*, 2009). The protein encoded by *MON1B* (MON1 Homolog B, Secretory Trafficking Associated) is necessary for clearance of cell ‘corpses’ following apoptosis, with defects associated with autoimmune pathology (Kinchen and Ravichandran, 2010). *FAF1* (Fas Associated Factor 1) encodes a protein which binds the Fas antigen to initiate or facilitate apoptosis, amongst a wide range of other biological processes (including neuronal cell survival) (Menges, Altomare and Testa, 2009).

Several MCP associated genes have been previously implicated in diseases such as Brugada Syndrome 9 and Spinal ataxia 19 & 22 (*KCND3*) (Giudicessi *et al.*, 2011; Duarri *et al.*, 2012; Lee *et al.*, 2012), Systemic lupus erythematosus (SLE) (Y RNAs) (Kowalski and Krude, 2015), Joubert syndrome 31 and short-rib thoracic dysplasia 13 (*CEP120*) (Roosing *et al.*, 2016), Amyotrophic lateral sclerosis (ALS) (*FAF1*) (Baron *et al.*, 2014), Urbach-Wiethe disease (*ECM1*) (Hamada *et al.*, 2003; Oyama *et al.*, 2003), mental retardation and other cohesinopathies such as Cornelia de Lange Syndrome (*STAG1*) (Liu and Krantz, 2009; Lehalle *et al.*, 2017), split hand/split foot malformation (*DYNC1I1*) (Roberts *et al.*, 1991; Tayebi *et al.*, 2014), and a wide range of cancers (*PRC1*) (Li *et al.*, 2018). Other disorders found to involve MCP-related genes include schizophrenia (*FOXP2* and *GABRB2*) (Petryshen *et al.*, 2005; Sanjuá *et al.*, 2006; Lo *et al.*, 2007; Laroche *et al.*, 2008; Tolosa *et al.*, 2010; Li *et al.*, 2013; Yin *et al.*, 2018), intellectual disability and epilepsy (*GABRB2*) (Srivastava *et al.*, 2014), and neuroleptic-induced tardive dyskinesia (*GABRB2*) (Inada *et al.*, 2008).

Overall, this indicated that MCP, a chronic pain phenotype, involves structural and functional changes to the brain, including impact upon neurogenesis and synaptic plasticity both during development and in adulthood. Also implicated was regulation of cell-cycle progression and apoptosis. There was also evidence of pleiotropy, with genes associated with a range of neurodegenerative, psychiatric, developmental and autoimmune disease traits, as well as being associated with MCP.

### Genetic correlations

Chronic pain and chronic pain disorders are often comorbid with psychiatric and neurodevelopmental disorders (Gureje *et al.*, 2008). This has been observed for Major Depressive Disorder (MDD) (Nicholl *et al.*, 2014; McIntosh *et al.*, 2016), post-traumatic stress-disorder (PTSD) (Shipherd *et al.*, 2007; Dunn *et al.*, 2011; Phifer *et al.*, 2011; Outcalt *et al.*, 2015; Akhtar *et al.*, 2018), schizophrenia (Watson, Chandarana and Merskey, 1981; de Almeida *et al.*, 2013; Engels *et al.*, 2014) and bipolar disorder (BD) (Nicholl *et al.*, 2014; Stubbs *et al.*, 2015). There are also reported differences in the perception of pain and interoception (sensing and integration of bodily signals) for people with schizophrenia (Lévesque *et al.*, 2012; Urban-Kowalczyk, Pigońska and Śmigielski, 2015), anorexia nervosa (AN) (Strigo *et al.*, 2013; Bär *et al.*, 2015; Bischoff-Grethe *et al.*, 2018) and autism spectrum disorders (ASD) (Clarke, 2015; Gu *et al.*, 2018), with some evidence of an increase in pain thresholds for AN and ASD.

There is significant cross-talk between the immune system and nervous system in nociception and sensitisation leading to chronic pain (Pinho-Ribeiro, Verri and Chiu, 2017; Kwiatkowski and Mika, 2018), and many autoimmune disorders cause or have been associated with chronic pain including neuroinflammation implicated in development of neuropathic pain (Ren and Dubner, 2010).

Similarly, obesity and chronic pain are often comorbid, with lifestyle factors such as MDD and sleep disturbance also impacting on chronic pain (Okifuji and Hare, 2015; Paley and Johnson, 2016).

Obesity and related chronic inflammation may affect chronic pain (Ramesh, Maclean and Philipp, 2013), and adipose tissue is metabolically active in ways that can affect pain perception and inflammation (Hotamisligil GS, Shargill NS, 1993; Olefsky and Glass, 2010; Chawla, Nguyen and Goh, 2011).

Sleep changes and loss of circadian rhythm is common in those with chronic pain (Alföldi, Wiklund and Gerdle, 2014), and myriad chronic diseases, including chronic pain, have shown diurnal patterns in symptom severity, intensity and mortality (Smolensky *et al.*, 2015; Segal *et al.*, 2018). Chronic pain is also a common component of many neurological diseases, particularly Parkinson’s disease (Borsook, 2012), and disorders such as Multiple Sclerosis and migraines are considered neurological in nature.

MCP showed moderate positive genetic correlation with a range of psychiatric disorders including MDD, SCZ, and PTSD, along with traits anxiety and neuroticism. The magnitude of genetic correlation between MCP and MDD was similar to that shown for von Korff chronic pain grade (a chronic pain phenotype) and MDD by McIntosh et al via a mixed-modelling approach (ρ = 0.53) (McIntosh *et al.*, 2016). This is in line with previous observations of association and indicates that shared genetic risk factors exist between MCP and a range of psychiatric disorders, most notably MDD, and that the genetic correlation between MCP and MDD matches with that between MDD and von Korff CPG, a validated chronic-pain questionnaire-derived phenotype (Von Korff *et al.*, 1992).

Autoimmune disorders rheumatoid arthritis, asthma and primary biliary cholangitis showed positive genetic correlation with MCP. However, gastrointestinal autoimmune disorders UC, IBD and Crohn’s Disease did not. This suggests separate genetic variation and mechanisms underlying chronic pain associated with these autoimmune disorders compared to those outwith the digestive system. Pain related to inflammatory bowel diseases may represent something less ‘chronic’ and more ‘on-going acute’, as stricture, abscesses and partial or complete obstruction of the small bowel result in pain (Docherty, Jones and Wallace, 2011). Structural and functional brain changes associated with the transition to chronic pain may also play a less central role in gastrointestinal autoimmune disorder-associated pain, due to potential for the enteric nervous system (ENS) to act independently from the CNS, and the role of the gut-brain axis (GBA) (Cryan and Dinan, 2012; Carabotti *et al.*, 2015).

There was significant negative genetic correlation between low relative amplitude, a circadian rhythmicity phenotype indicating poor rhythmicity (Ferguson *et al.*, 2018). Opposing direction of effect of genetic variants on MCP versus low RA may mean that insomnia and other sleep difficulties (for which low RA represents a proxy phenotype) associated with MCP are due to environmental and lifestyle factors related to chronic pain, rather than shared genetic factors predisposing to increased risk for both traits. There was also significant negative genetic correlation between MCP and both AN and ASD, which may be linked to changes in interoception and atypical pain experience seen in individuals with these conditions (Strigo *et al.*, 2013; Bär *et al.*, 2015; Clarke, 2015; Bischoff-Grethe *et al.*, 2018; Gu *et al.*, 2018), and may suggest a genetic basis for increased pain thresholds.

### SNP heritability of MCP

LDSR analyses gave a heritability estimate of 10.2% for MCP, lower than the pseudo-h^2^ estimate of 10.3% given by BOLT-LMM. this suggests SNP-heritability (h^2^) of MCP to be roughly-10%, slightly lower than an estimate of ‘any chronic pain’ of 16%, and markedly lower than a heritability estimate of 30% for ‘severe chronic pain’ derived from a pedigree-based analyses (Hocking *et al.*, 2012).

### Causal associations between MDD and MCP

Mendelian randomisation analyses indicated a bi-directional causal relationship between MDD and MCP, with widespread pleiotropy and a less significant causal estimate value for MCP as the exposure – this suggests most instruments for MCP are pleiotropic, affecting MDD through pathways other than directly through MCP. In contrast, only a small subset of instruments for MDD as the exposure were found to be pleiotropic.

In both cases the causal estimate values tend to be small (< 0.2), emphasising contributions of the environment, lifestyle, and other conditions and disorders to development of both MDD and chronic pain.

### Relationship between MCP and CWP

**I**t has been argued that CWP, and other clinical syndromes involving chronic pain all over the body, represent the upper end of a spectrum of centralisation of pain, or the extreme of a chronic pain state (Phillips and Clauw, 2011). It has also been argued that there are not “natural cut-off points” when it comes to chronic widespread pain versus localised chronic pain (Kamaleri *et al.*, 2008). MCP PRS was significantly associated with increased odds of having chronic pain all over the body/ CWP, suggesting that chronic widespread pain represents the upper end of a spectrum ‘widespreadness’ of chronic pain, as previously suggested (Kamaleri *et al.*, 2008; Phillips and Clauw, 2011), and in the least that shared genetic variants predict both MCP and CWP.

## Conclusions

Multisite chronic pain (MCP), a chronic pain phenotype derived from the number of sites at which chronic pain is experienced, is a complex trait with moderate heritability. To date, this study represents the largest GWAS of any chronic pain phenotype and elucidates potential underlying mechanisms of chronic pain development. Genetic correlations with a range of psychiatric, personality, autoimmune, anthropometric and circadian traits were identified.

The genes potentially associated with MCP implicated neurogenesis, neuronal development and neural connectivity, along with cell-cycle and apoptotic processes, and expression was primarily within brain tissues, specifically the cerebellum and cerebellar cortex. This is in line with theories of functional and structural changes to the brain contributing to development of chronic pain (Apkarian, Hashmi and Baliki, 2011; Baliki *et al.*, 2012, 2014; Baliki and Apkarian, 2015; Fasick *et al.*, 2015), and may also explain genetic correlations between a range of psychiatric and neurodevelopmental phenotypes and MCP.

Finally, a bi-directional causal relationship was identified between MDD and MCP, underlining the importance of studying psychiatric phenotypes within the context of chronic pain research and *vice versa*.

## Acknowledgements

We thank all participants in the UK Biobank study. UK Biobank was established by the Wellcome Trust, Medical Research Council, Department of Health, Scottish Government and Northwest Regional Development Agency. UK Biobank has also had funding from the Welsh Assembly Government and the British Heart Foundation. Data collection was funded by UK Biobank.

RJS is supported by a UKRI Innovation-HDR-UK Fellowship (MR/S003061/1). JW is supported by the JMAS Sim Fellowship for depression research from the Royal College of Physicians of Edinburgh (173558). AF is supported by an MRC Doctoral Training Programme Studentship at the University of Glasgow (MR/K501335/1). KJAJ is supported by an MRC Doctoral Training Programme Studentship at the Universities of Glasgow and Edinburgh. DJS acknowledges the support of the Brain and Behavior Research Foundation (Independent Investigator Award 1930), a Lister Prize Fellowship (173096) and the MRC Mental Health Data Pathfinder Award (MC_PC_17217).

