## Supplementary Information for "Genome-wide Association Study of Multisite Chronic Pain in UK Biobank"

### Defining Genes of Interest

The top 10 most-significant FUMA MAGMA gene-based test results (ranked by Bonferroni-corrected p value for the gene-based test) were taken.

In addition, genomic rick loci (GenomicRiskLoci.txt), and ANNOVAR annotations (annov.txt) were taken from the FUMA SNP2GENE output and matched by 'uniqID' to make a subset of data consisting of ANNOVAR annotations for lead SNPs at genomic risk loci. N = 46 genes (39 risk loci, 1 locus has 2 lead SNPs. Lead SNPs can have multiple ANNOVAR gene annotations).

Genes that were already in the MAGMA gene-based-test list were removed (-7), as were RNAs and pseudogenes that were not well-characterised or associated with diseases or traits from preliminary OMIM (Online Mendelian Inheritance in Man), GeneCards and PubMed 'Gene' database searches.

This subset was combined with the top 10 FUMA MAGMA gene-based test results to give N = 35 genes.

In summary, this gene set consists of the top 10 most-significant MCP-associated genes from the FUMA MAGMA gene-based test results, along with genes associated with the 39 genomic risk loci for MCP.

### MR-RAPS

Briefly, the main problems associated with pleiotropy in MR analyses are that instruments may be invalid due to pleiotropy, associated measurement error, weak-instrument bias, and selection bias. MR-RAPS treats pleiotropy-related issues as an 'errors in regression' problem (Zhao *et al.*, 2018), in contrast to MR modifications with their roots in meta-analyses (such as Egger and Inverse-Variance-Weighted (IVW)) which treat pleiotropy in a manner similar to heterogeneity between individual studies in a meta-analysis (Smith and Hemani, 2014; Zheng, Baird, *et al.*, 2017).

The basic model types fitted by MR-RAPS are: no pleiotropy, systematic pleiotropy (all instruments subject to pleiotropy), and a combination of idiosyncratic (only some instruments subject to pleiotropy) and systematic pleiotropy. MR-RAPS was used to fit a total of 6 regressions per analysis (Table 2), and diagnostics for the fit to each model were evaluated to select the optimal model type prior to interpretation of the causal effect estimate.

[Supplementary Table 1: MR-RAPS Models]

[Supplementary Figure 1 : MDD Exposure QQ Plots]

[Supplementary Figure 2 : MCP Exposure QQ Plots]

LocusZoom Plots

Plots were created using locuszoom v1.4 standalone (Pruim *et al*., 2010) with pop flag set to EUR, build set to hg19 and source set to 1000G_Nov2014. Associated regions were defined as SNP with an r2 > 0.1 in a 500mb radius of the lead SNP. Plot boundaries were defined as +/- 1 MB outwith the associated region. See Supplementary Figures 3-49.
