## Supplementary Figure 3 for "Genome-wide Association Study of Multisite Chronic Pain in UK Biobank"

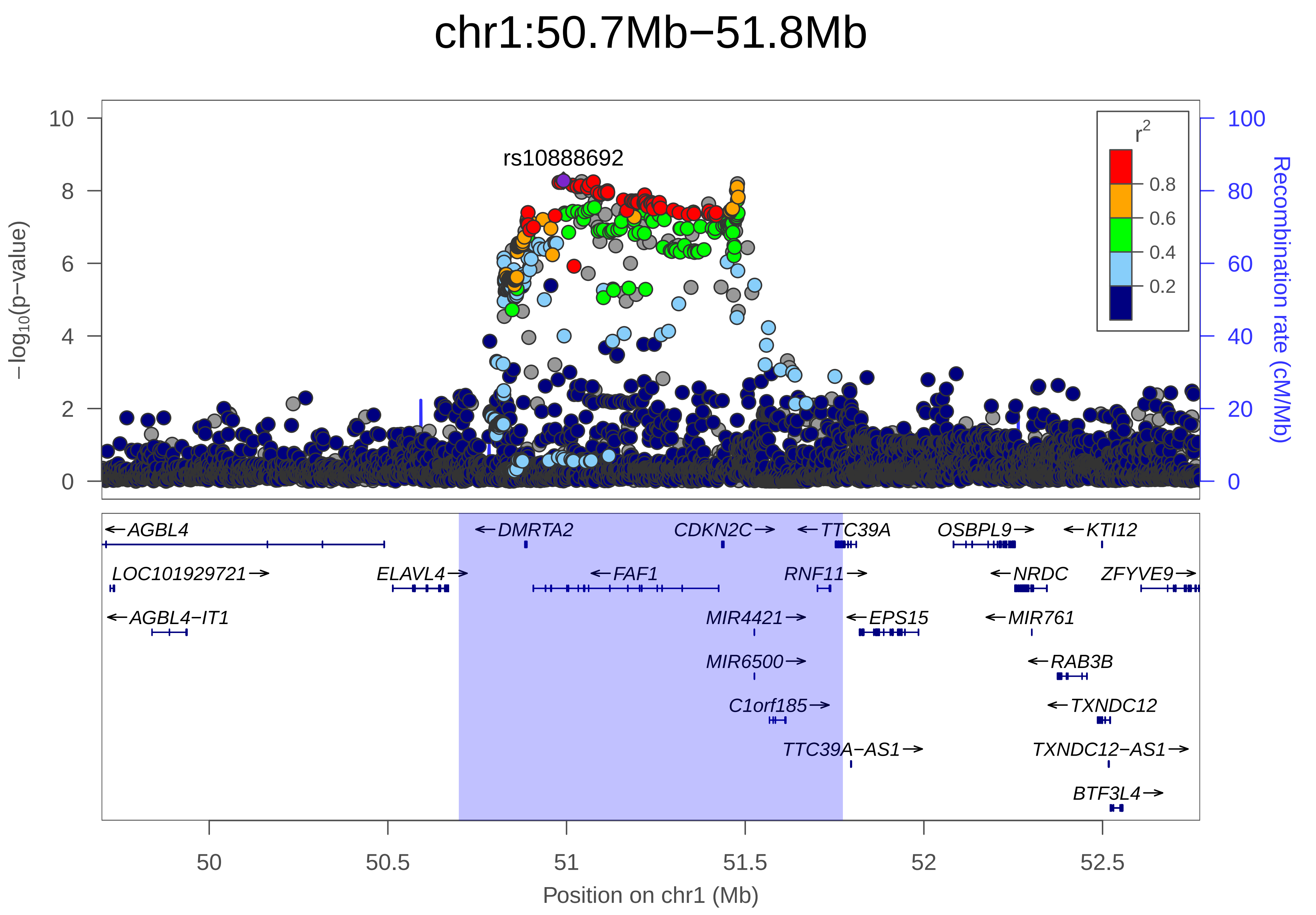

chr1:50.7Mb−51.8Mb
rs10888692
−log10(p−value)
Recombination rate (cM/Mb)
r2
0.8
0.6
0.4
0.2
10
8
6
4
2
0
100
80
60
40
20
0
AGBL4
DMRTA2
CDKN2C
TTC39A
OSBPL9
KTI12
LOC101929721
ELAVL4
FAF1
RNF11
NRDC
ZFYVE9
AGBL4−IT1
MIR4421
EPS15
MIR761
MIR6500
RAB3B
C1orf185
TXNDC12
TTC39A−AS1
TXNDC12−AS1
BTF3L4
50
50.5
51
51.5
52
52.5
Position on chr1 (Mb)

| | |
|---|---|
| date: | Mon Dec 10 07:08:39 2018 |
| build: | hg19 |
| display range: | chr1:49698760−52773078 [49698760−52773078] |
| hilite range: | 50.699Mb − 51.773Mb [ 50699000 − 51773000 ] |
| reference SNP: | chr1:50991473 |
| number of SNPs plotted: | 6051 |
| min P−value: | 5.3E−9 [chr1:50991473] |
| max P−value: | 1E0 [chr1:49794844] |

Make more plots at http://csg.sph.umich.edu/locuszoom/
