## Supplementary Figure 4 for "Genome-wide Association Study of Multisite Chronic Pain in UK Biobank"

### chr1:112.1Mb−112.4Mb

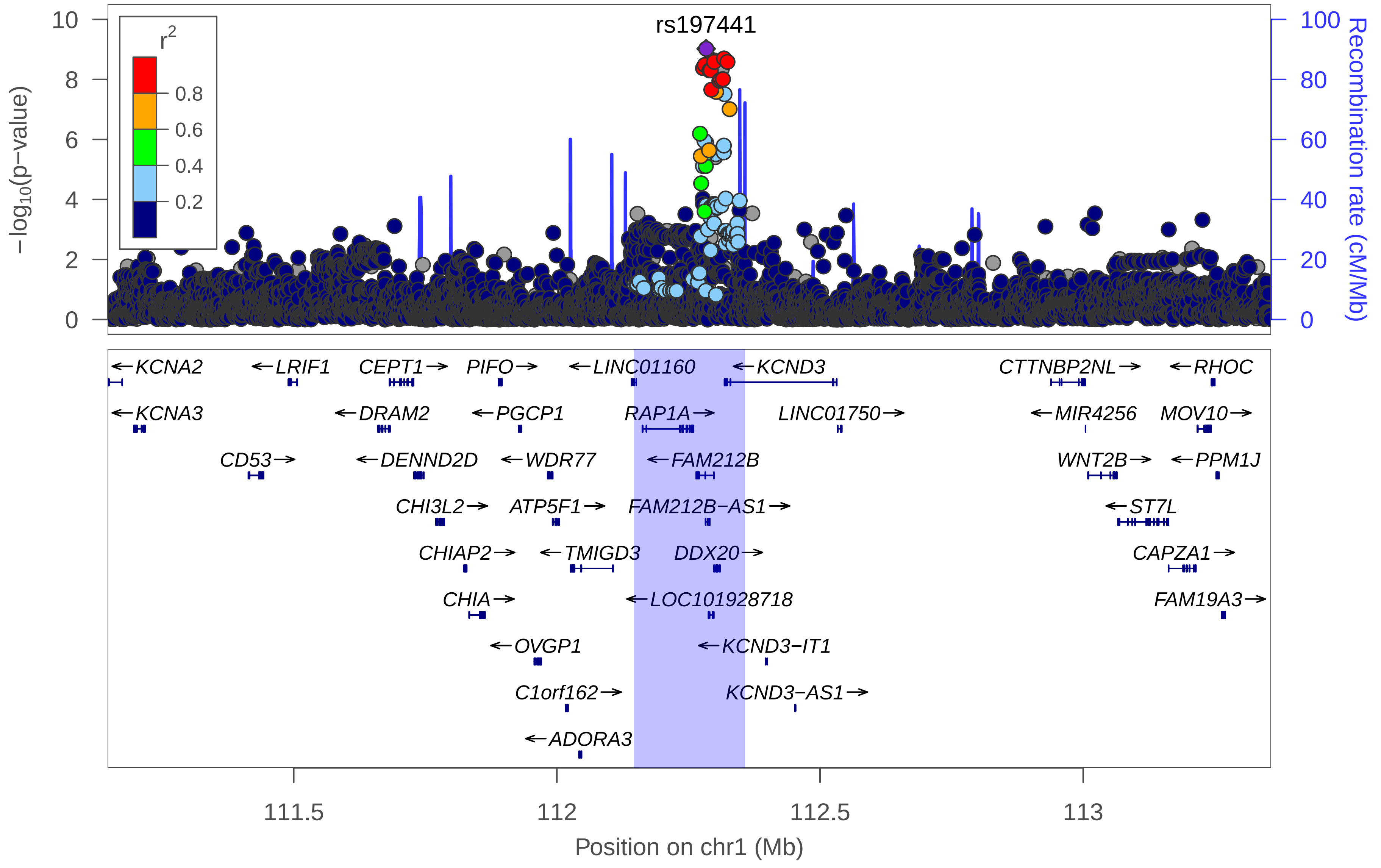

| | |
|---|---|
| date: | Mon Dec 10 07:16:54 2018 |
| build: | hg19 |
| display range: | chr1:111146255−113357144 [111146255−113357144] |
| hilite range: | 112.146Mb − 112.357Mb [ 112146000 − 112357000 ] |
| reference SNP: | chr1:112283655 |
| number of SNPs plotted: | 8017 |
| min P−value: | 9.5E−10 [chr1:112283655] |
| max P−value: | 1E0 [chr1:111339808] |

Make more plots at http://csg.sph.umich.edu/locuszoom/
