## Supplementary Figure 5 for "Genome-wide Association Study of Multisite Chronic Pain in UK Biobank"

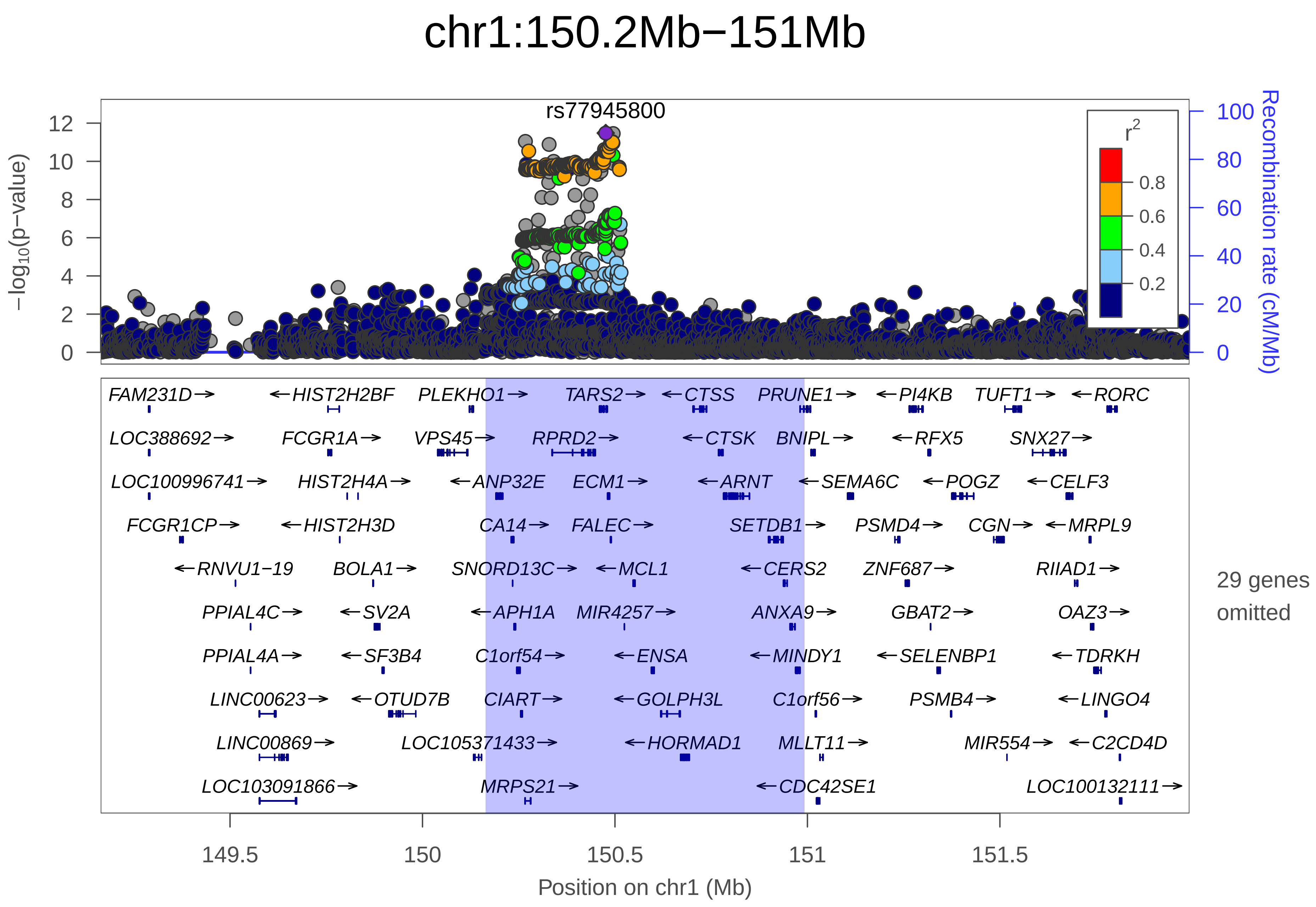
chr1:150.2Mb−151Mb
rs77945800
−log10(p−value)
Recombination rate (cM/Mb)
r2
0.8
0.6
0.4
0.2
FAM231D→ ←HIST2H2BF PLEKHO1→ TARS2→ ←CTSS PRUNE1→ ←PI4KB TUFT1→ ←RORC
LOC388692→ FCGR1A→ VPS45→ RPRD2→ ←CTSK BNIPL→ ←RFX5 SNX27→
LOC100996741→ HIST2H4A→ ←ANP32E ECM1→ ←ARNT ←SEMA6C ←POGZ ←CELF3
FCGR1CP→ ←HIST2H3D CA14→ FALEC→ SETDB1→ PSMD4→ CGN→ ←MRPL9
←RNVU1−19 BOLA1→ SNORD13C→ ←MCL1 ←CERS2 ZNF687→ RIIAD1→
PPIAL4C→ ←SV2A ←APH1A MIR4257→ ANXA9→ GBAT2→ OAZ3→
PPIAL4A→ ←SF3B4 C1orf54→ ←ENSA ←MINDY1 ←SELENBP1 ←TDRKH
LINC00623→ ←OTUD7B CIART→ ←GOLPH3L C1orf56→ PSMB4→ ←LINGO4
LINC00869→ LOC105371433→ ←HORMAD1 MLLT11→ MIR554→ ←C2CD4D
LOC103091866→ MRPS21→ ←CDC42SE1 LOC100132111→
29 genes omitted
Position on chr1 (Mb)

| | |
|---|---|
| date: | Mon Dec 10 07:36:07 2018 |
| build: | hg19 |
| display range: | chr1:149163915−151992437 [149163915−151992437] |
| hilite range: | 150.164Mb − 150.992Mb [ 150164000 − 150992000 ] |
| reference SNP: | chr1:150476049 |
| number of SNPs plotted: | 6387 |
| min P−value: | 3.3E−12 [chr1:150476049] |
| max P−value: | 1E0 [chr1:149219722] |
| omitted Genes: | RNVU1−20, LOC101929798, HIST2H4B |
| omitted Genes: | HIST2H3A, HIST2H3C, HIST2H2AA4 |
| omitted Genes: | HIST2H2AA3, HIST2H2BC, HIST2H2BE |
| omitted Genes: | HIST2H2AC, HIST2H2AB, MTMR11 |
| omitted Genes: | PRPF3, MIR6878, ADAMTSL4 |
| omitted Genes: | ADAMTSL4−AS1, GABPB2, TNFAIP8L2 |
| omitted Genes: | TNFAIP8L2−SCNM1, LYSMD1, SCNM1 |
| omitted Genes: | TMOD4, VPS72, PIP5K1A |
| omitted Genes: | LOC100507670, THEM5, THEM4 |
| omitted Genes: | S100A10, NBPF18P |

Make more plots at http://csg.sph.umich.edu/locuszoom/
