## Supplementary Figure 6 for "Genome-wide Association Study of Multisite Chronic Pain in UK Biobank"

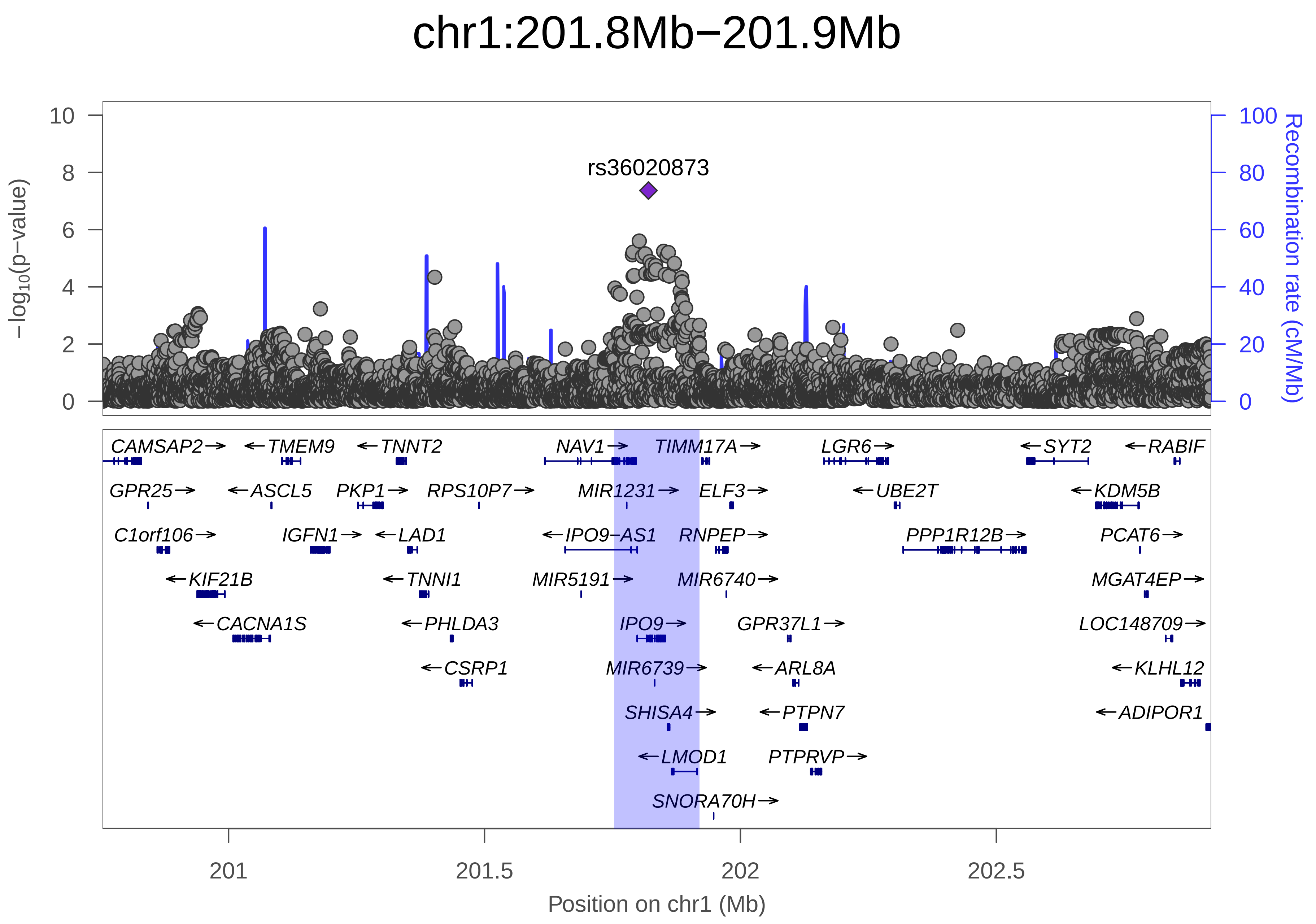

chr1:201.8Mb−201.9Mb
rs36020873
−log10(p−value)
Recombination rate (cM/Mb)
0
2
4
6
8
10
0
20
40
60
80
100
CAMSAP2→
←TMEM9
←TNNT2
NAV1→
TIMM17A→
LGR6→
←SYT2
←RABIF
GPR25→
←ASCL5
PKP1→
RPS10P7→
MIR1231→
ELF3→
←UBE2T
←KDM5B
C1orf106→
IGFN1→
←LAD1
←IPO9−AS1
RNPEP→
PPP1R12B→
PCAT6→
←KIF21B
←TNNI1
MIR5191→
MIR6740→
MGAT4EP→
←CACNA1S
←PHLDA3
IPO9→
GPR37L1→
LOC148709→
←CSRP1
MIR6739→
←ARL8A
←KLHL12
SHISA4→
←PTPN7
←ADIPOR1
←LMOD1
PTPRVP→
SNORA70H→
201
201.5
202
202.5
Position on chr1 (Mb)

| | |
|---|---|
| date: | Mon Dec 10 08:03:18 2018 |
| build: | hg19 |
| display range: | chr1:200753696−202919960 [200753696−202919960] |
| hilite range: | 201.754Mb − 201.92Mb [ 201754000 − 201920000 ] |
| reference SNP: | chr1:201820307 |
| number of SNPs plotted: | 7407 |
| min P−value: | 4.3E−8 [chr1:201820307] |
| max P−value: | 1E0 [chr1:200753725] |
| Warning: | No usable LD information for reference SNP. |

Make more plots at http://csg.sph.umich.edu/locuszoom/
