## Supplementary Figure 7 for "Genome-wide Association Study of Multisite Chronic Pain in UK Biobank"

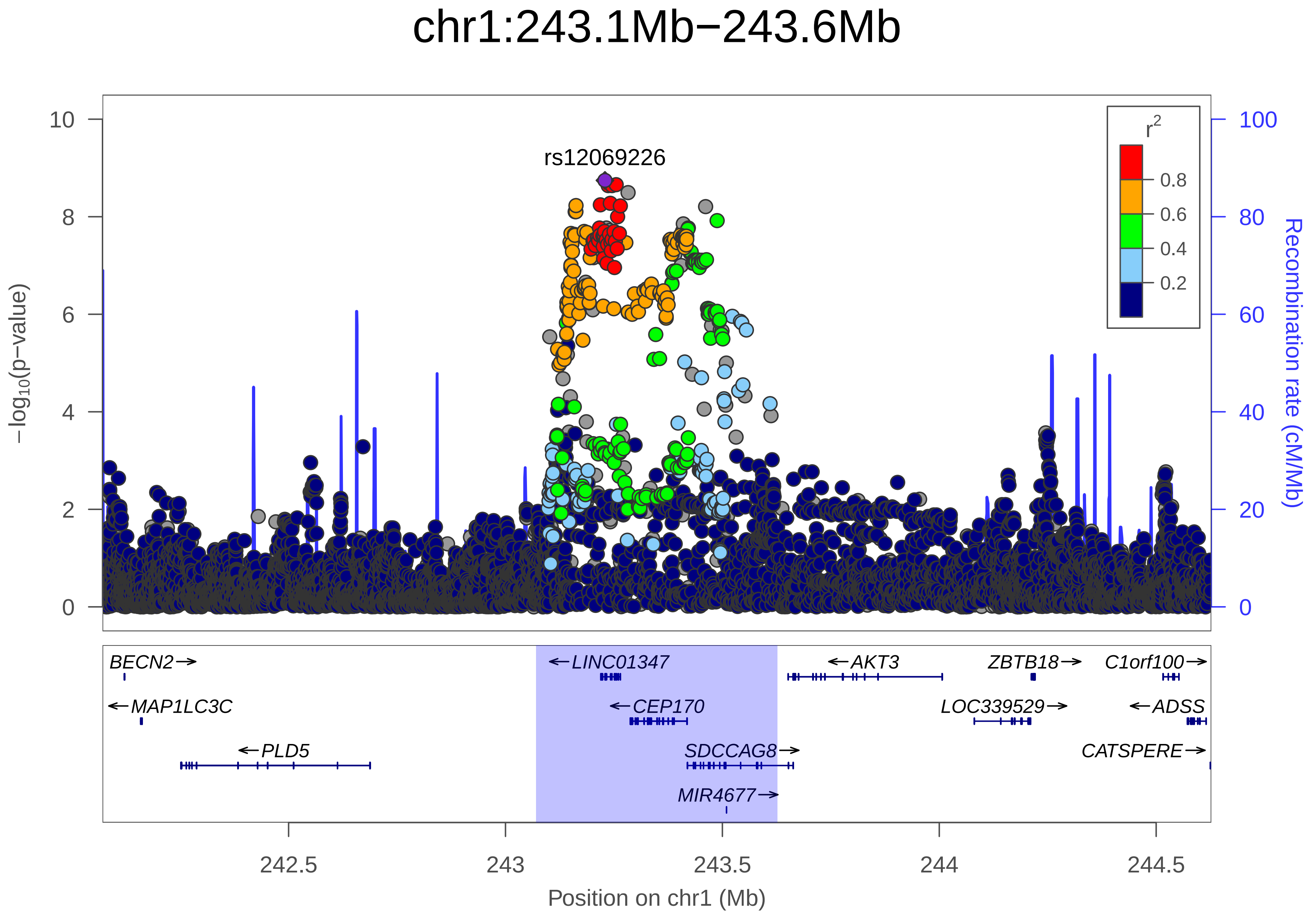
chr1:243.1Mb−243.6Mb
rs12069226
r2
0.8
0.6
0.4
0.2
−log10(p−value)
Recombination rate (cM/Mb)
0
2
4
6
8
10
0
20
40
60
80
100
BECN2→
←LINC01347
←AKT3
ZBTB18→
C1orf100→
←MAP1LC3C
←CEP170
LOC339529→
←ADSS
←PLD5
SDCCAG8→
CATSPERE→
MIR4677→
242.5
243
243.5
244
244.5
Position on chr1 (Mb)

| | |
|---|---|
| date: | Mon Dec 10 08:33:54 2018 |
| build: | hg19 |
| display range: | chr1:242070992−244627135 [242070992−244627135] |
| hilite range: | 243.071Mb − 243.627Mb [ 243071000 − 243627000 ] |
| reference SNP: | chr1:243229308 |
| number of SNPs plotted: | 9367 |
| min P−value: | 1.8E−9 [chr1:243229308] |
| max P−value: | 1E0 [chr1:242080136] |

Make more plots at http://csg.sph.umich.edu/locuszoom/
