## Supplementary Figure 8 for "Genome-wide Association Study of Multisite Chronic Pain in UK Biobank"

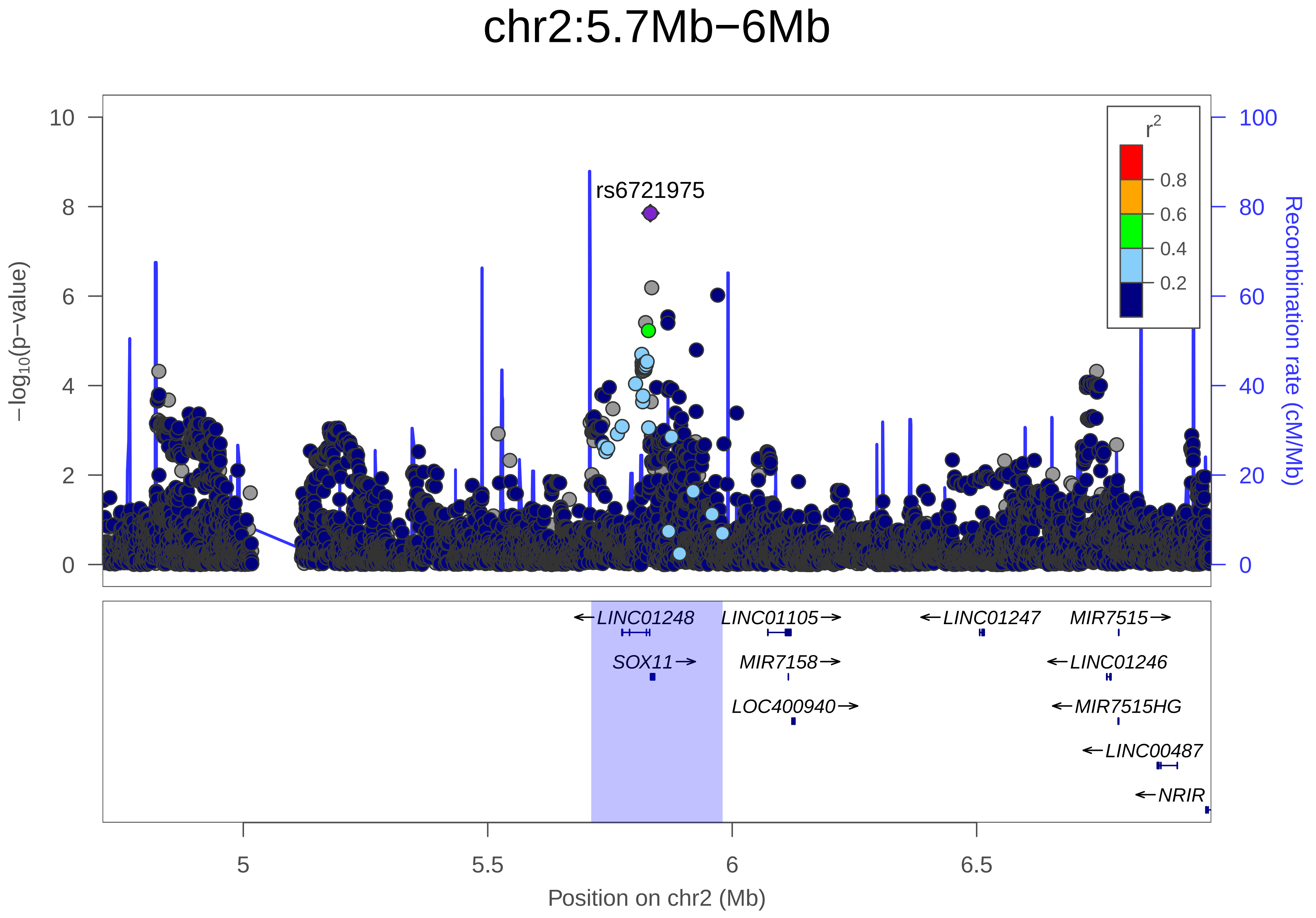

chr2:5.7Mb−6Mb
−log10(p−value)
Recombination rate (cM/Mb)
rs6721975
r2
0.8
0.6
0.4
0.2
LINC01248
LINC01105
LINC01247
MIR7515
SOX11
MIR7158
LINC01246
LOC400940
MIR7515HG
LINC00487
NRIR
5
5.5
6
6.5
Position on chr2 (Mb)

| | |
|---|---|
| date: | Mon Dec 10 07:02:39 2018 |
| build: | hg19 |
| display range: | chr2:4712068−6979944 [4712068−6979944] |
| hilite range: | 5.712Mb − 5.98Mb [ 5712000 − 5980000 ] |
| reference SNP: | chr2:5832667 |
| number of SNPs plotted: | 9093 |
| min P−value: | 1.4E−8 [chr2:5832667] |
| max P−value: | 1E0 [chr2:4789141] |

Make more plots at http://csg.sph.umich.edu/locuszoom/
