## Supplementary Figure 9 for "Genome-wide Association Study of Multisite Chronic Pain in UK Biobank"

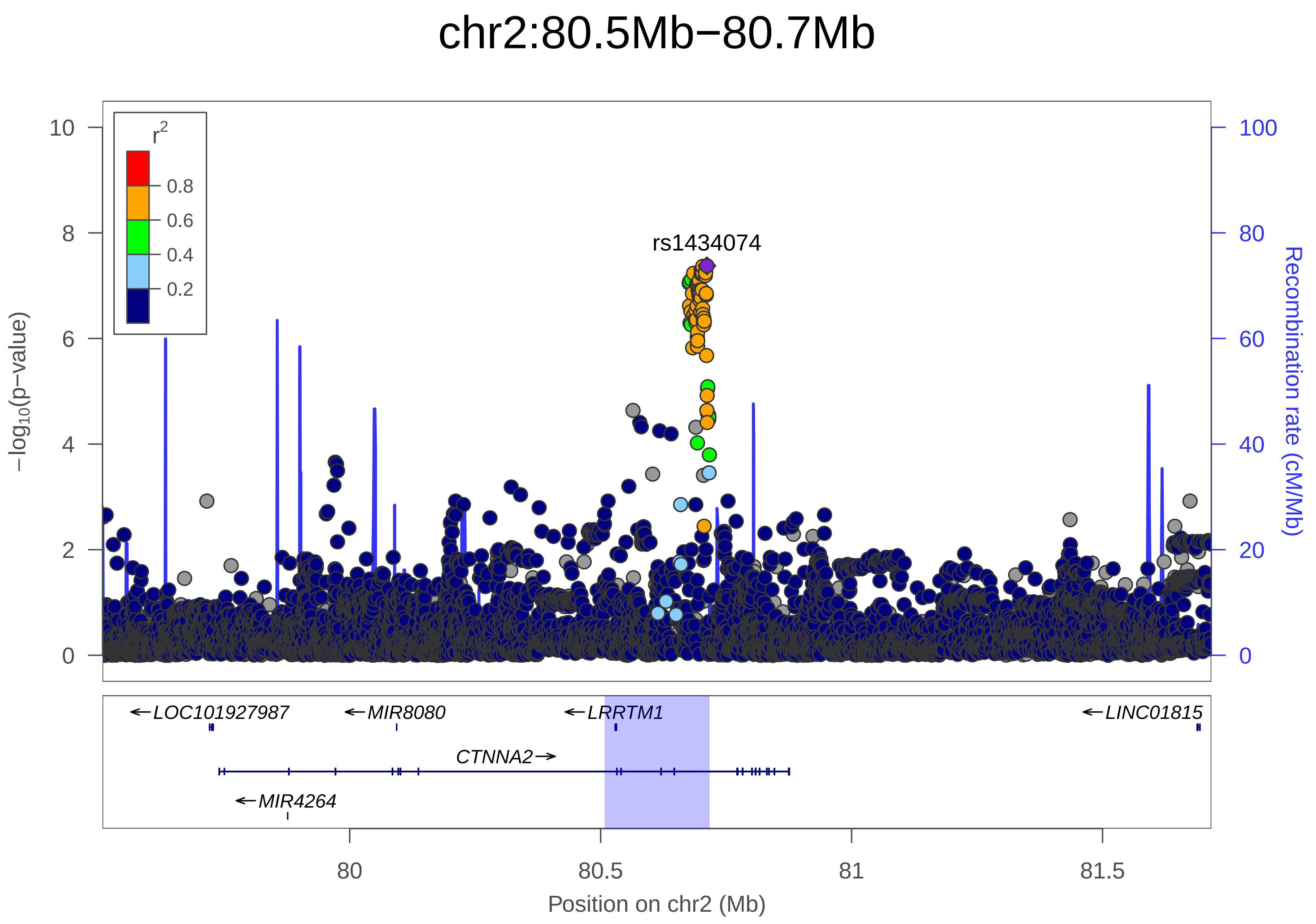

chr2:80.5Mb−80.7Mb
r2
0.8
0.6
0.4
0.2
rs1434074
−log10(p−value)
Recombination rate (cM/Mb)
0
2
4
6
8
10
0
20
40
60
80
100
←LOC101927987
←MIR8080
←LRRTM1
←LINC01815
CTNNA2→
←MIR4264
80
80.5
81
81.5
Position on chr2 (Mb)

| | |
|---|---|
| date: | Mon Dec 10 07:16:53 2018 |
| build: | hg19 |
| display range: | chr2:79507517−81716691 [79507517−81716691] |
| hilite range: | 80.508Mb − 80.717Mb [ 80508000 − 80717000 ] |
| reference SNP: | chr2:80711752 |
| number of SNPs plotted: | 7782 |
| min P−value: | 4.2E−8 [chr2:80711752] |
| max P−value: | 1E0 [chr2:79511882] |

Make more plots at http://csg.sph.umich.edu/locuszoom/
