## Supplementary Figure 10 for "Genome-wide Association Study of Multisite Chronic Pain in UK Biobank"

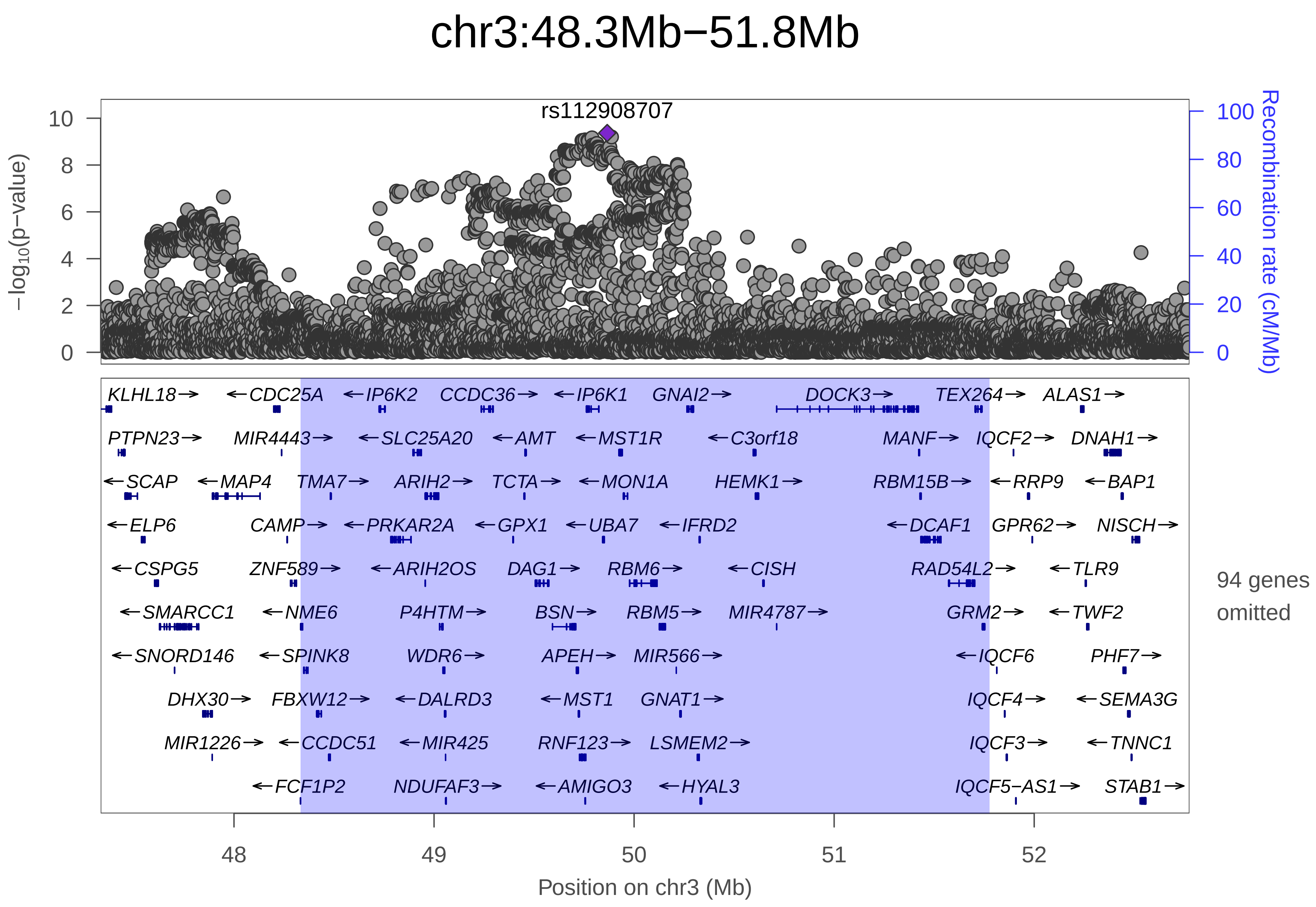

chr3:48.3Mb−51.8Mb
rs112908707
−log10(p−value)
Recombination rate (cM/Mb)
0
2
4
6
8
10
0
20
40
60
80
100
KLHL18→
←CDC25A
←IP6K2
CCDC36→
←IP6K1
GNAI2→
DOCK3→
TEX264→
ALAS1→
PTPN23→
MIR4443→
←SLC25A20
←AMT
←MST1R
←C3orf18
MANF→
IQCF2→
DNAH1→
←SCAP
←MAP4
TMA7→
ARIH2→
TCTA→
←MON1A
HEMK1→
RBM15B→
←RRP9
←BAP1
←ELP6
CAMP→
←PRKAR2A
←GPX1
←UBA7
←IFRD2
←DCAF1
GPR62→
NISCH→
←CSPG5
ZNF589→
←ARIH2OS
DAG1→
RBM6→
←CISH
RAD54L2→
←TLR9
←SMARCC1
←NME6
P4HTM→
BSN→
RBM5→
MIR4787→
GRM2→
←TWF2
←SNORD146
←SPINK8
WDR6→
APEH→
MIR566→
←IQCF6
PHF7→
DHX30→
FBXW12→
←DALRD3
←MST1
GNAT1→
IQCF4→
←SEMA3G
MIR1226→
←CCDC51
←MIR425
RNF123→
LSMEM2→
IQCF3→
←TNNC1
←FCF1P2
NDUFAF3→
←AMIGO3
←HYAL3
IQCF5−AS1→
STAB1→
94 genes omitted
48
49
50
51
52
Position on chr3 (Mb)

| | |
|---|---|
| date: | Mon Dec 10 08:03:21 2018 |
| build: | hg19 |
| display range: | chr3:47333477−52776104 [47333477−52776104] |
| hilite range: | 48.333Mb − 51.776Mb [ 48333000 − 51776000 ] |
| reference SNP: | chr3:49865628 |
| number of SNPs plotted: | 9922 |
| min P−value: | 4.3E−10 [chr3:49865628] |
| max P−value: | 1E0 [chr3:48407462] |
| omitted Genes: | MIR2115, PLXNB1, ATRIP |
| omitted Genes: | TREX1, SHISA5, PFKFB4 |
| omitted Genes: | MIR6823, UCN2, COL7A1 |
| omitted Genes: | MIR711, UQCRC1, SNORA94 |
| omitted Genes: | TMEM89, SLC26A6, MIR6824 |
| omitted Genes: | CELSR3, MIR4793, CELSR3−AS1 |
| omitted Genes: | NCKIPSD, PRKAR2A−AS1, MIR191 |
| omitted Genes: | IMPDH2, QRICH1, QARS |
| omitted Genes: | MIR6890, USP19, LAMB2 |
| omitted Genes: | LAMB2P1, CCDC71, KLHDC8B |
| omitted Genes: | C3orf84, C3orf62, MIR4271 |
| omitted Genes: | USP4, RHOA, NICN1 |
| omitted Genes: | BSN−AS2, GMPPB, CDHR4 |
| omitted Genes: | FAM212A, MIR5193, TRAIP |
| omitted Genes: | CAMKV, RBM5−AS1, SEMA3F−AS1 |

Make more plots at http://csg.sph.umich.edu/locuszoom/
