## Supplementary Figure 11 for "Genome-wide Association Study of Multisite Chronic Pain in UK Biobank"

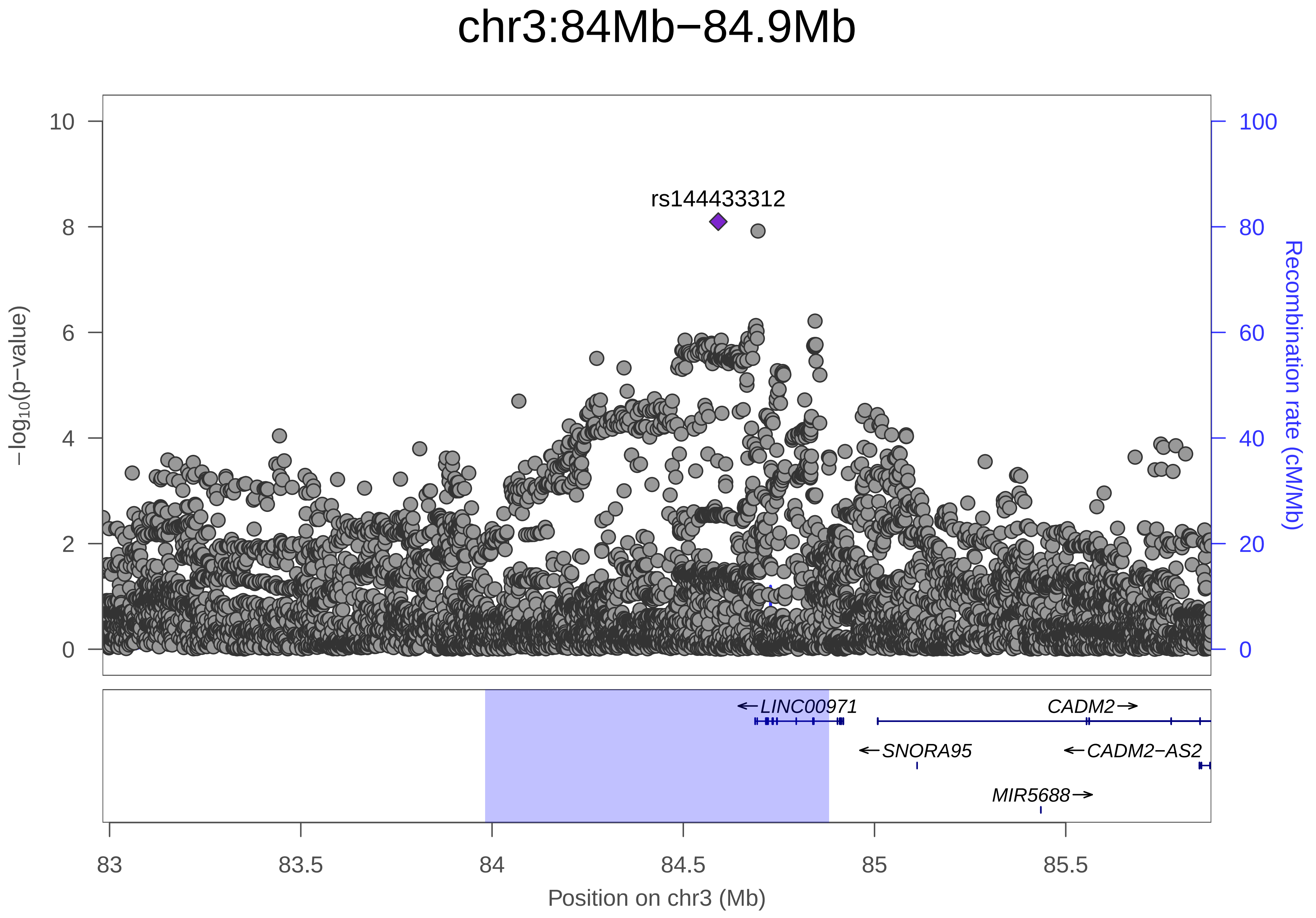
chr3:84Mb−84.9Mb
rs144433312
−log10(p−value)
Recombination rate (cM/Mb)
10
8
6
4
2
0
100
80
60
40
20
0
←LINC00971
CADM2→
←SNORA95
←CADM2−AS2
MIR5688→
83
83.5
84
84.5
85
85.5
Position on chr3 (Mb)

| | |
|---|---|
| date: | Mon Dec 10 08:35:05 2018 |
| build: | hg19 |
| display range: | chr3:82981526−85880591 [82981526−85880591] |
| hilite range: | 83.982Mb − 84.881Mb [ 83982000 − 84881000 ] |
| reference SNP: | chr3:84591507 |
| number of SNPs plotted: | 9938 |
| min P−value: | 8E−9 [chr3:84591507] |
| max P−value: | 1E0 [chr3:83781331] |
| Warning: | No usable LD information for reference SNP. |

Make more plots at http://csg.sph.umich.edu/locuszoom/
