## Supplementary Figure 12 for "Genome-wide Association Study of Multisite Chronic Pain in UK Biobank"

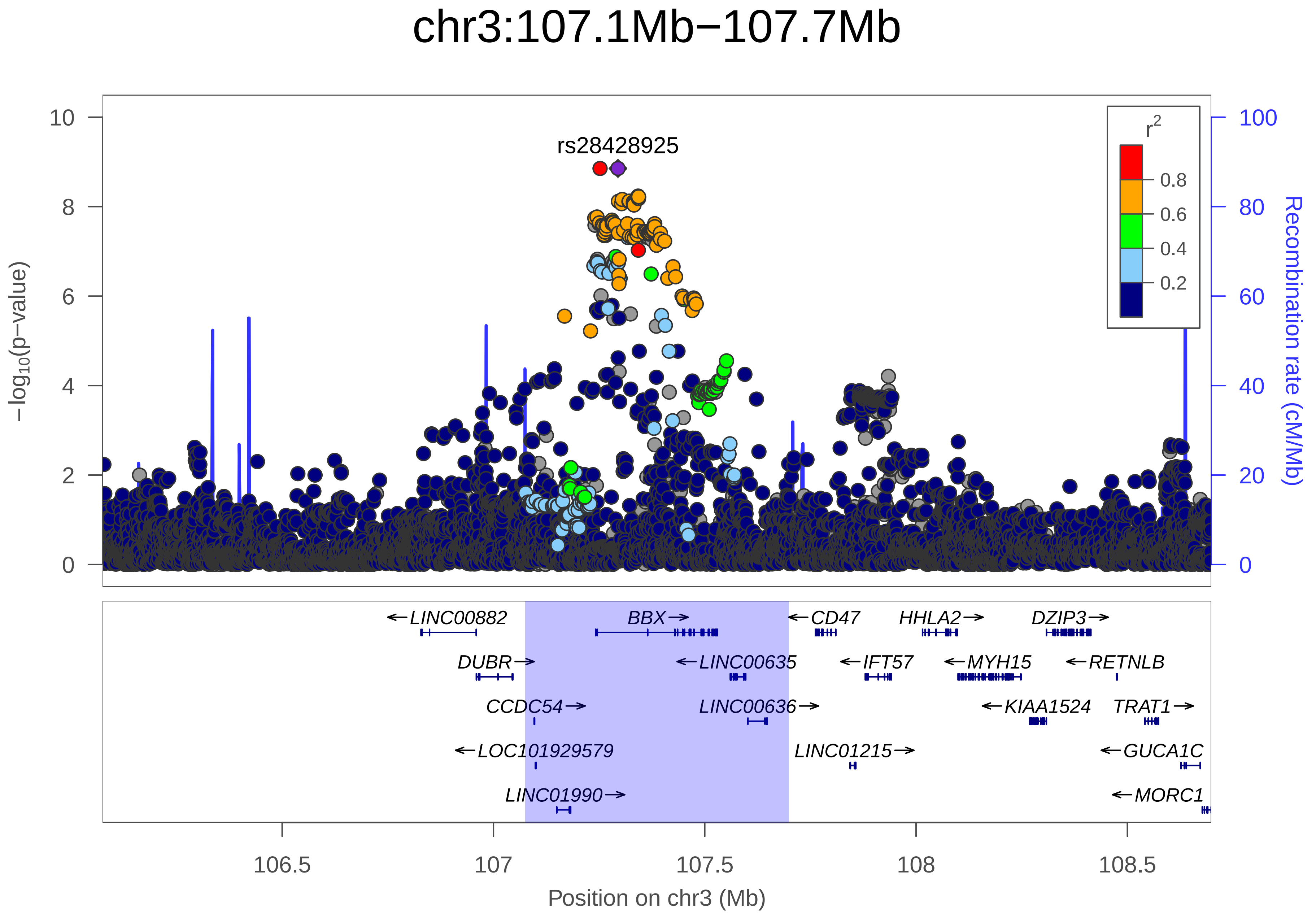

chr3:107.1Mb−107.7Mb
rs28428925
r²
0.8
0.6
0.4
0.2
10
8
6
4
2
0
100
80
60
40
20
0
−log10(p−value)
Recombination rate (cM/Mb)
LINC00882
BBX
CD47
HHLA2
DZIP3
DUBR
LINC00635
IFT57
MYH15
RETNLB
CCDC54
LINC00636
KIAA1524
TRAT1
LOC101929579
LINC01215
GUCA1C
LINC01990
MORC1
106.5
107
107.5
108
108.5
Position on chr3 (Mb)

| | |
|---|---|
| date: | Mon Dec 10 07:02:39 2018 |
| build: | hg19 |
| display range: | chr3:106074910−108698638 [106074910−108698638] |
| hilite range: | 107.075Mb − 107.699Mb [ 107075000 − 107699000 ] |
| reference SNP: | chr3:107294634 |
| number of SNPs plotted: | 8580 |
| min P−value: | 1.4E−9 [chr3:107252190] |
| max P−value: | 1E0 [chr3:106141516] |

Make more plots at http://csg.sph.umich.edu/locuszoom/
