## Supplementary Figure 13 for "Genome-wide Association Study of Multisite Chronic Pain in UK Biobank"

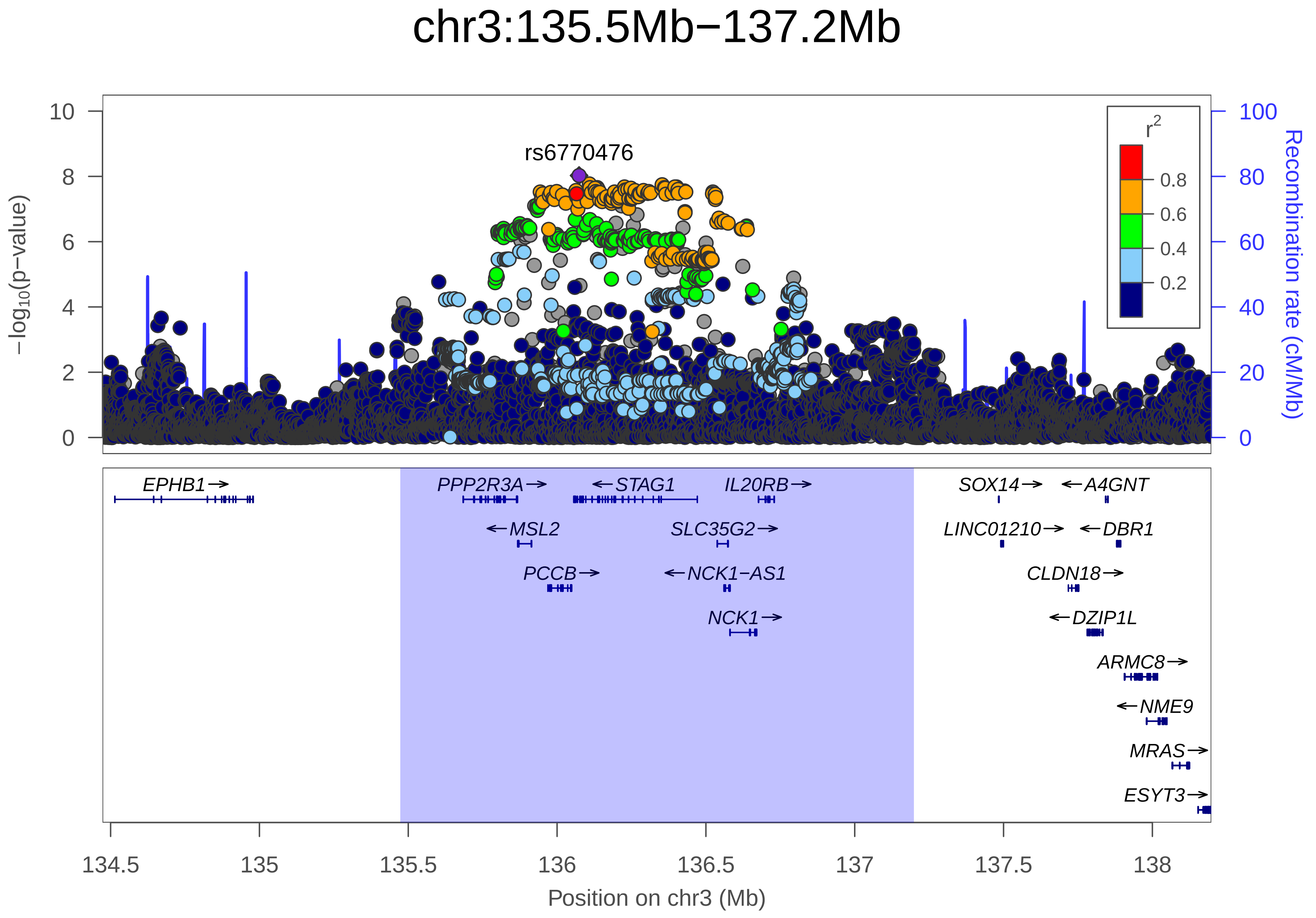
chr3:135.5Mb−137.2Mb
rs6770476
−log10(p−value)
Recombination rate (cM/Mb)
r2
0.8
0.6
0.4
0.2
10
8
6
4
2
0
100
80
60
40
20
0
EPHB1→
PPP2R3A→
←STAG1
IL20RB→
SOX14→
←A4GNT
←MSL2
SLC35G2→
LINC01210→
←DBR1
PCCB→
←NCK1−AS1
CLDN18→
NCK1→
←DZIP1L
ARMC8→
←NME9
MRAS→
ESYT3→
134.5
135
135.5
136
136.5
137
137.5
138
Position on chr3 (Mb)

| | |
|---|---|
| date: | Mon Dec 10 07:27:20 2018 |
| build: | hg19 |
| display range: | chr3:134472797−138198063 [134472797−138198063] |
| hilite range: | 135.473Mb − 137.198Mb [ 135473000 − 137198000 ] |
| reference SNP: | chr3:136073920 |
| number of SNPs plotted: | 10150 |
| min P−value: | 9.4E−9 [chr3:136073920] |
| max P−value: | 1E0 [chr3:134502897] |

Make more plots at http://csg.sph.umich.edu/locuszoom/
