## Supplementary Figure 14 for "Genome-wide Association Study of Multisite Chronic Pain in UK Biobank"

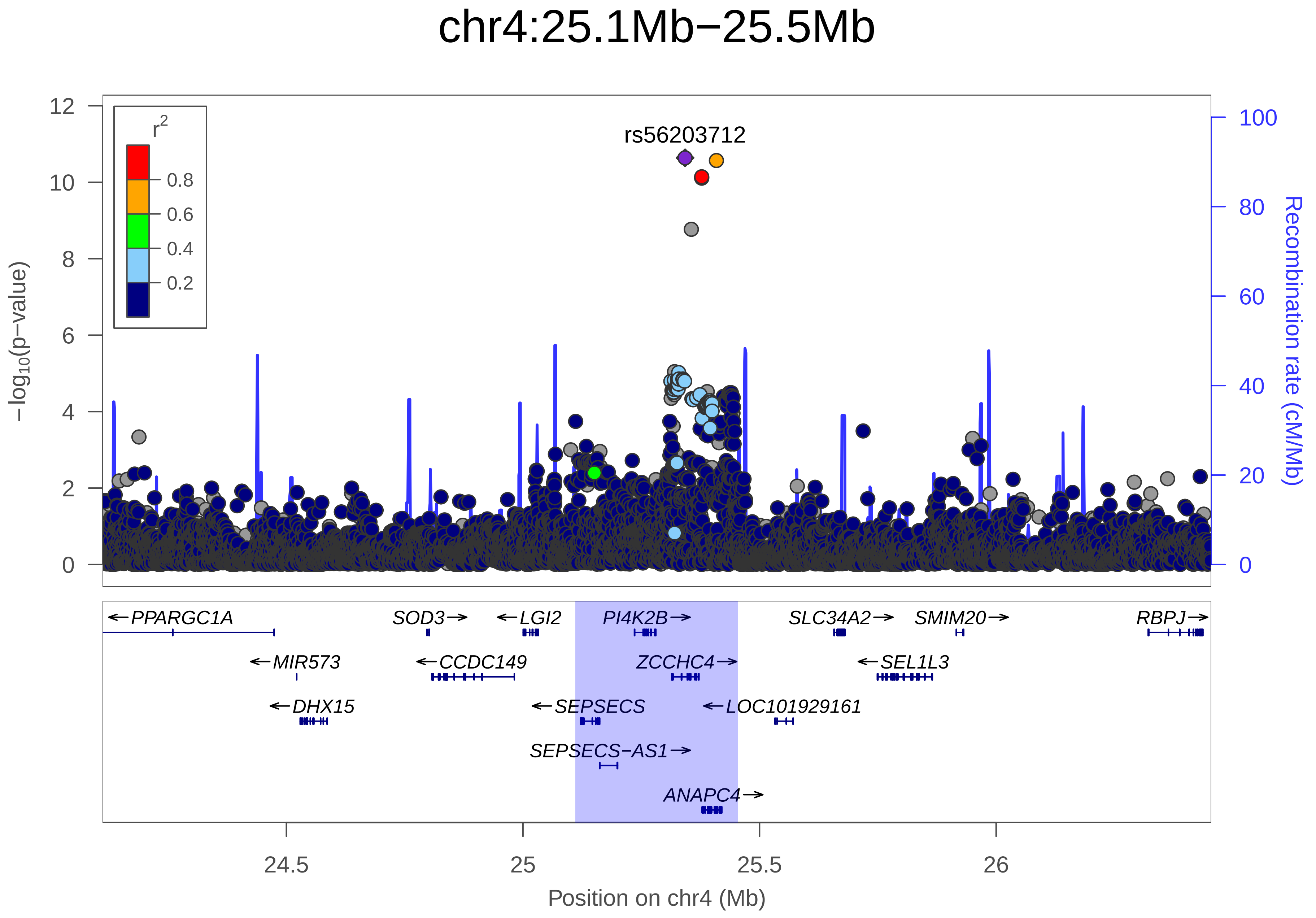
chr4:25.1Mb−25.5Mb
rs56203712
r2
0.8
0.6
0.4
0.2
−log10(p−value)
Recombination rate (cM/Mb)
PPARGC1A
SOD3
LGI2
PI4K2B
SLC34A2
SMIM20
RBPJ
MIR573
CCDC149
ZCCHC4
SEL1L3
DHX15
SEPSECS
LOC101929161
SEPSECS−AS1
ANAPC4
24.5
25
25.5
26
Position on chr4 (Mb)

| | |
|---|---|
| date: | Mon Dec 10 07:55:50 2018 |
| build: | hg19 |
| display range: | chr4:24111387−26454781 [24111387−26454781] |
| hilite range: | 25.111Mb − 25.455Mb [ 25111000 − 25455000 ] |
| reference SNP: | chr4:25342606 |
| number of SNPs plotted: | 8601 |
| min P−value: | 2.3E−11 [chr4:25342606] |
| max P−value: | 1E0 [chr4:24176976] |

Make more plots at http://csg.sph.umich.edu/locuszoom/
