## Supplementary Figure 15 for "Genome-wide Association Study of Multisite Chronic Pain in UK Biobank"

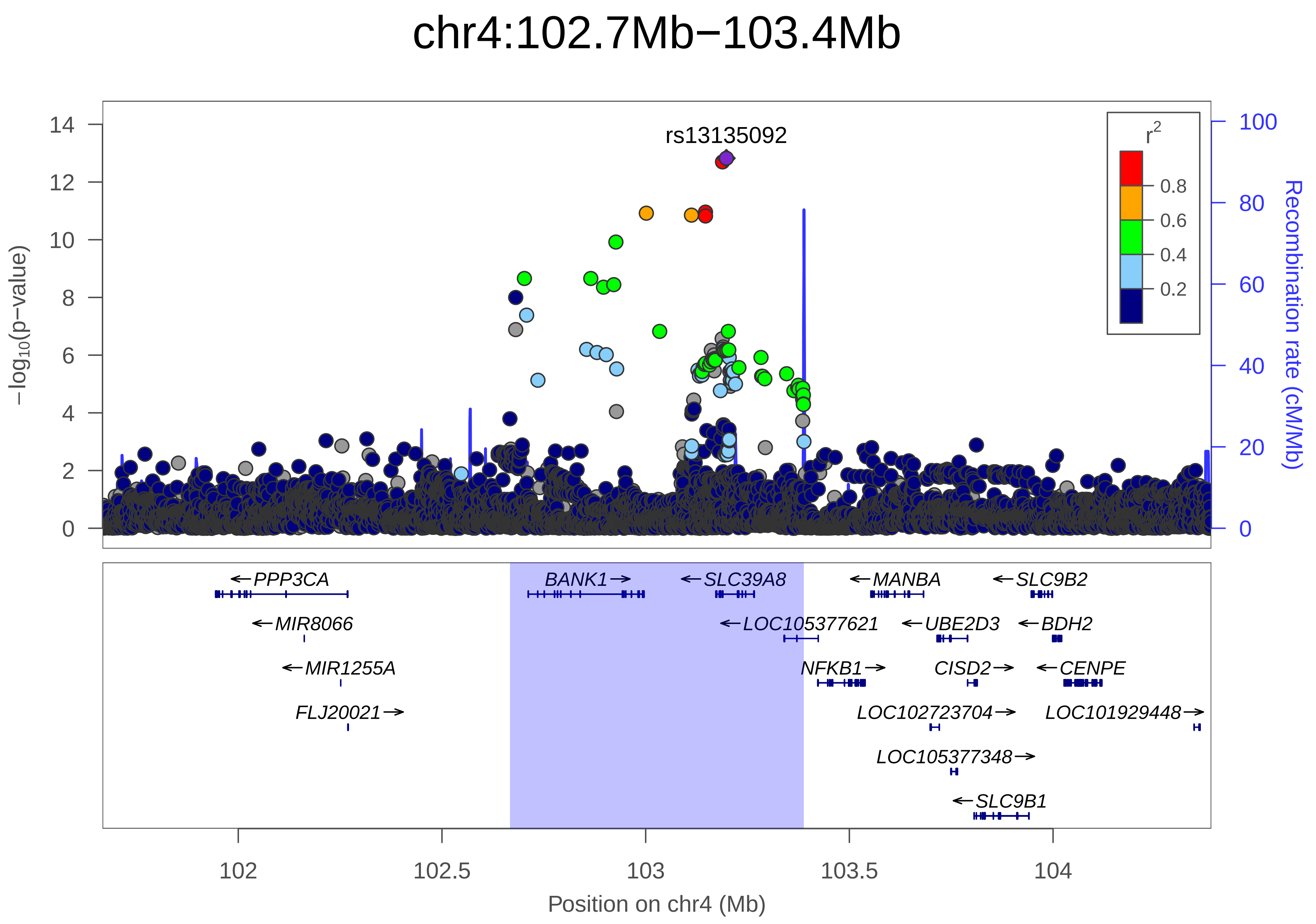
chr4:102.7Mb−103.4Mb
rs13135092
r2
0.8
0.6
0.4
0.2
14
12
10
8
6
4
2
0
100
80
60
40
20
0
−log10(p−value)
Recombination rate (cM/Mb)
PPP3CA
MIR8066
MIR1255A
FLJ20021
BANK1
SLC39A8
LOC105377621
NFKB1
MANBA
UBE2D3
CISD2
LOC102723704
LOC105377348
SLC9B1
SLC9B2
BDH2
CENPE
LOC101929448
102
102.5
103
103.5
104
Position on chr4 (Mb)

| | |
|---|---|
| date: | Mon Dec 10 08:13:58 2018 |
| build: | hg19 |
| display range: | chr4:101666785−104388441 [101666785−104388441] |
| hilite range: | 102.667Mb − 103.388Mb [ 102667000 − 103388000 ] |
| reference SNP: | chr4:103198082 |
| number of SNPs plotted: | 8557 |
| min P−value: | 1.5E−13 [chr4:103198082] |
| max P−value: | 1E0 [chr4:101914313] |

Make more plots at http://csg.sph.umich.edu/locuszoom/
