## Supplementary Figure 16 for "Genome-wide Association Study of Multisite Chronic Pain in UK Biobank"

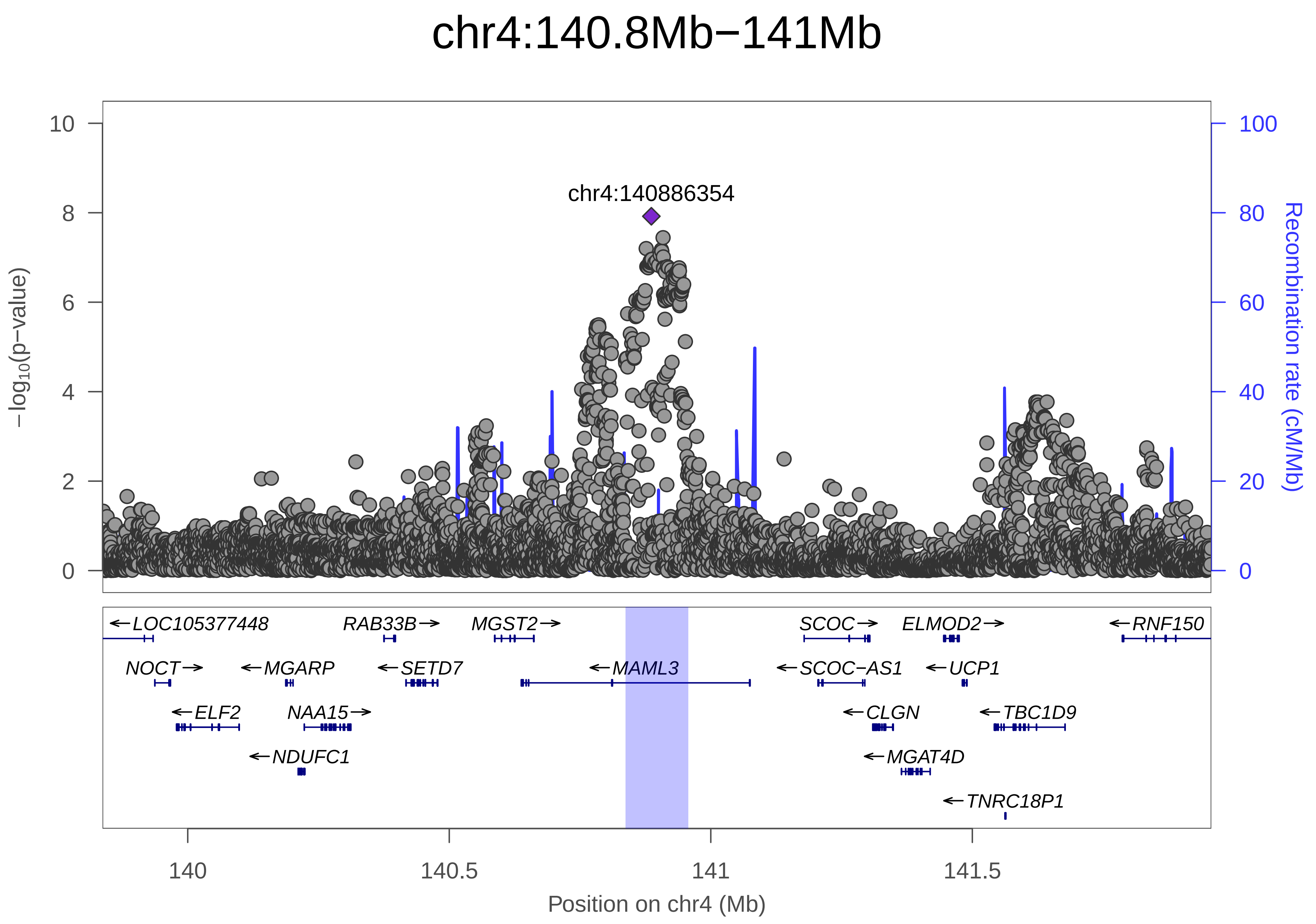

chr4:140.8Mb−141Mb
chr4:140886354
10
8
6
4
2
0
−log10(p−value)
100
80
60
40
20
0
Recombination rate (cM/Mb)
←LOC105377448
RAB33B→
MGST2→
SCOC→
ELMOD2→
←RNF150
NOCT→
←MGARP
←SETD7
←MAML3
←SCOC−AS1
←UCP1
←ELF2
NAA15→
←CLGN
←TBC1D9
←NDUFC1
←MGAT4D
←TNRC18P1
140
140.5
141
141.5
Position on chr4 (Mb)

| | |
|---|---|
| date: | Mon Dec 10 07:02:39 2018 |
| build: | hg19 |
| display range: | chr4:139837043−141956936 [139837043−141956936] |
| hilite range: | 140.837Mb − 140.957Mb [ 140837000 − 140957000 ] |
| reference SNP: | chr4:140886354 |
| number of SNPs plotted: | 6925 |
| min P−value: | 1.2E−8 [chr4:140886354] |
| max P−value: | 1E0 [chr4:139842608] |
| Warning: | No usable LD information for reference SNP. |

Make more plots at http://csg.sph.umich.edu/locuszoom/
