## Supplementary Figure 17 for "Genome-wide Association Study of Multisite Chronic Pain in UK Biobank"

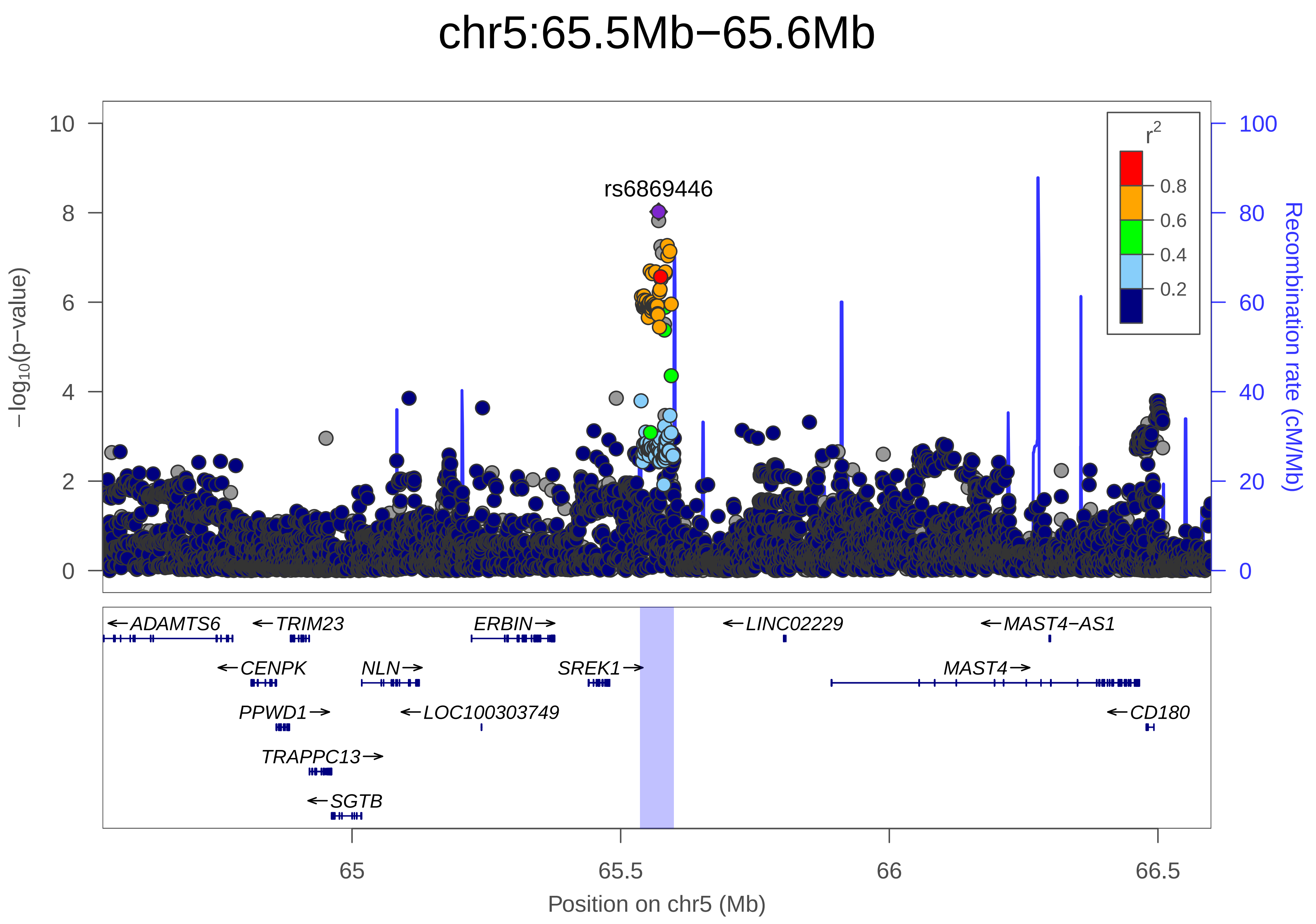

chr5:65.5Mb−65.6Mb
rs6869446
r2
0.8
0.6
0.4
0.2
−log10(p−value)
Recombination rate (cM/Mb)
10
8
6
4
2
0
100
80
60
40
20
0
←ADAMTS6
←TRIM23
ERBIN→
←LINC02229
←MAST4−AS1
←CENPK
NLN→
SREK1→
MAST4→
PPWD1→
←LOC100303749
←CD180
TRAPPC13→
←SGTB
65
65.5
66
66.5
Position on chr5 (Mb)

| | |
|---|---|
| date: | Mon Dec 10 07:16:54 2018 |
| build: | hg19 |
| display range: | chr5:64535570−66599360 [64535570−66599360] |
| hilite range: | 65.536Mb − 65.599Mb [ 65536000 − 65599000 ] |
| reference SNP: | chr5:65570607 |
| number of SNPs plotted: | 6793 |
| min P−value: | 9.5E−9 [chr5:65570607] |
| max P−value: | 1E0 [chr5:64548647] |

Make more plots at http://csg.sph.umich.edu/locuszoom/
