## Supplementary Figure 18 for "Genome-wide Association Study of Multisite Chronic Pain in UK Biobank"

### chr5:103.7Mb−104.2Mb

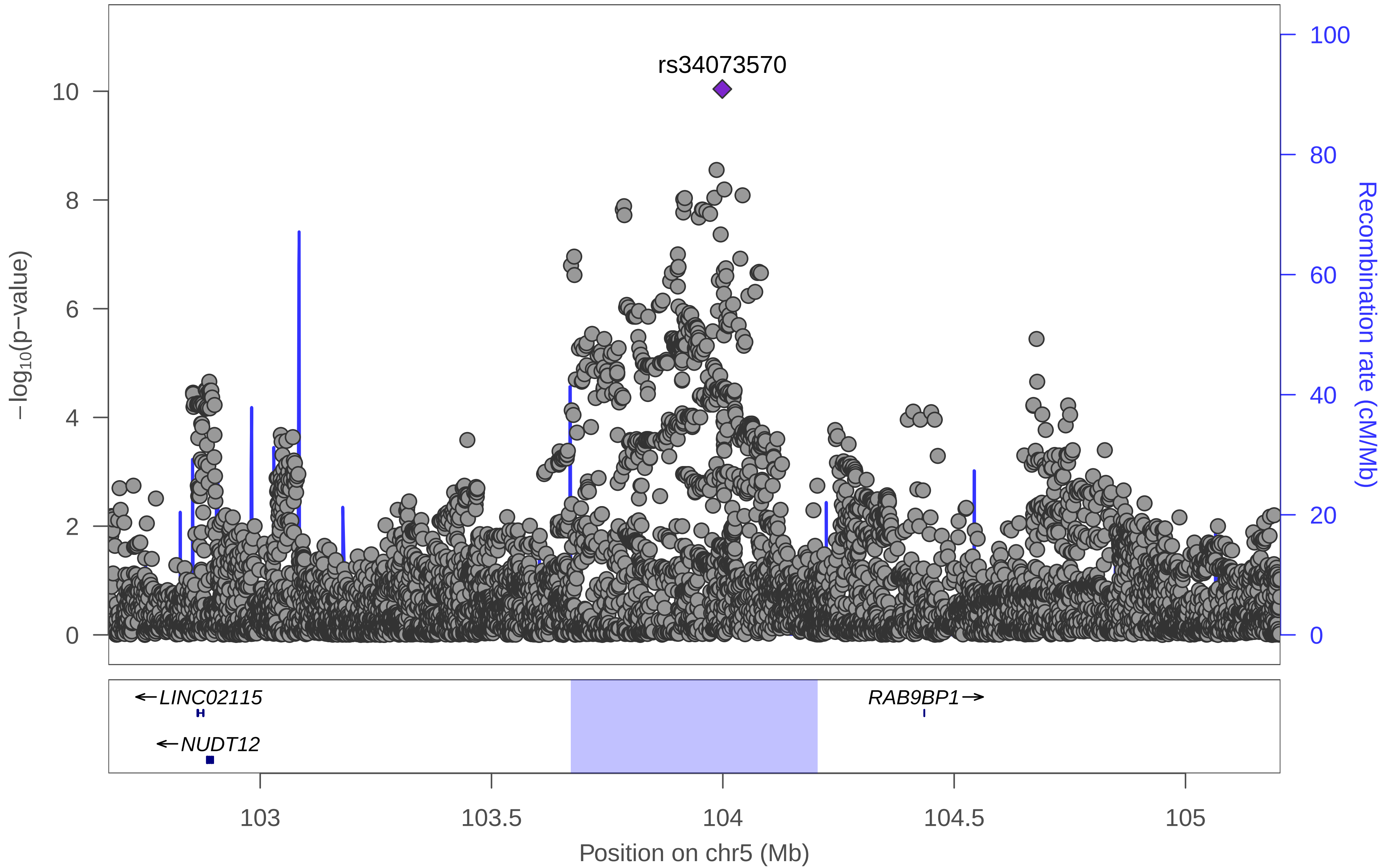

| | |
|---|---|
| date: | Mon Dec 10 07:35:09 2018 |
| build: | hg19 |
| display range: | chr5:102671867−105205240 [102671867−105205240] |
| hilite range: | 103.672Mb − 104.205Mb [ 103672000 − 104205000 ] |
| reference SNP: | chr5:103998895 |
| number of SNPs plotted: | 8835 |
| min P−value: | 9.1E−11 [chr5:103998895] |
| max P−value: | 1E0 [chr5:102750828] |
| Warning: | No usable LD information for reference SNP. |

Make more plots at http://csg.sph.umich.edu/locuszoom/
