## Supplementary Figure 19 for "Genome-wide Association Study of Multisite Chronic Pain in UK Biobank"

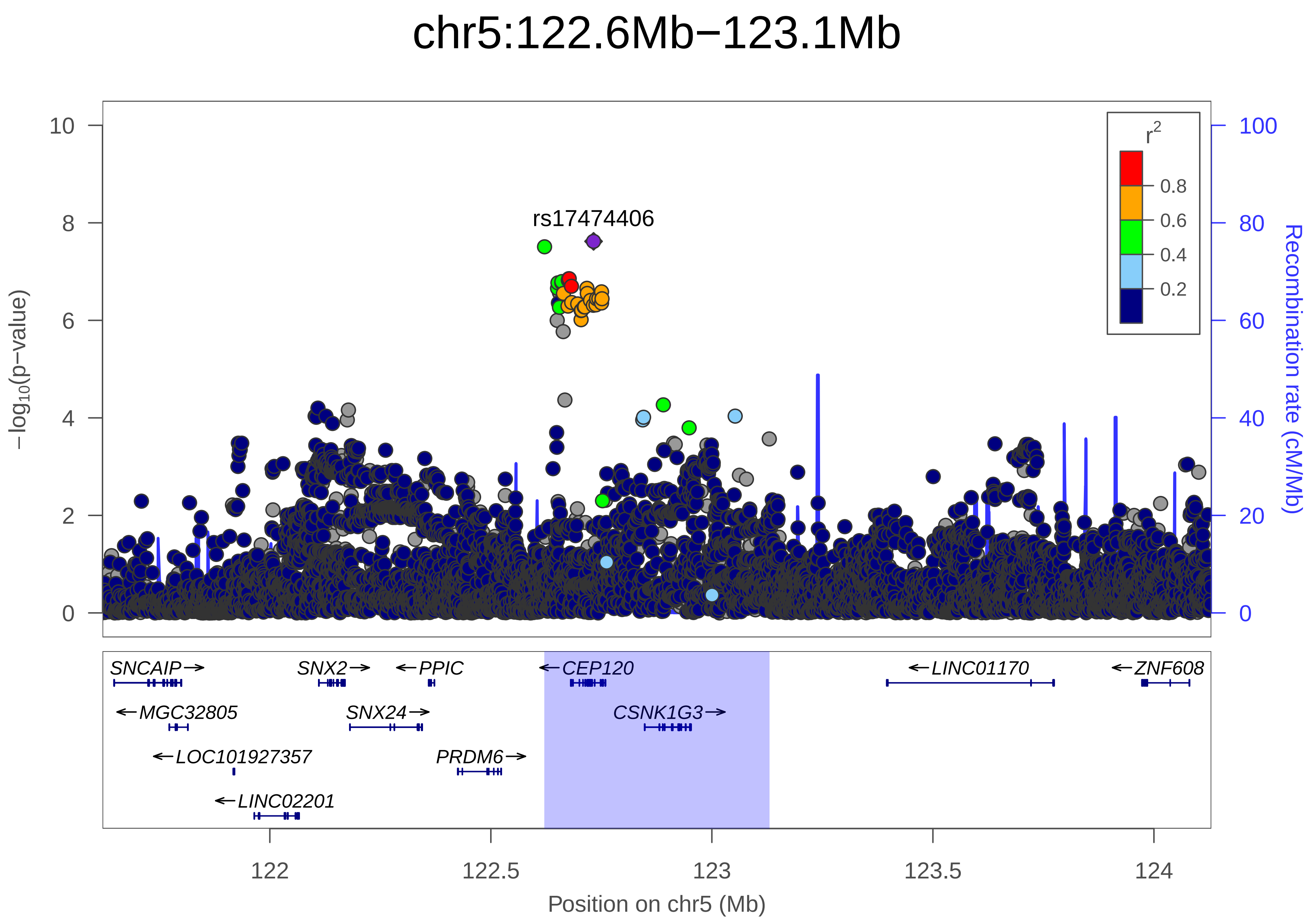
chr5:122.6Mb−123.1Mb
rs17474406
−log10(p−value)
Recombination rate (cM/Mb)
r2
0.8
0.6
0.4
0.2
SNCAIP
MGC32805
LOC101927357
LINC02201
SNX2
SNX24
PPIC
PRDM6
CEP120
CSNK1G3
LINC01170
ZNF608
Position on chr5 (Mb)

| | |
|---|---|
| date: | Mon Dec 10 08:03:19 2018 |
| build: | hg19 |
| display range: | chr5:121621471−124129838 [121621471−124129838] |
| hilite range: | 122.621Mb − 123.13Mb [ 122621000 − 123130000 ] |
| reference SNP: | chr5:122732342 |
| number of SNPs plotted: | 8836 |
| min P−value: | 2.4E−8 [chr5:122732342] |
| max P−value: | 1E0 [chr5:121649096] |

Make more plots at http://csg.sph.umich.edu/locuszoom/
