## Supplementary Figure 20 for "Genome-wide Association Study of Multisite Chronic Pain in UK Biobank"

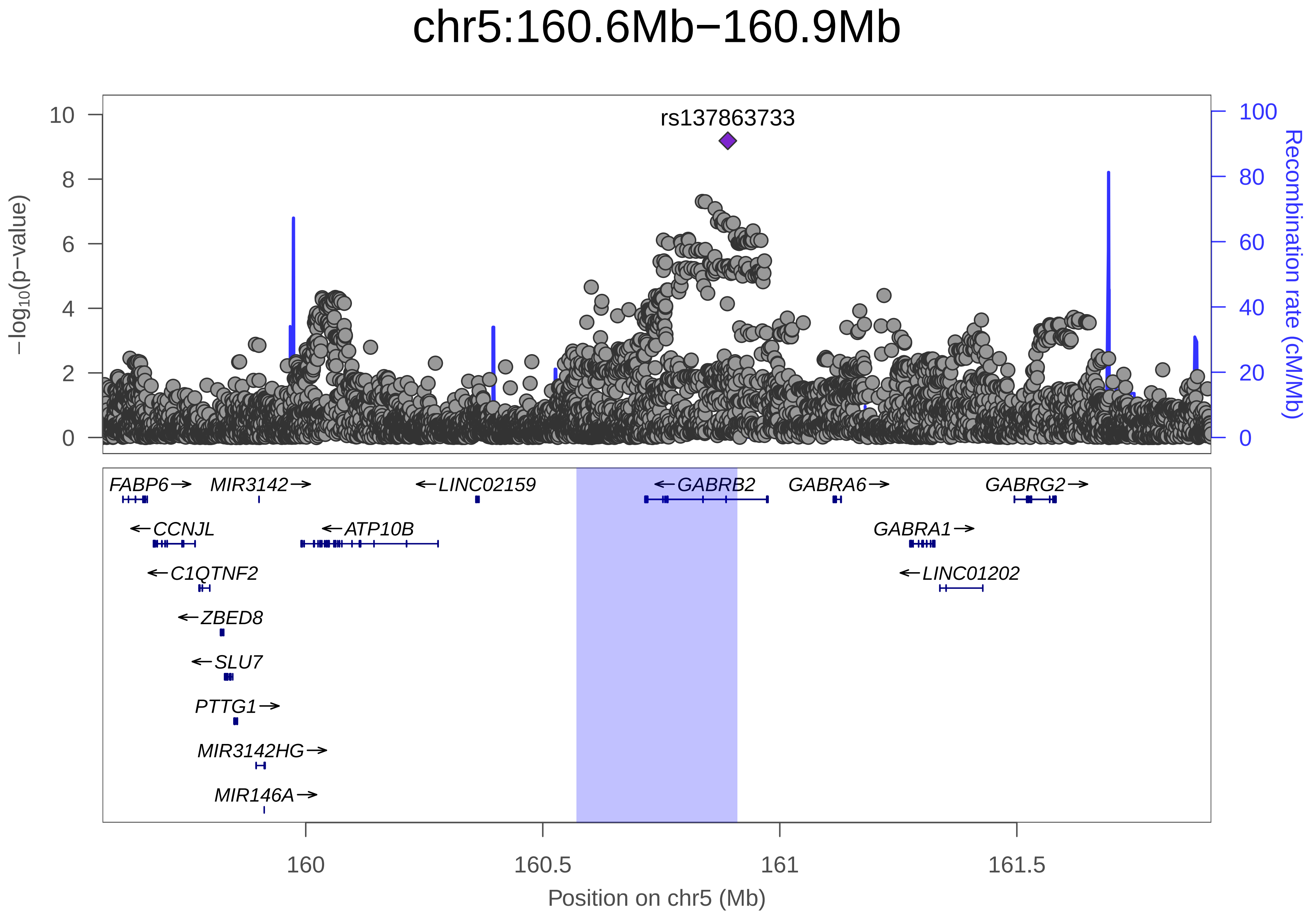
chr5:160.6Mb−160.9Mb
rs137863733
−log10(p−value)
Recombination rate (cM/Mb)
10
8
6
4
2
0
100
80
60
40
20
0
FABP6→
MIR3142→
←LINC02159
←GABRB2
GABRA6→
GABRG2→
←CCNJL
←ATP10B
GABRA1→
←C1QTNF2
←LINC01202
←ZBED8
←SLU7
PTTG1→
MIR3142HG→
MIR146A→
160
160.5
161
161.5
Position on chr5 (Mb)

| | |
|---|---|
| date: | Mon Dec 10 07:05:43 2018 |
| build: | hg19 |
| display range: | chr5:159571301−161910212 [159571301−161910212] |
| hilite range: | 160.571Mb − 160.91Mb [ 160571000 − 160910000 ] |
| reference SNP: | chr5:160890323 |
| number of SNPs plotted: | 8543 |
| min P−value: | 6.5E−10 [chr5:160890323] |
| max P−value: | 1E0 [chr5:159661723] |
| Warning: | No usable LD information for reference SNP. |

Make more plots at http://csg.sph.umich.edu/locuszoom/
