## Supplementary Figure 21 for "Genome-wide Association Study of Multisite Chronic Pain in UK Biobank"

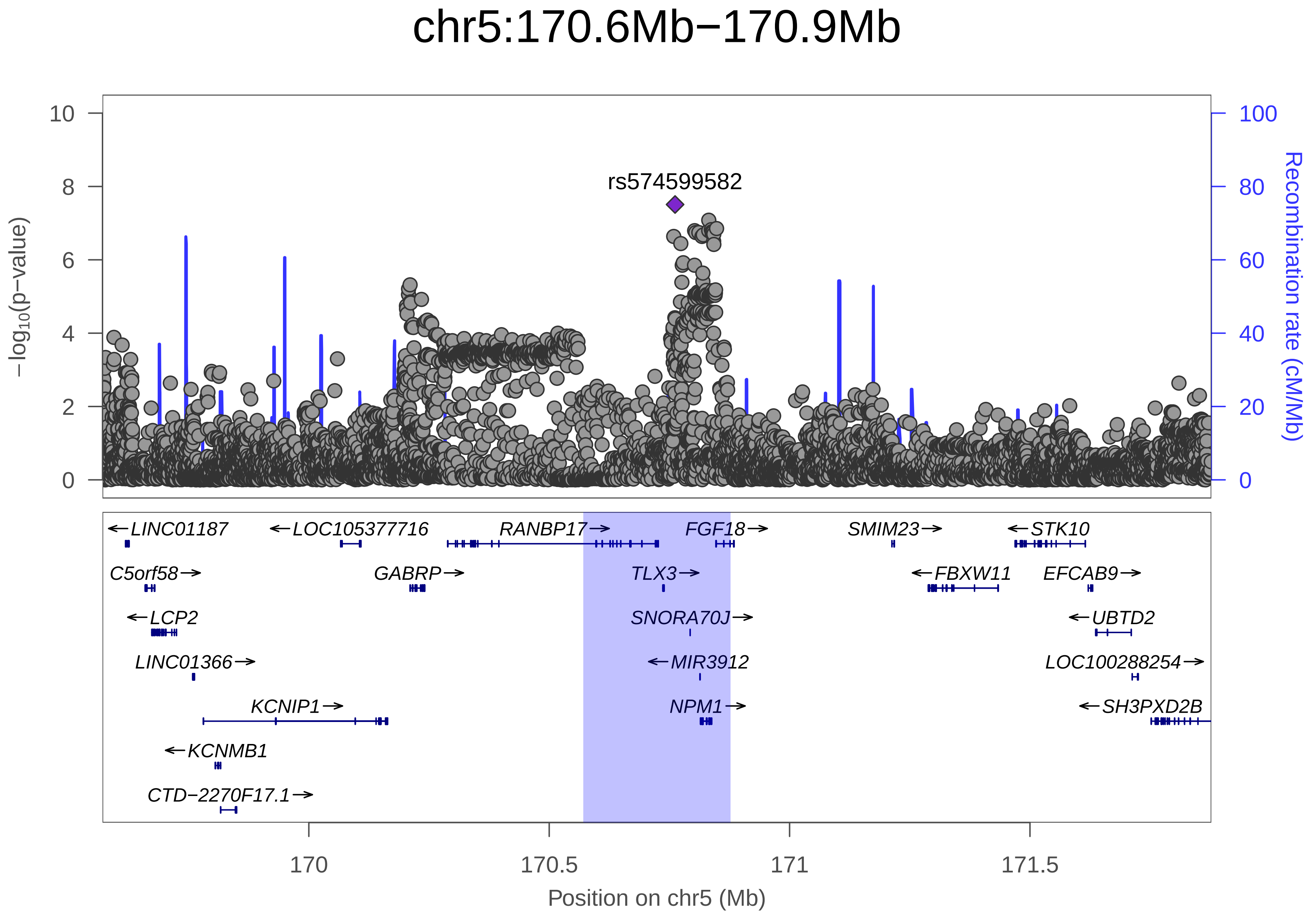

chr5:170.6Mb−170.9Mb
rs574599582
−log10(p−value)
Recombination rate (cM/Mb)
0
2
4
6
8
10
0
20
40
60
80
100
←LINC01187
←LOC105377716
RANBP17→
FGF18→
SMIM23→
←STK10
C5orf58→
GABRP→
TLX3→
←FBXW11
EFCAB9→
←LCP2
SNORA70J→
←UBTD2
LINC01366→
←MIR3912
LOC100288254→
KCNIP1→
NPM1→
←SH3PXD2B
←KCNMB1
CTD−2270F17.1→
170
170.5
171
171.5
Position on chr5 (Mb)

| | |
|---|---|
| date: | Mon Dec 10 07:16:53 2018 |
| build: | hg19 |
| display range: | chr5:169570818−171877246 [169570818−171877246] |
| hilite range: | 170.571Mb − 170.877Mb [ 170571000 − 170877000 ] |
| reference SNP: | chr5:170761810 |
| number of SNPs plotted: | 8230 |
| min P−value: | 3.1E−8 [chr5:170761810] |
| max P−value: | 1E0 [chr5:169576518] |
| Warning: | No usable LD information for reference SNP. |

Make more plots at http://csg.sph.umich.edu/locuszoom/
