## Supplementary Figure 22 for "Genome-wide Association Study of Multisite Chronic Pain in UK Biobank"

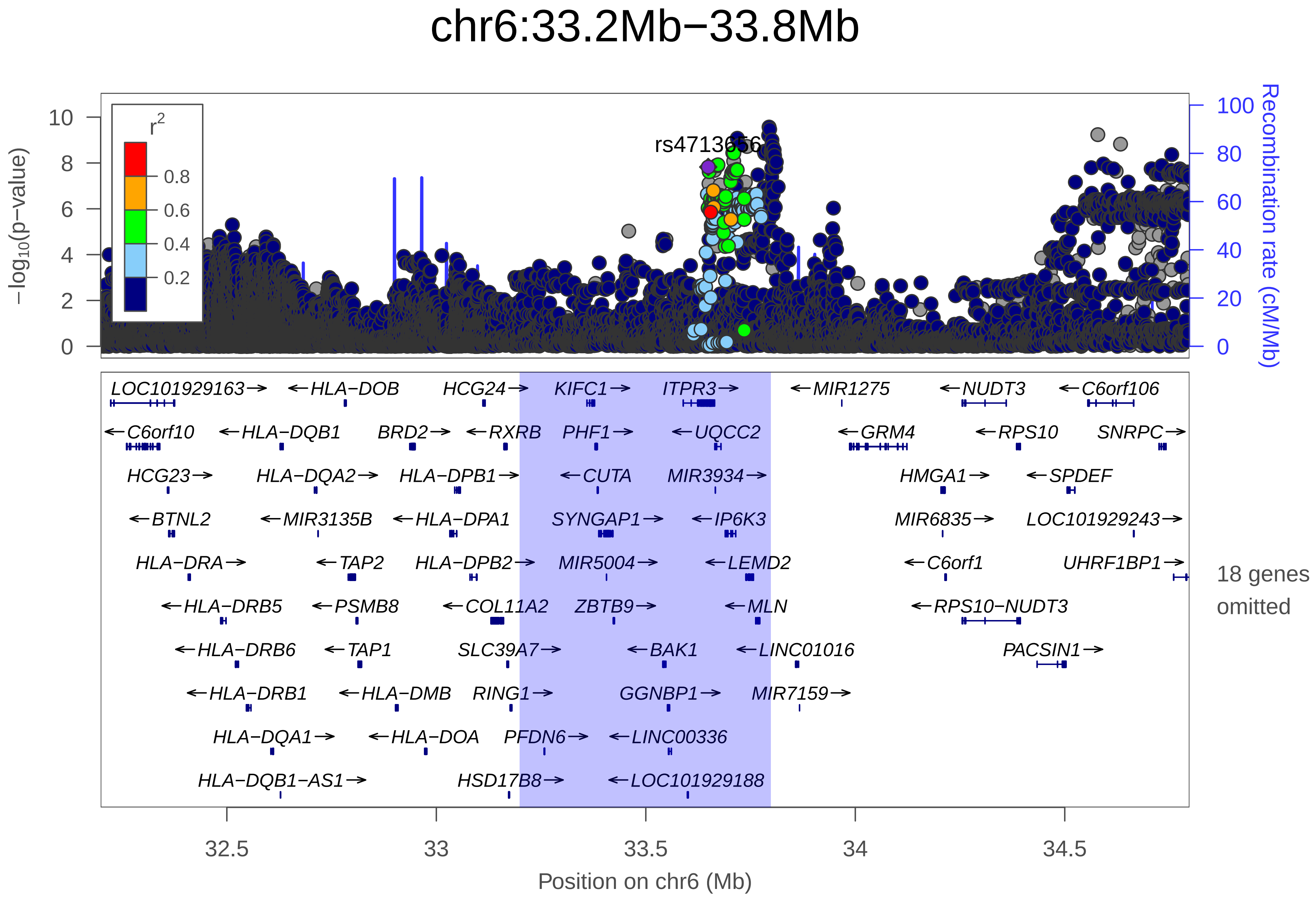
chr6:33.2Mb−33.8Mb
r2
0.8
0.6
0.4
0.2
rs4713656
−log10(p−value)
0
2
4
6
8
10
Recombination rate (cM/Mb)
0
20
40
60
80
100
LOC101929163→
←HLA−DOB
HCG24→
KIFC1→
ITPR3→
←MIR1275
←NUDT3
←C6orf106
←C6orf10
←HLA−DQB1
BRD2→
←RXRB
PHF1→
←UQCC2
←GRM4
←RPS10
SNRPC→
HCG23→
HLA−DQA2→
HLA−DPB1→
←CUTA
MIR3934→
HMGA1→
←SPDEF
←BTNL2
←MIR3135B
←HLA−DPA1
SYNGAP1→
←IP6K3
MIR6835→
LOC101929243→
HLA−DRA→
←TAP2
HLA−DPB2→
MIR5004→
←LEMD2
←C6orf1
UHRF1BP1→
←HLA−DRB5
←PSMB8
←COL11A2
ZBTB9→
←MLN
←RPS10−NUDT3
←HLA−DRB6
←TAP1
SLC39A7→
←BAK1
←LINC01016
PACSIN1→
←HLA−DRB1
←HLA−DMB
RING1→
GGNBP1→
MIR7159→
HLA−DQA1→
←HLA−DOA
PFDN6→
←LINC00336
HLA−DQB1−AS1→
HSD17B8→
←LOC101929188
18 genes omitted
32.5
33
33.5
34
34.5
Position on chr6 (Mb)

| | |
|---|---|
| date: | Mon Dec 10 07:55:15 2018 |
| build: | hg19 |
| display range: | chr6:32198771−34797628 [32198771−34797628] |
| hilite range: | 33.199Mb − 33.798Mb [ 33199000 − 33798000 ] |
| reference SNP: | chr6:33648997 |
| number of SNPs plotted: | 30845 |
| min P−value: | 2.7E−10 [chr6:33794215] |
| max P−value: | 1E0 [chr6:32217955] |
| omitted Genes: | HLA−DQB2, PSMB8−AS1, PSMB9 |
| omitted Genes: | LOC100294145, HLA−DMA, MIR219A1 |
| omitted Genes: | HCG25, VPS52, RPS18 |
| omitted Genes: | B3GALT4, WDR46, MIR6873 |
| omitted Genes: | MIR6834, RGL2, TAPBP |
| omitted Genes: | ZBTB22, MIR1234, DAXX |

Make more plots at http://csg.sph.umich.edu/locuszoom/
