## Supplementary Figure 23 for "Genome-wide Association Study of Multisite Chronic Pain in UK Biobank"

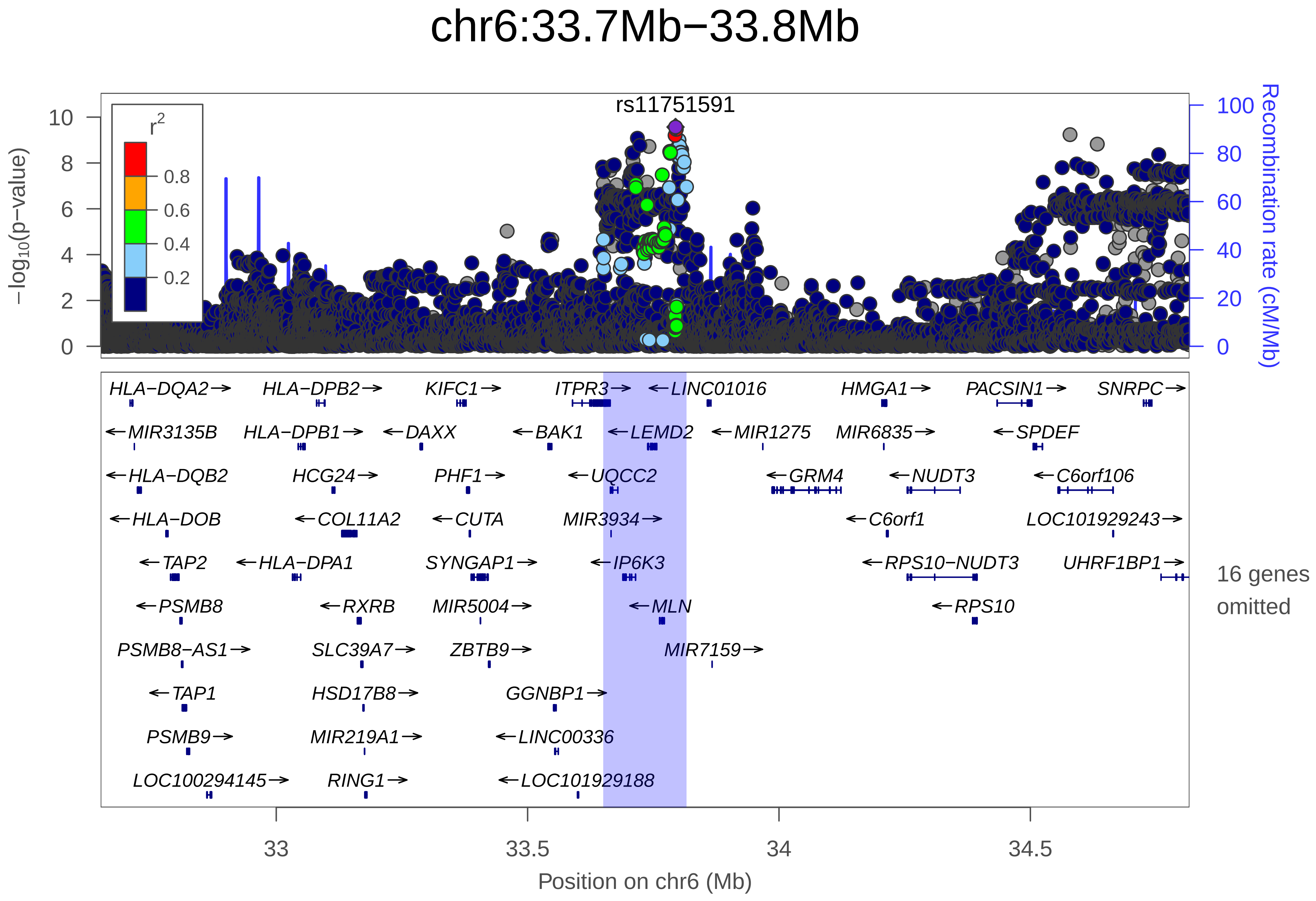
chr6:33.7Mb−33.8Mb
rs11751591
r2
0.8
0.6
0.4
0.2
−log10(p−value)
Recombination rate (cM/Mb)
HLA−DQA2→
HLA−DPB2→
KIFC1→
ITPR3→
←LINC01016
HMGA1→
PACSIN1→
SNRPC→
←MIR3135B
HLA−DPB1→
←DAXX
←BAK1
←LEMD2
←MIR1275
MIR6835→
←SPDEF
←HLA−DQB2
HCG24→
PHF1→
←UQCC2
←GRM4
←NUDT3
←C6orf106
←HLA−DOB
←COL11A2
←CUTA
MIR3934→
←C6orf1
LOC101929243→
←TAP2
←HLA−DPA1
SYNGAP1→
←IP6K3
←RPS10−NUDT3
UHRF1BP1→
16 genes omitted
←PSMB8
←RXRB
MIR5004→
←MLN
←RPS10
PSMB8−AS1→
SLC39A7→
ZBTB9→
MIR7159→
←TAP1
HSD17B8→
GGNBP1→
PSMB9→
MIR219A1→
←LINC00336
LOC100294145→
RING1→
←LOC101929188
33
33.5
34
34.5
Position on chr6 (Mb)

| | |
|---|---|
| date: | Mon Dec 10 08:23:45 2018 |
| build: | hg19 |
| display range: | chr6:32650743−34816451 [32650743−34816451] |
| hilite range: | 33.651Mb − 33.816Mb [ 33651000 − 33816000 ] |
| reference SNP: | chr6:33794215 |
| number of SNPs plotted: | 14696 |
| min P−value: | 2.7E−10 [chr6:33794215] |
| max P−value: | 1E0 [chr6:32657351] |
| omitted Genes: | HLA−DMB, HLA−DMA, BRD2 |
| omitted Genes: | HLA−DOA, HCG25, VPS52 |
| omitted Genes: | RPS18, B3GALT4, WDR46 |
| omitted Genes: | MIR6873, PFDN6, MIR6834 |
| omitted Genes: | RGL2, TAPBP, ZBTB22 |
| omitted Genes: | MIR1234 |

Make more plots at http://csg.sph.umich.edu/locuszoom/
