## Supplementary Figure 24 for "Genome-wide Association Study of Multisite Chronic Pain in UK Biobank"

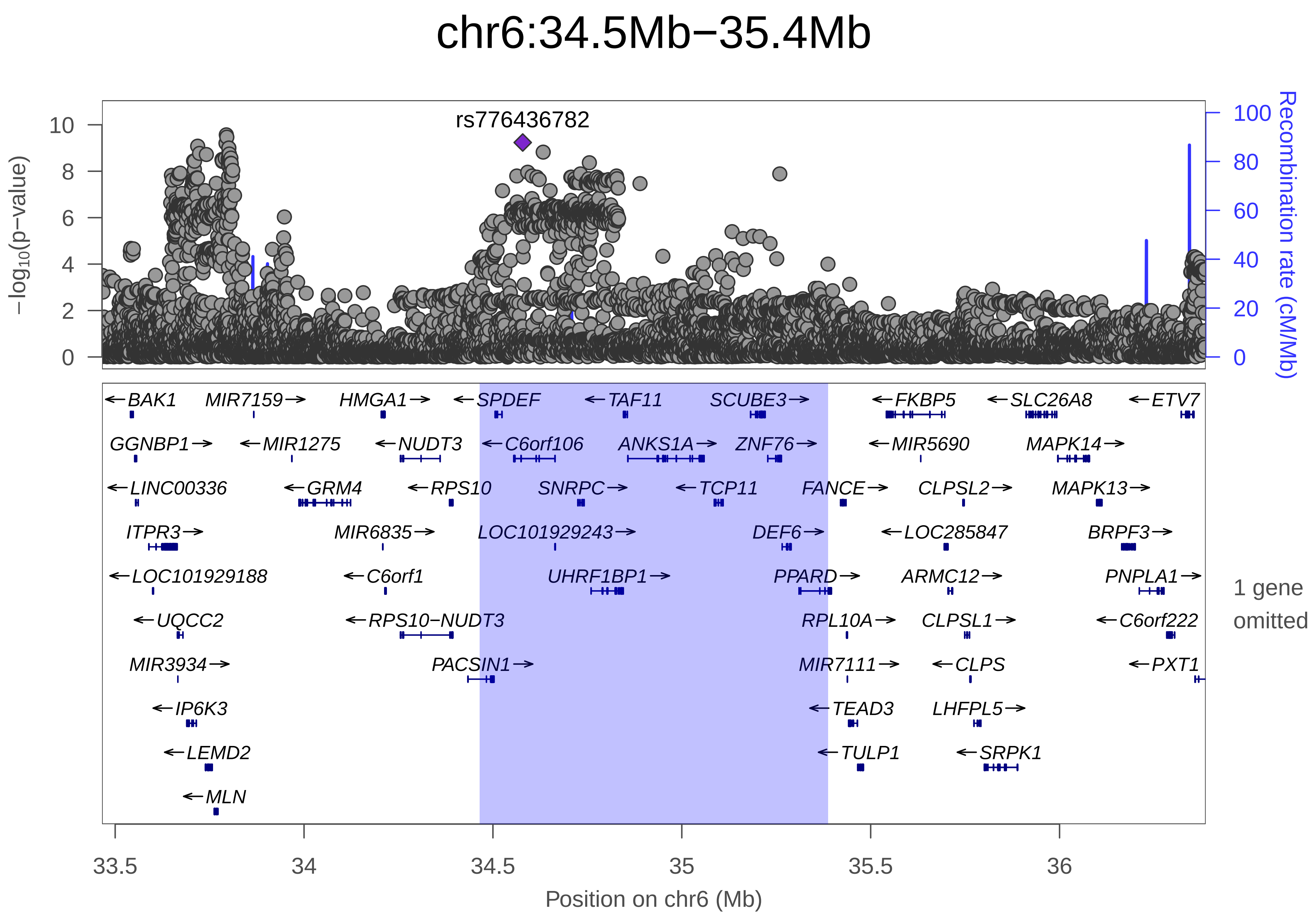
chr6:34.5Mb−35.4Mb
rs776436782
−log10(p−value)
Recombination rate (cM/Mb)
0
2
4
6
8
10
0
20
40
60
80
100
BAK1
MIR7159
HMGA1
SPDEF
TAF11
SCUBE3
FKBP5
SLC26A8
ETV7
GGNBP1
MIR1275
NUDT3
C6orf106
ANKS1A
ZNF76
MIR5690
MAPK14
LINC00336
GRM4
RPS10
SNRPC
TCP11
FANCE
CLPSL2
MAPK13
ITPR3
MIR6835
LOC101929243
DEF6
LOC285847
BRPF3
LOC101929188
C6orf1
UHRF1BP1
PPARD
ARMC12
PNPLA1
UQCC2
RPS10−NUDT3
RPL10A
CLPSL1
C6orf222
MIR3934
PACSIN1
MIR7111
CLPS
PXT1
IP6K3
TEAD3
LHFPL5
LEMD2
TULP1
SRPK1
MLN
1 gene omitted
33.5
34
34.5
35
35.5
36
Position on chr6 (Mb)

| | |
|---|---|
| date: | Mon Dec 10 07:02:39 2018 |
| build: | hg19 |
| display range: | chr6:33465215−36386872 [33465215−36386872] |
| hilite range: | 34.465Mb − 35.387Mb [ 34465000 − 35387000 ] |
| reference SNP: | chr6:34578879 |
| number of SNPs plotted: | 10776 |
| min P−value: | 2.7E−10 [chr6:33794215] |
| max P−value: | 1E0 [chr6:33565474] |
| omitted Genes: | LINC01016 |
| Warning: | No usable LD information for reference SNP. |

Make more plots at http://csg.sph.umich.edu/locuszoom/
