## Supplementary Figure 25 for "Genome-wide Association Study of Multisite Chronic Pain in UK Biobank"

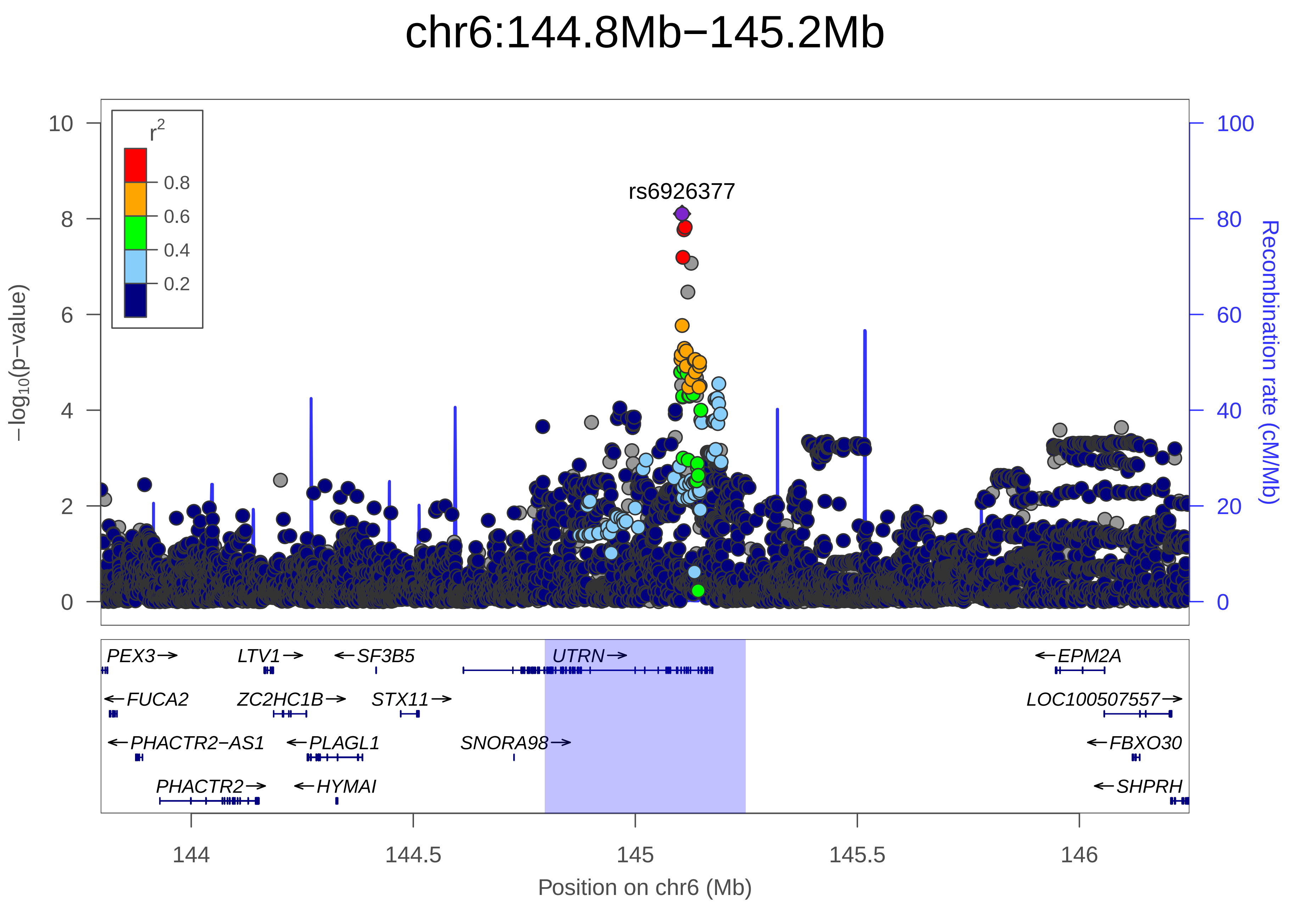

chr6:144.8Mb−145.2Mb
rs6926377
r2
0.8
0.6
0.4
0.2
−log10(p−value)
Recombination rate (cM/Mb)
PEX3
LTV1
SF3B5
UTRN
EPM2A
FUCA2
ZC2HC1B
STX11
LOC100507557
PHACTR2−AS1
PLAGL1
SNORA98
FBXO30
PHACTR2
HYMAI
SHPRH
144
144.5
145
145.5
146
Position on chr6 (Mb)

| | |
|---|---|
| date: | Mon Dec 10 07:16:54 2018 |
| build: | hg19 |
| display range: | chr6:143796045−146247885 [143796045−146247885] |
| hilite range: | 144.796Mb − 145.248Mb [ 144796000 − 145248000 ] |
| reference SNP: | chr6:145105354 |
| number of SNPs plotted: | 7812 |
| min P−value: | 7.9E−9 [chr6:145105354] |
| max P−value: | 1E0 [chr6:143918183] |

Make more plots at http://csg.sph.umich.edu/locuszoom/
