## Supplementary Figure 26 for "Genome-wide Association Study of Multisite Chronic Pain in UK Biobank"

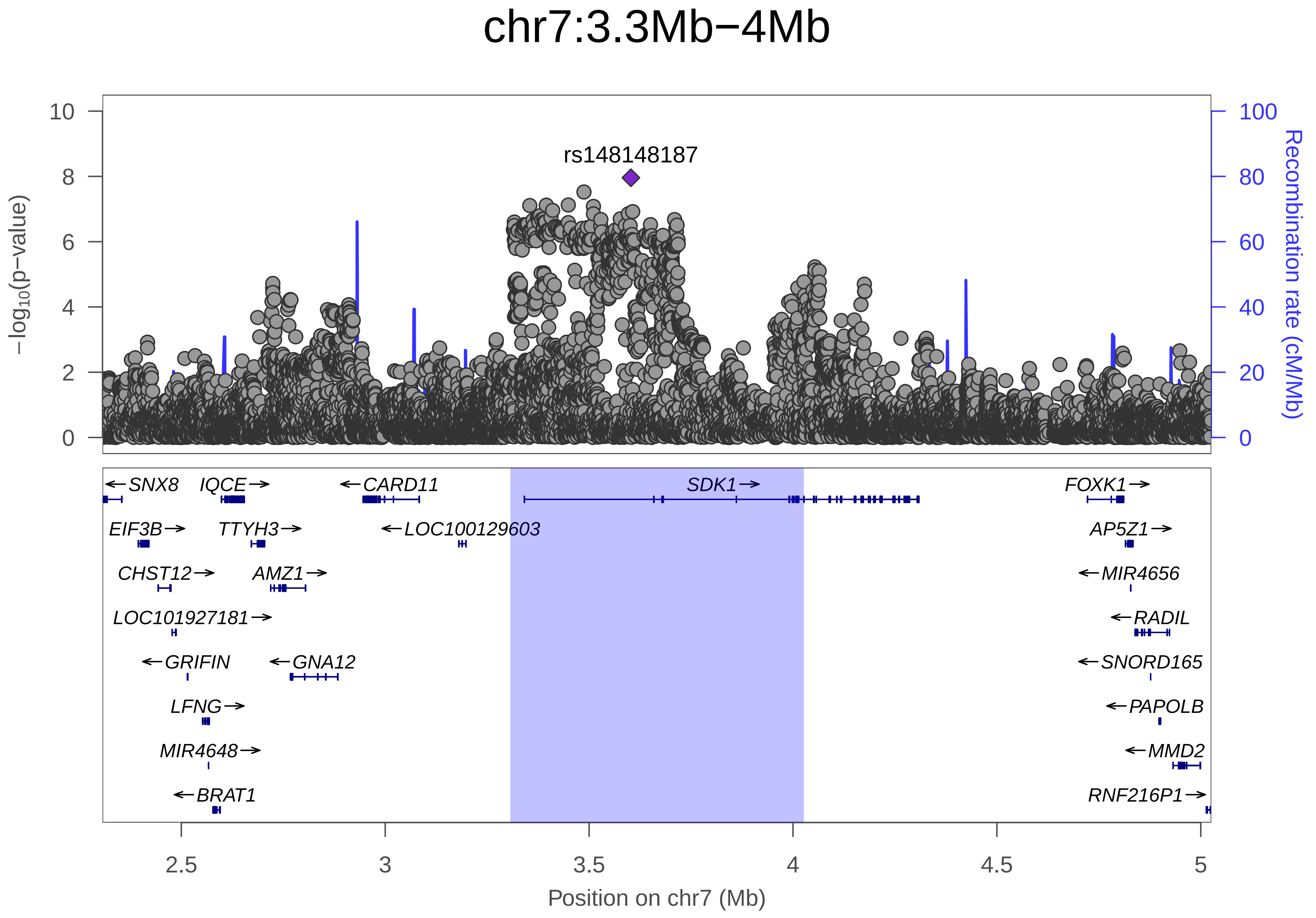

chr7:3.3Mb−4Mb
rs148148187
−log10(p−value)
Recombination rate (cM/Mb)
0
2
4
6
8
10
0
20
40
60
80
100
SNX8
IQCE
CARD11
SDK1
FOXK1
EIF3B
TTYH3
LOC100129603
AP5Z1
CHST12
AMZ1
MIR4656
LOC101927181
RADIL
GRIFIN
GNA12
SNORD165
LFNG
PAPOLB
MIR4648
MMD2
BRAT1
RNF216P1
2.5
3
3.5
4
4.5
5
Position on chr7 (Mb)

| | |
|---|---|
| date: | Mon Dec 10 07:55:13 2018 |
| build: | hg19 |
| display range: | chr7:2306708−5025830 [2306708−5025830] |
| hilite range: | 3.307Mb − 4.026Mb [ 3307000 − 4026000 ] |
| reference SNP: | chr7:3602520 |
| number of SNPs plotted: | 13754 |
| min P−value: | 1.1E−8 [chr7:3602520] |
| max P−value: | 1E0 [chr7:2339405] |
| Warning: | No usable LD information for reference SNP. |

Make more plots at http://csg.sph.umich.edu/locuszoom/
