## Supplementary Figure 27 for "Genome-wide Association Study of Multisite Chronic Pain in UK Biobank"

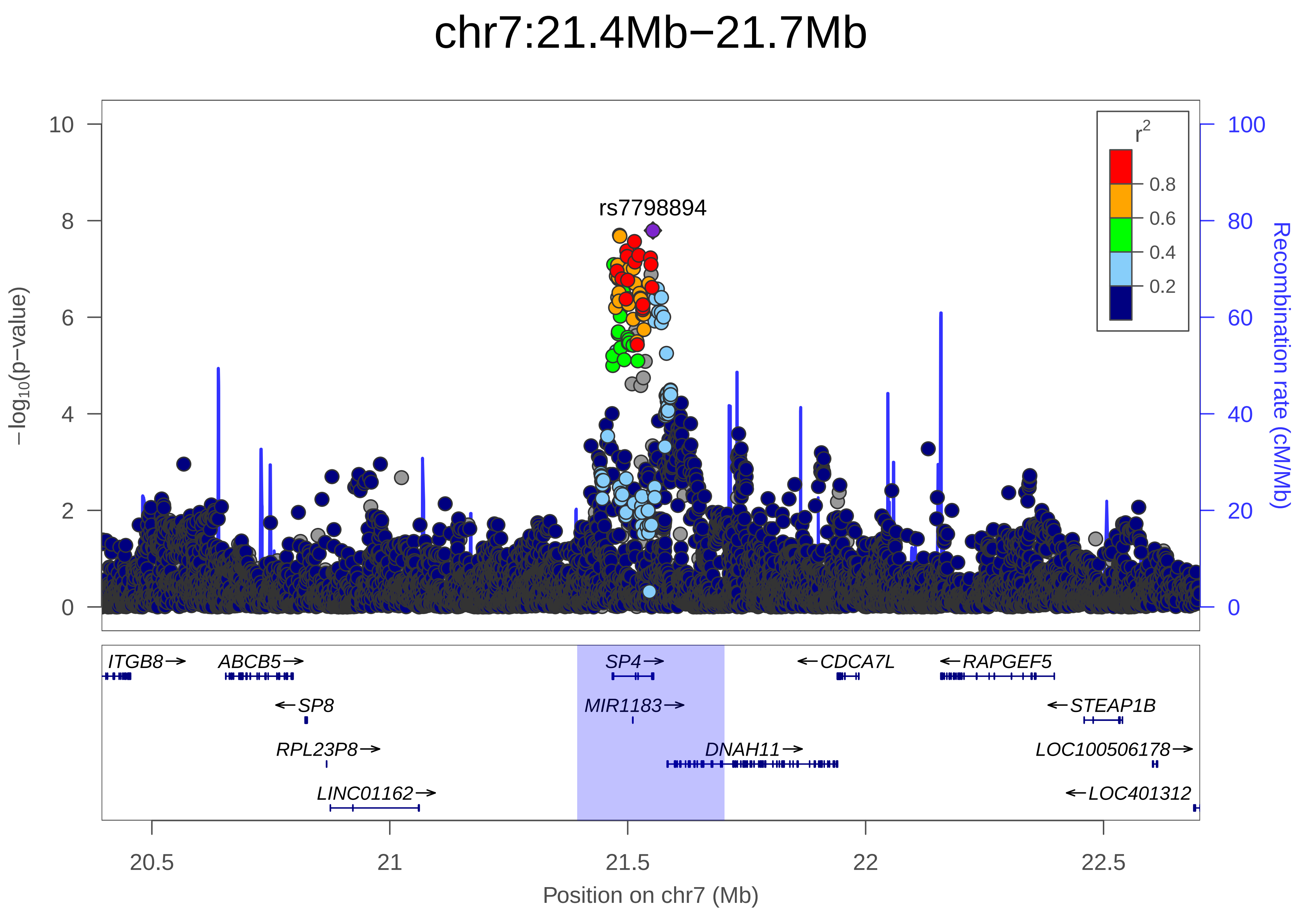

chr7:21.4Mb−21.7Mb
rs7798894
r2
0.8
0.6
0.4
0.2
10
8
6
4
2
0
100
80
60
40
20
0
−log10(p−value)
Recombination rate (cM/Mb)
ITGB8→
ABCB5→
SP4→
←CDCA7L
←RAPGEF5
←SP8
MIR1183→
←STEAP1B
RPL23P8→
DNAH11→
LOC100506178→
LINC01162→
←LOC401312
20.5
21
21.5
22
22.5
Position on chr7 (Mb)

| | |
|---|---|
| date: | Mon Dec 10 08:18:45 2018 |
| build: | hg19 |
| display range: | chr7:20394246−22702980 [20394246−22702980] |
| hilite range: | 21.394Mb − 21.703Mb [ 21394000 − 21703000 ] |
| reference SNP: | chr7:21552995 |
| number of SNPs plotted: | 9738 |
| min P−value: | 1.6E−8 [chr7:21552995] |
| max P−value: | 1E0 [chr7:20394485] |

Make more plots at http://csg.sph.umich.edu/locuszoom/
