## Supplementary Figure 28 for "Genome-wide Association Study of Multisite Chronic Pain in UK Biobank"

### chr7:95.6Mb−96.1Mb

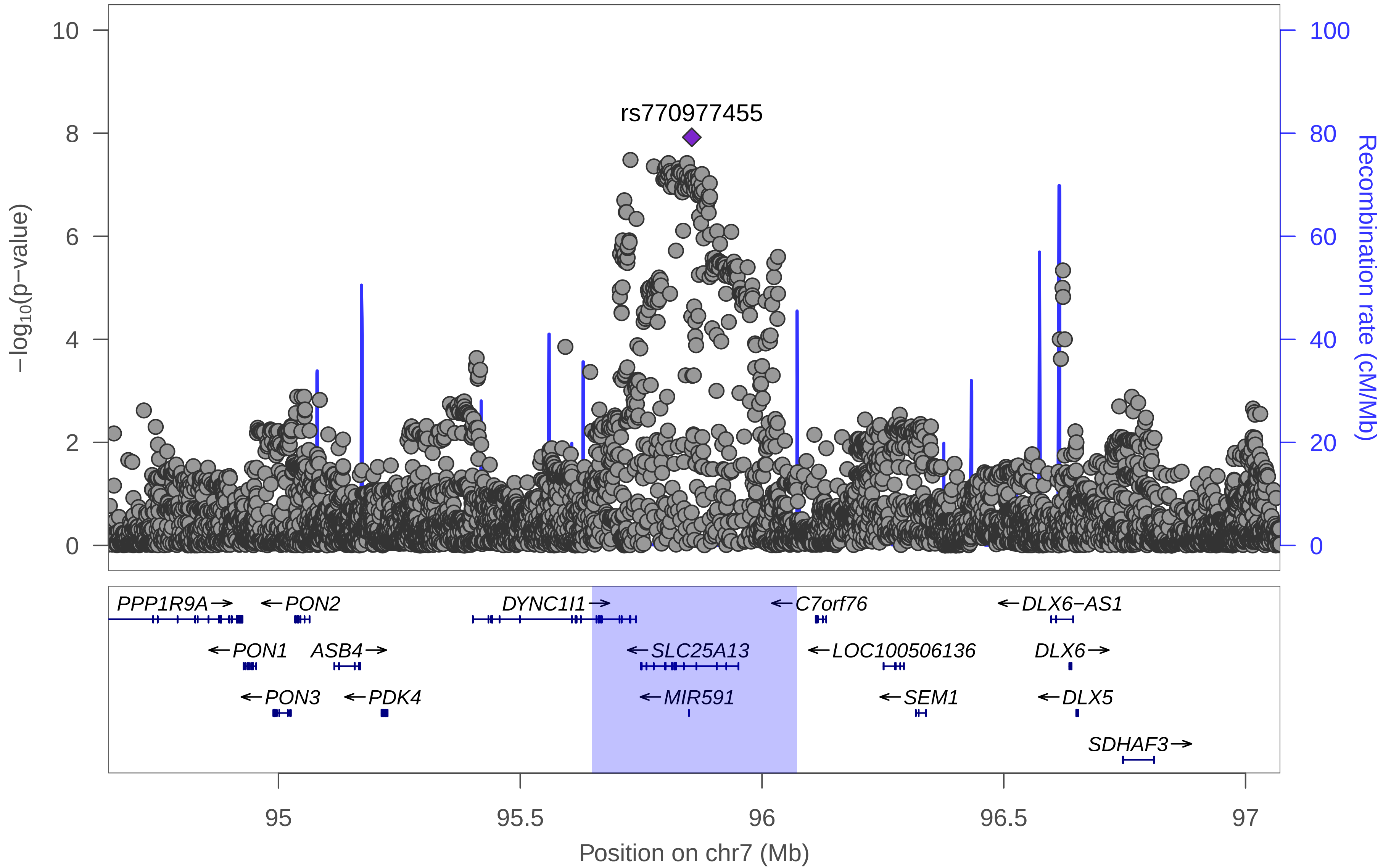

| | |
|---|---|
| date: | Mon Dec 10 08:41:19 2018 |
| build: | hg19 |
| display range: | chr7:94648166−97072003 [94648166−97072003] |
| hilite range: | 95.648Mb − 96.072Mb [ 95648000 − 96072000 ] |
| reference SNP: | chr7:95854845 |
| number of SNPs plotted: | 7304 |
| min P−value: | 1.2E−8 [chr7:95854845] |
| max P−value: | 1E0 [chr7:94812564] |
| Warning: | No usable LD information for reference SNP. |

Make more plots at http://csg.sph.umich.edu/locuszoom/
