## Supplementary Figure 29 for "Genome-wide Association Study of Multisite Chronic Pain in UK Biobank"

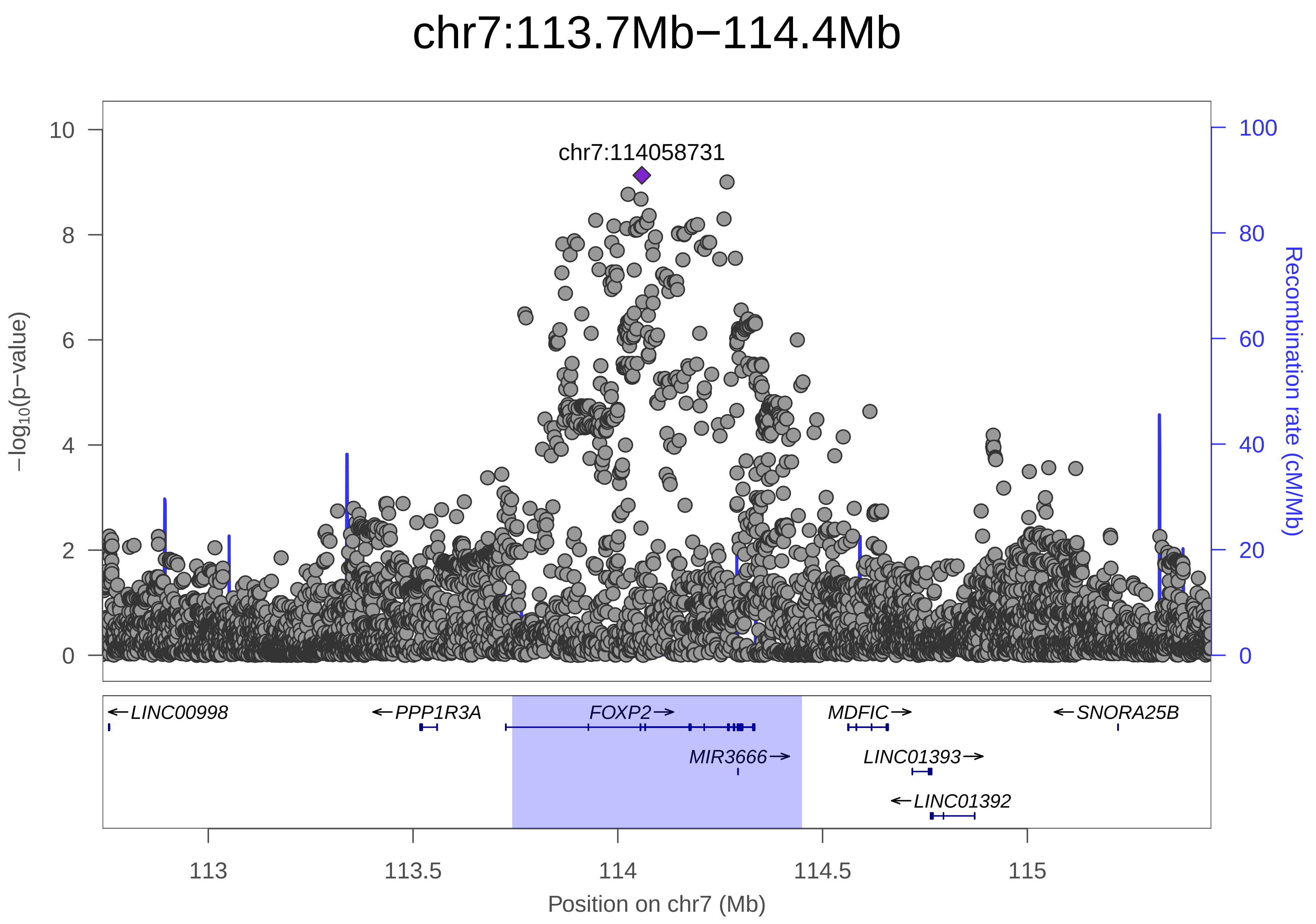
chr7:113.7Mb−114.4Mb
chr7:114058731
−log10(p−value)
Recombination rate (cM/Mb)
0
2
4
6
8
10
0
20
40
60
80
100
LINC00998
PPP1R3A
FOXP2
MDFIC
SNORA25B
MIR3666
LINC01393
LINC01392
113
113.5
114
114.5
115
Position on chr7 (Mb)

| | |
|---|---|
| date: | Mon Dec 10 07:02:39 2018 |
| build: | hg19 |
| display range: | chr7:112741727−115449185 [112741727−115449185] |
| hilite range: | 113.742Mb − 114.449Mb [ 113742000 − 114449000 ] |
| reference SNP: | chr7:114058731 |
| number of SNPs plotted: | 7233 |
| min P−value: | 7.4E−10 [chr7:114058731] |
| max P−value: | 1E0 [chr7:112971737] |
| Warning: | No usable LD information for reference SNP. |

Make more plots at http://csg.sph.umich.edu/locuszoom/
