## Supplementary Figure 30 for "Genome-wide Association Study of Multisite Chronic Pain in UK Biobank"

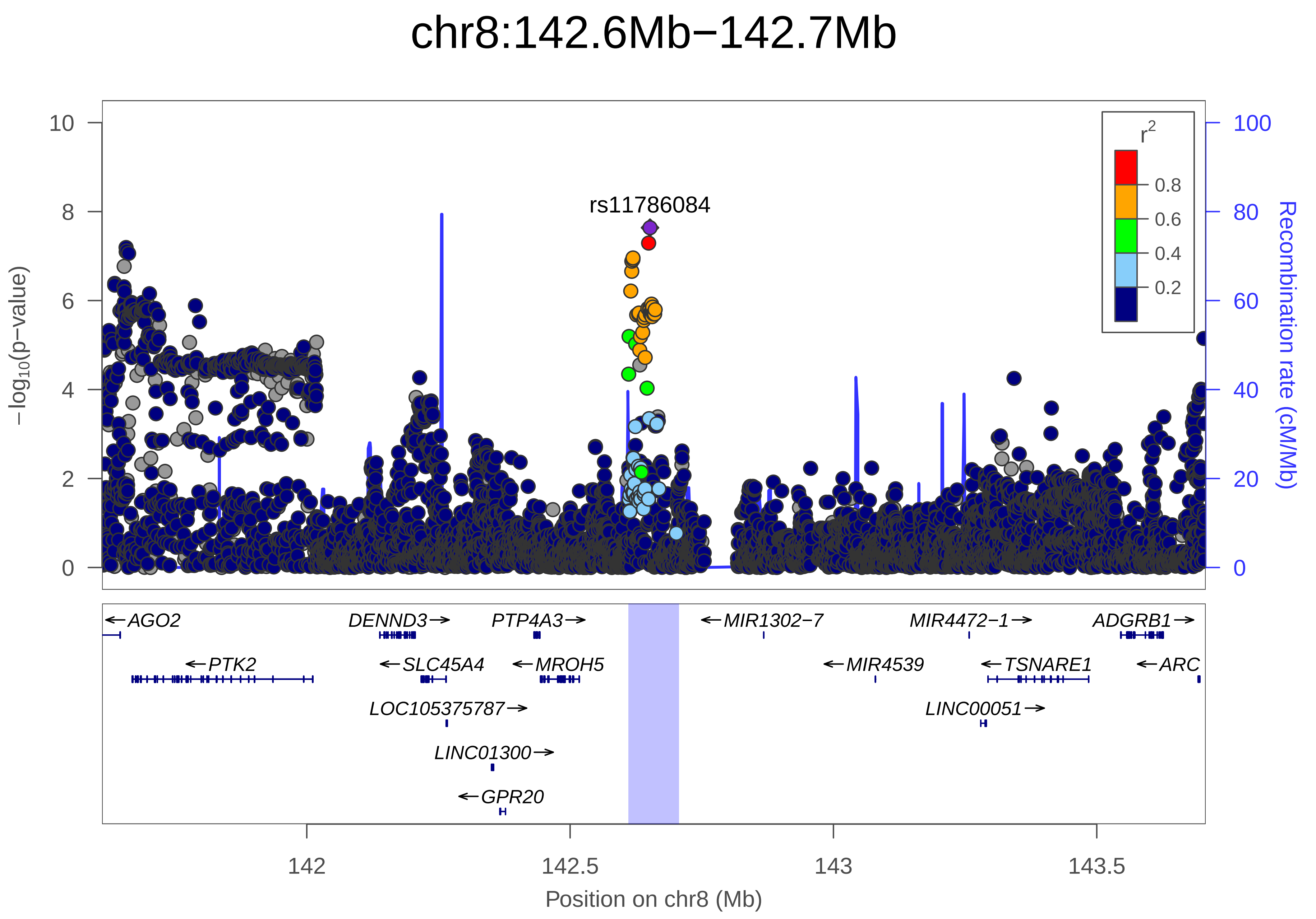
chr8:142.6Mb−142.7Mb
−log10(p−value)
Recombination rate (cM/Mb)
rs11786084
r2
0.8
0.6
0.4
0.2
AGO2
PTK2
DENND3
SLC45A4
LOC105375787
LINC01300
GPR20
PTP4A3
MROH5
MIR1302−7
MIR4539
MIR4472−1
TSNARE1
LINC00051
ADGRB1
ARC
142
142.5
143
143.5
Position on chr8 (Mb)

| | |
|---|---|
| date: | Mon Dec 10 07:16:54 2018 |
| build: | hg19 |
| display range: | chr8:141611220−143706843 [141611220−143706843] |
| hilite range: | 142.611Mb − 142.707Mb [ 142611000 − 142707000 ] |
| reference SNP: | chr8:142651709 |
| number of SNPs plotted: | 7878 |
| min P−value: | 2.3E−8 [chr8:142651709] |
| max P−value: | 1E0 [chr8:141703315] |

Make more plots at http://csg.sph.umich.edu/locuszoom/
