## Supplementary Figure 31 for "Genome-wide Association Study of Multisite Chronic Pain in UK Biobank"

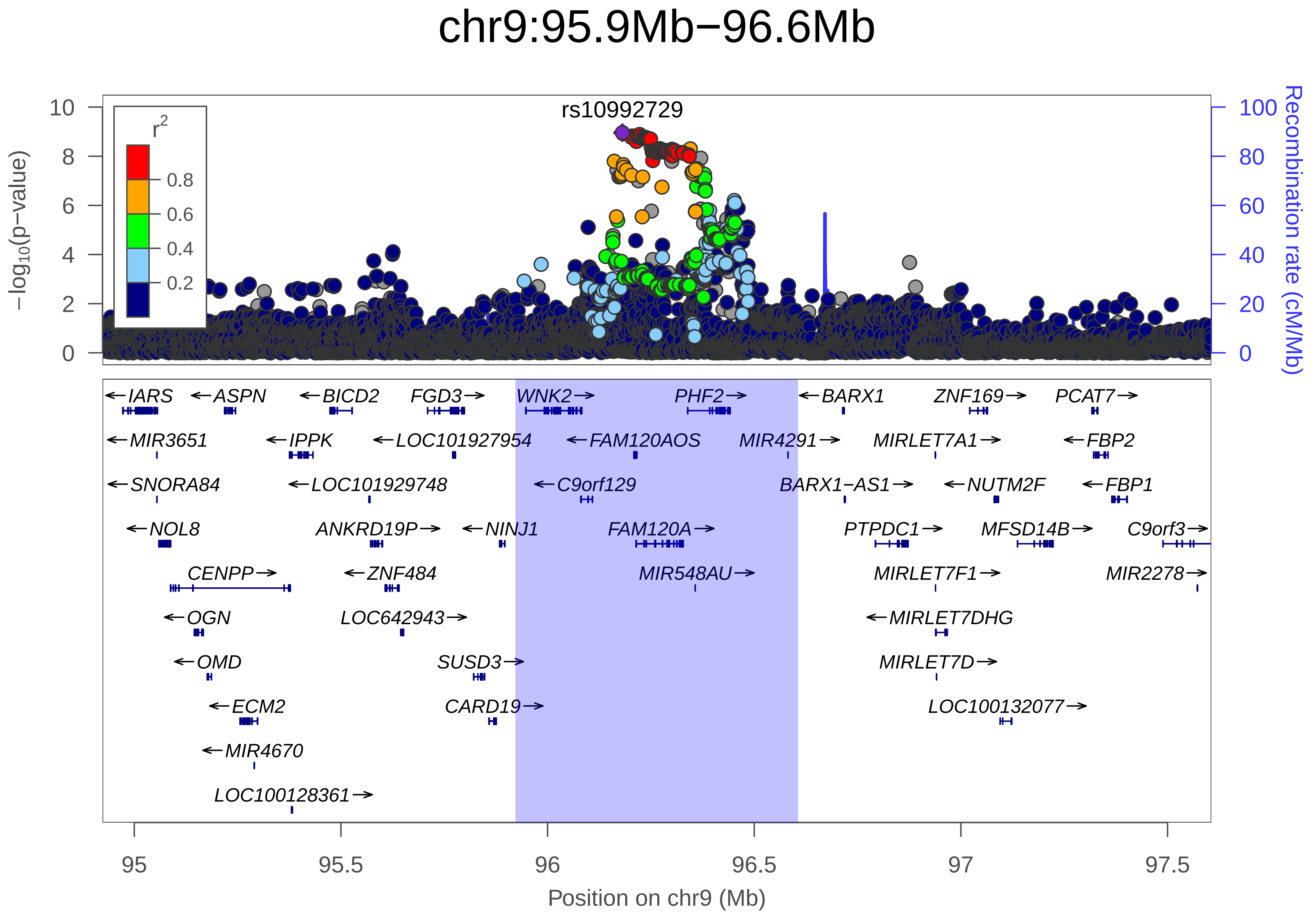

chr9:95.9Mb−96.6Mb
rs10992729
r2
0.8
0.6
0.4
0.2
−log10(p−value)
Recombination rate (cM/Mb)
IARS
ASPN
BICD2
FGD3
WNK2
PHF2
BARX1
ZNF169
PCAT7
MIR3651
IPPK
LOC101927954
FAM120AOS
MIR4291
MIRLET7A1
FBP2
SNORA84
LOC101929748
C9orf129
BARX1−AS1
NUTM2F
FBP1
NOL8
ANKRD19P
NINJ1
FAM120A
PTPDC1
MFSD14B
C9orf3
CENPP
ZNF484
MIR548AU
MIRLET7F1
MIR2278
OGN
LOC642943
MIRLET7DHG
OMD
SUSD3
MIRLET7D
ECM2
CARD19
LOC100132077
MIR4670
LOC100128361
Position on chr9 (Mb)

| | |
|---|---|
| date: | Mon Dec 10 07:41:34 2018 |
| build: | hg19 |
| display range: | chr9:94923174−97605924 [94923174−97605924] |
| hilite range: | 95.923Mb − 96.606Mb [ 95923000 − 96606000 ] |
| reference SNP: | chr9:96181075 |
| number of SNPs plotted: | 10172 |
| min P−value: | 1.1E−9 [chr9:96181075] |
| max P−value: | 1E0 [chr9:94983580] |

Make more plots at http://csg.sph.umich.edu/locuszoom/
