## Supplementary Figure 32 for "Genome-wide Association Study of Multisite Chronic Pain in UK Biobank"

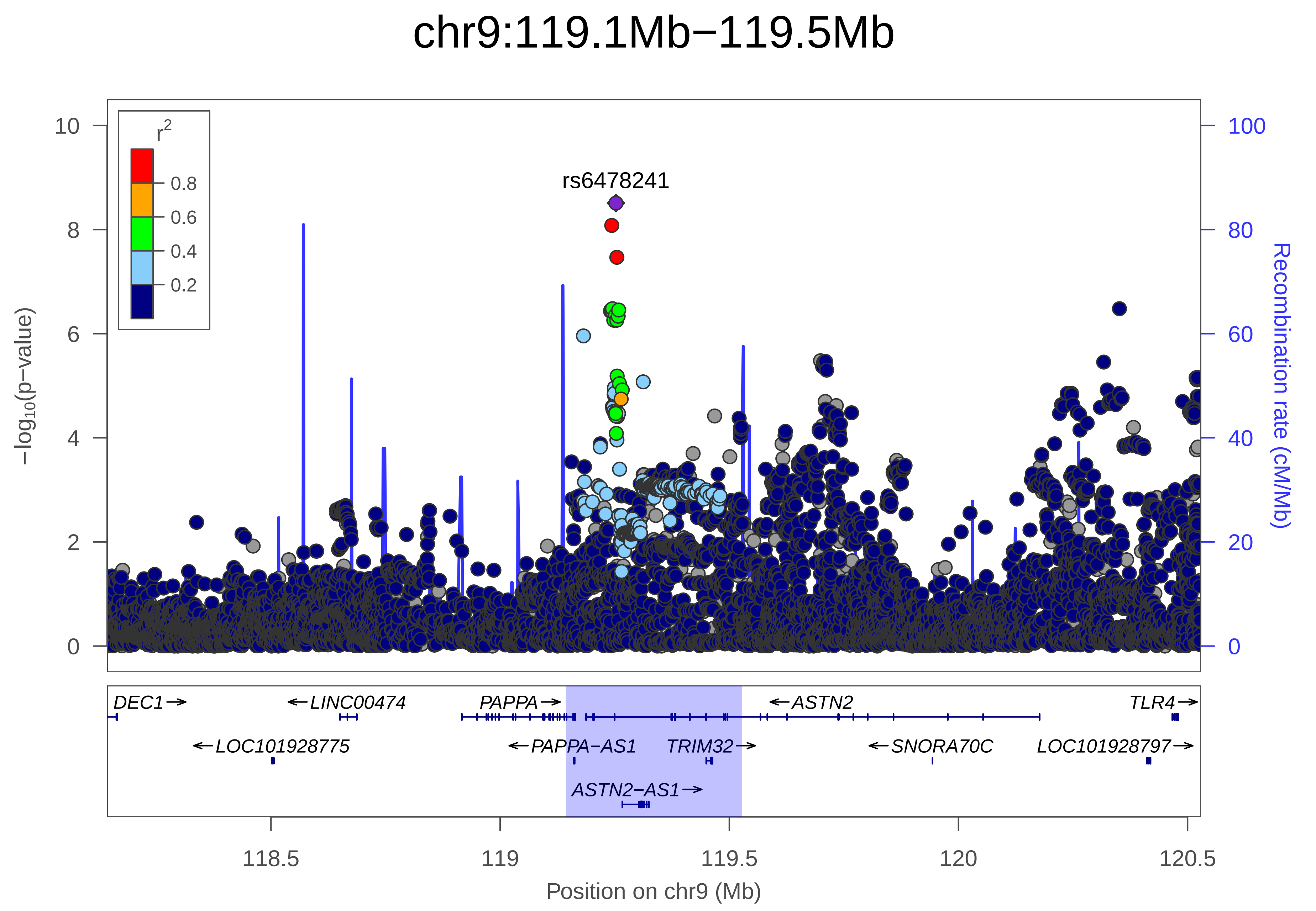

chr9:119.1Mb−119.5Mb
rs6478241
r2
0.8
0.6
0.4
0.2
−log10(p−value)
Recombination rate (cM/Mb)
DEC1→
←LINC00474
PAPPA→
←ASTN2
TLR4→
←LOC101928775
←PAPPA−AS1
TRIM32→
←SNORA70C
LOC101928797→
ASTN2−AS1→
118.5
119
119.5
120
120.5
Position on chr9 (Mb)

| | |
|---|---|
| date: | Mon Dec 10 08:03:18 2018 |
| build: | hg19 |
| display range: | chr9:118142572−120528478 [118142572−120528478] |
| hilite range: | 119.143Mb − 119.528Mb [ 119143000 − 119528000 ] |
| reference SNP: | chr9:119252629 |
| number of SNPs plotted: | 8388 |
| min P−value: | 3.1E−9 [chr9:119252629] |
| max P−value: | 1E0 [chr9:118289206] |

Make more plots at http://csg.sph.umich.edu/locuszoom/
