## Supplementary Figure 33 for "Genome-wide Association Study of Multisite Chronic Pain in UK Biobank"

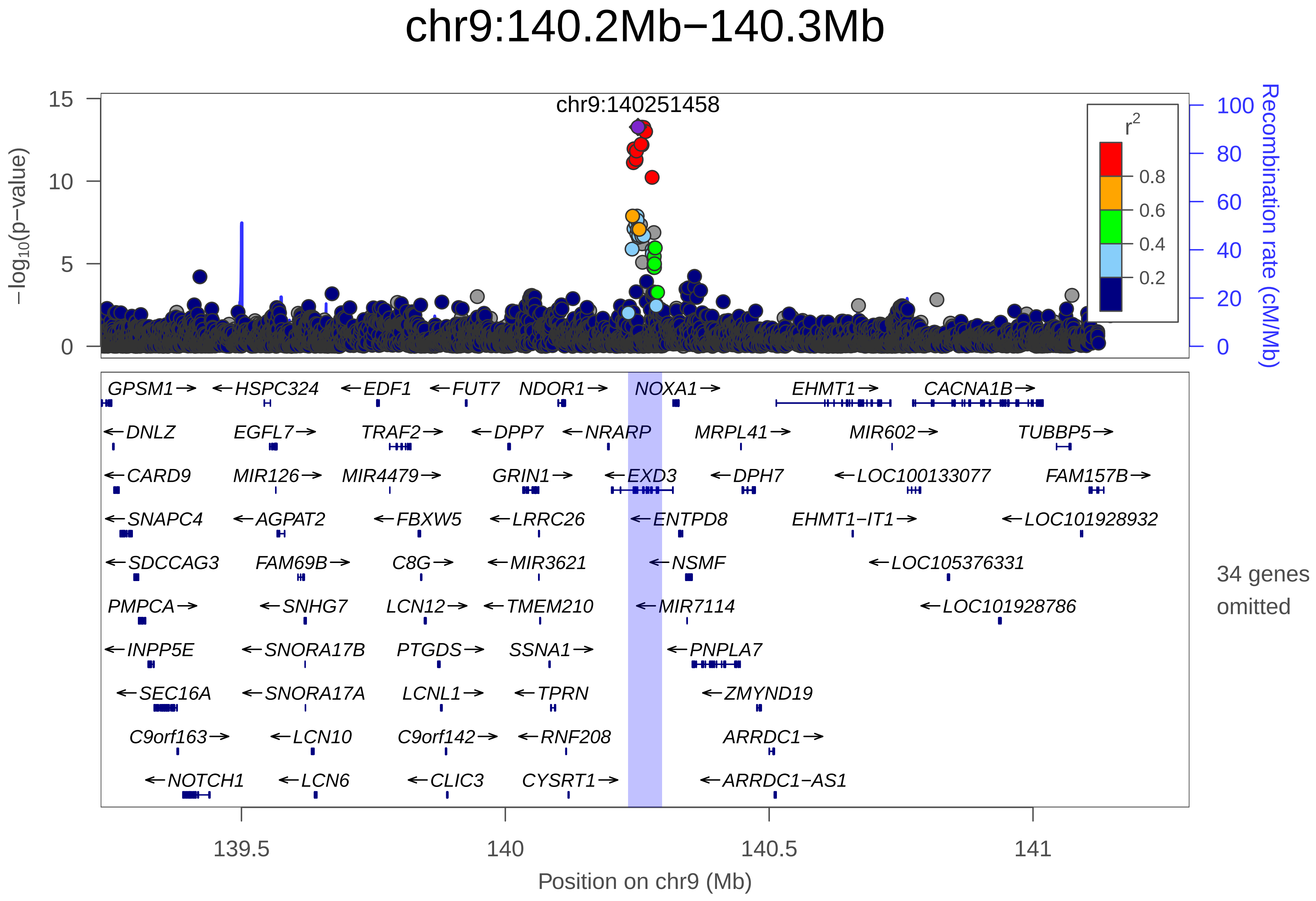

chr9:140.2Mb−140.3Mb
chr9:140251458
−log10(p−value)
Recombination rate (cM/Mb)
r2
0.8
0.6
0.4
0.2
GPSM1→
←HSPC324
←EDF1
←FUT7
NDOR1→
NOXA1→
EHMT1→
CACNA1B→
←DNLZ
EGFL7→
TRAF2→
←DPP7
←NRARP
MRPL41→
MIR602→
TUBBP5→
←CARD9
MIR126→
MIR4479→
GRIN1→
←EXD3
←DPH7
←LOC100133077
FAM157B→
←SNAPC4
←AGPAT2
←FBXW5
←LRRC26
←ENTPD8
EHMT1−IT1→
←LOC101928932
←SDCCAG3
FAM69B→
C8G→
←MIR3621
←NSMF
←LOC105376331
PMPCA→
←SNHG7
LCN12→
←TMEM210
←MIR7114
←LOC101928786
←INPP5E
←SNORA17B
PTGDS→
SSNA1→
←PNPLA7
←SEC16A
←SNORA17A
LCNL1→
←TPRN
←ZMYND19
C9orf163→
←LCN10
C9orf142→
←RNF208
ARRDC1→
←NOTCH1
←LCN6
←CLIC3
CYSRT1→
←ARRDC1−AS1
34 genes omitted
139.5
140
140.5
141
Position on chr9 (Mb)

| | |
|---|---|
| date: | Mon Dec 10 07:02:41 2018 |
| build: | hg19 |
| display range: | chr9:139233166−141296505 [139233166−141296505] |
| hilite range: | 140.233Mb − 140.297Mb [ 140233000 − 140297000 ] |
| reference SNP: | chr9:140251458 |
| number of SNPs plotted: | 6416 |
| min P−value: | 5.3E−14 [chr9:140251458] |
| max P−value: | 1E0 [chr9:139281570] |
| omitted Genes: | MIR4673, MIR4674, NALT1 |
| omitted Genes: | LINC01451, LOC100128593, MIR6722 |
| omitted Genes: | LCN8, LCN15, TMEM141 |
| omitted Genes: | CCDC183, CCDC183−AS1, RABL6 |
| omitted Genes: | MIR4292, C9orf172, PHPT1 |
| omitted Genes: | MAMDC4, ABCA2, C9orf139 |
| omitted Genes: | NPDC1, ENTPD2, SAPCD2 |
| omitted Genes: | UAP1L1, MAN1B1−AS1, MAN1B1 |
| omitted Genes: | ANAPC2, TMEM203, RNF224 |
| omitted Genes: | SLC34A3, TUBB4B, FAM166A |
| omitted Genes: | STPG3−AS1, STPG3, NELFB |
| omitted Genes: | TOR4A |

Make more plots at http://csg.sph.umich.edu/locuszoom/
