## Supplementary Figure 34 for "Genome-wide Association Study of Multisite Chronic Pain in UK Biobank"

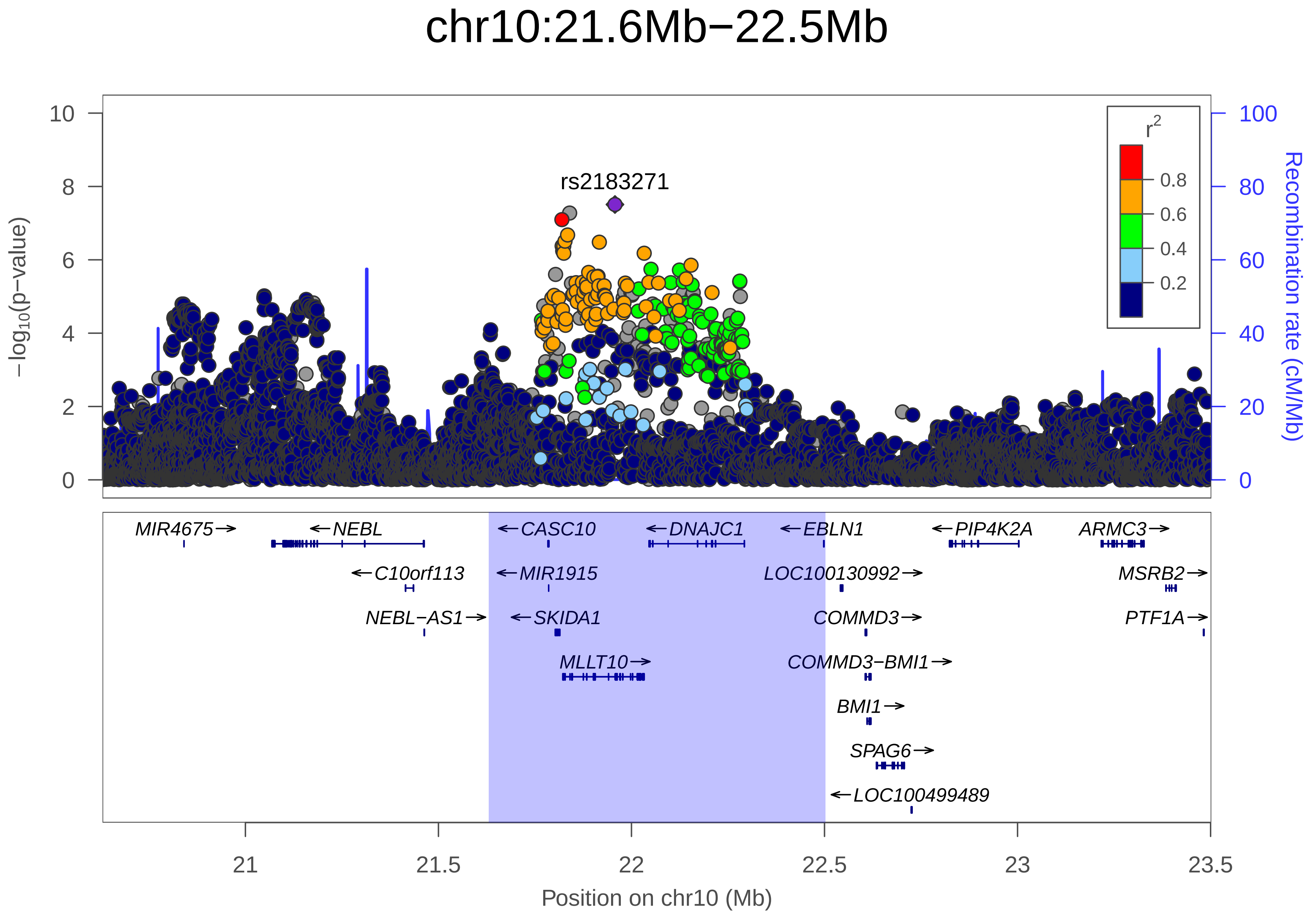

chr10:21.6Mb−22.5Mb
rs2183271
r²
0.8
0.6
0.4
0.2
−log10(p−value)
Recombination rate (cM/Mb)
10
8
6
4
2
0
100
80
60
40
20
0
MIR4675→
←NEBL
←CASC10
←DNAJC1
←EBLN1
←PIP4K2A
ARMC3→
←C10orf113
←MIR1915
LOC100130992→
MSRB2→
NEBL−AS1→
←SKIDA1
COMMD3→
PTF1A→
MLLT10→
COMMD3−BMI1→
BMI1→
SPAG6→
←LOC100499489
21
21.5
22
22.5
23
23.5
Position on chr10 (Mb)

| | |
|---|---|
| date: | Mon Dec 10 07:16:54 2018 |
| build: | hg19 |
| display range: | chr10:20629890−23501895 [20629890−23501895] |
| hilite range: | 21.63Mb − 22.502Mb [ 21630000 − 22502000 ] |
| reference SNP: | chr10:21957229 |
| number of SNPs plotted: | 8667 |
| min P−value: | 3.1E−8 [chr10:21957229] |
| max P−value: | 1E0 [chr10:20777029] |

Make more plots at http://csg.sph.umich.edu/locuszoom/
