## Supplementary Figure 35 for "Genome-wide Association Study of Multisite Chronic Pain in UK Biobank"

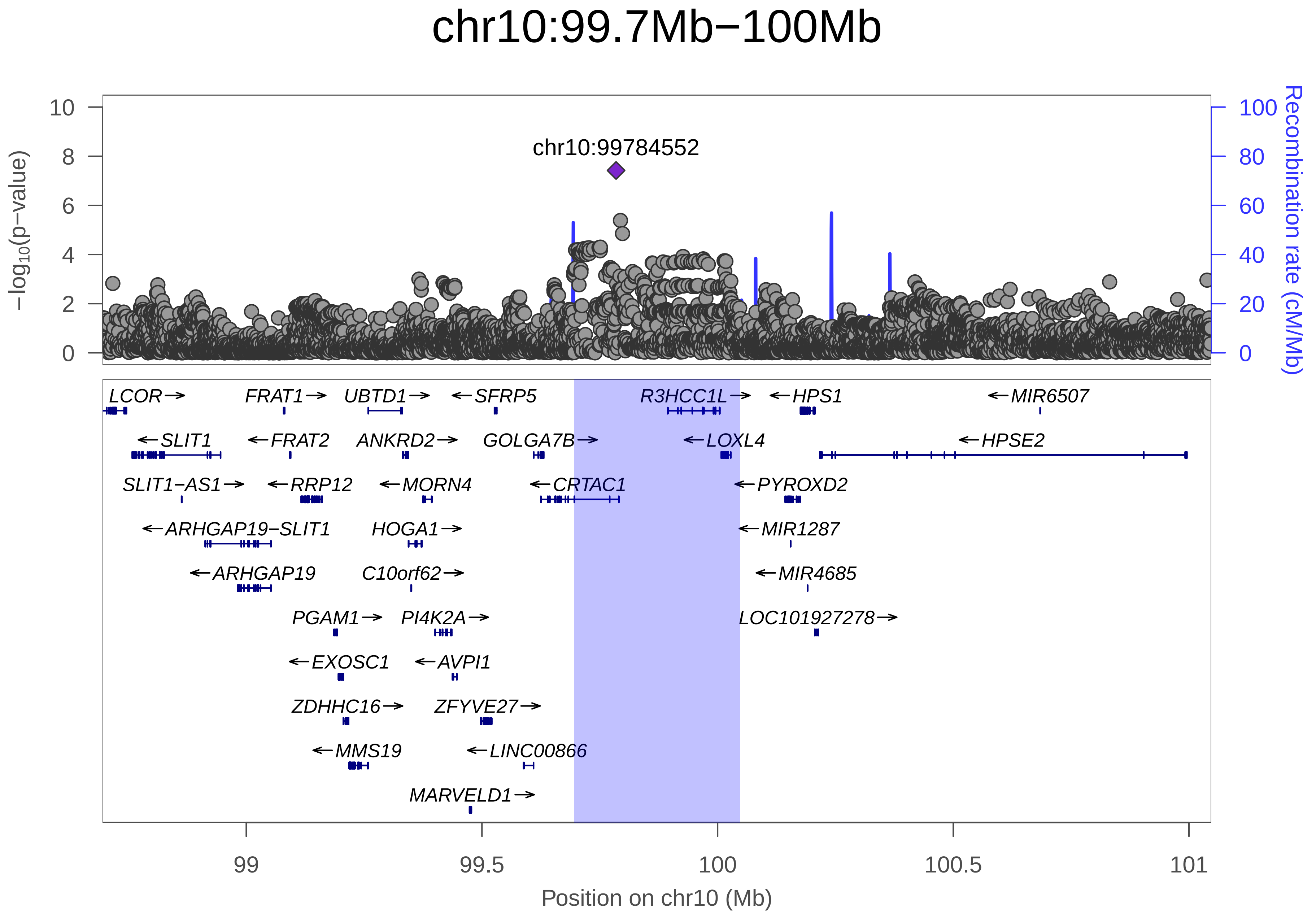
chr10:99.7Mb−100Mb
chr10:99784552
−log10(p−value)
Recombination rate (cM/Mb)
0
2
4
6
8
10
0
20
40
60
80
100
LCOR
FRAT1
UBTD1
SFRP5
R3HCC1L
HPS1
MIR6507
SLIT1
FRAT2
ANKRD2
GOLGA7B
LOXL4
HPSE2
SLIT1−AS1
RRP12
MORN4
CRTAC1
PYROXD2
ARHGAP19−SLIT1
HOGA1
MIR1287
ARHGAP19
C10orf62
MIR4685
PGAM1
PI4K2A
LOC101927278
EXOSC1
AVPI1
ZDHHC16
ZFYVE27
MMS19
LINC00866
MARVELD1
99
99.5
100
100.5
101
Position on chr10 (Mb)

| | |
|---|---|
| date: | Mon Dec 10 07:34:30 2018 |
| build: | hg19 |
| display range: | chr10:98694970−101047501 [98694970−101047501] |
| hilite range: | 99.695Mb − 100.048Mb [ 99695000 − 100048000 ] |
| reference SNP: | chr10:99784552 |
| number of SNPs plotted: | 7405 |
| min P−value: | 3.8E−8 [chr10:99784552] |
| max P−value: | 1E0 [chr10:98791578] |
| Warning: | No usable LD information for reference SNP. |

Make more plots at http://csg.sph.umich.edu/locuszoom/
