## Supplementary Figure 36 for "Genome-wide Association Study of Multisite Chronic Pain in UK Biobank"

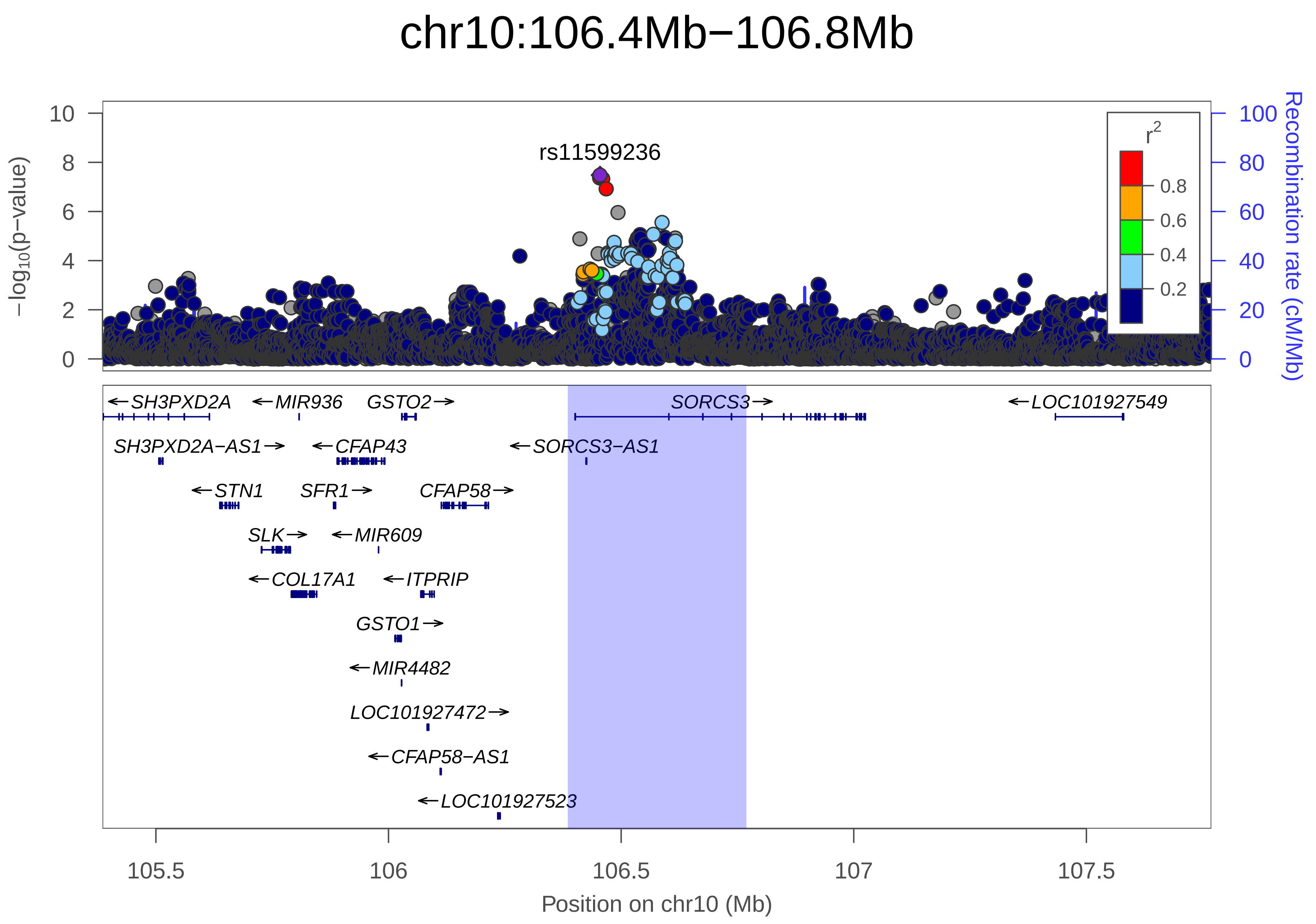
chr10:106.4Mb−106.8Mb
rs11599236
r²
0.8
0.6
0.4
0.2
−log10(p−value)
Recombination rate (cM/Mb)
SH3PXD2A
MIR936
GSTO2
SORCS3
LOC101927549
SH3PXD2A−AS1
CFAP43
SORCS3−AS1
STN1
SFR1
CFAP58
SLK
MIR609
COL17A1
ITPRIP
GSTO1
MIR4482
LOC101927472
CFAP58−AS1
LOC101927523
105.5
106
106.5
107
107.5
Position on chr10 (Mb)

| | |
|---|---|
| date: | Mon Dec 10 08:03:19 2018 |
| build: | hg19 |
| display range: | chr10:105385271−107768514 [105385271−107768514] |
| hilite range: | 106.385Mb − 106.769Mb [ 106385000 − 106769000 ] |
| reference SNP: | chr10:106454672 |
| number of SNPs plotted: | 7417 |
| min P−value: | 3.3E−8 [chr10:106454672] |
| max P−value: | 1E0 [chr10:105390122] |

Make more plots at http://csg.sph.umich.edu/locuszoom/
