## Supplementary Figure 37 for "Genome-wide Association Study of Multisite Chronic Pain in UK Biobank"

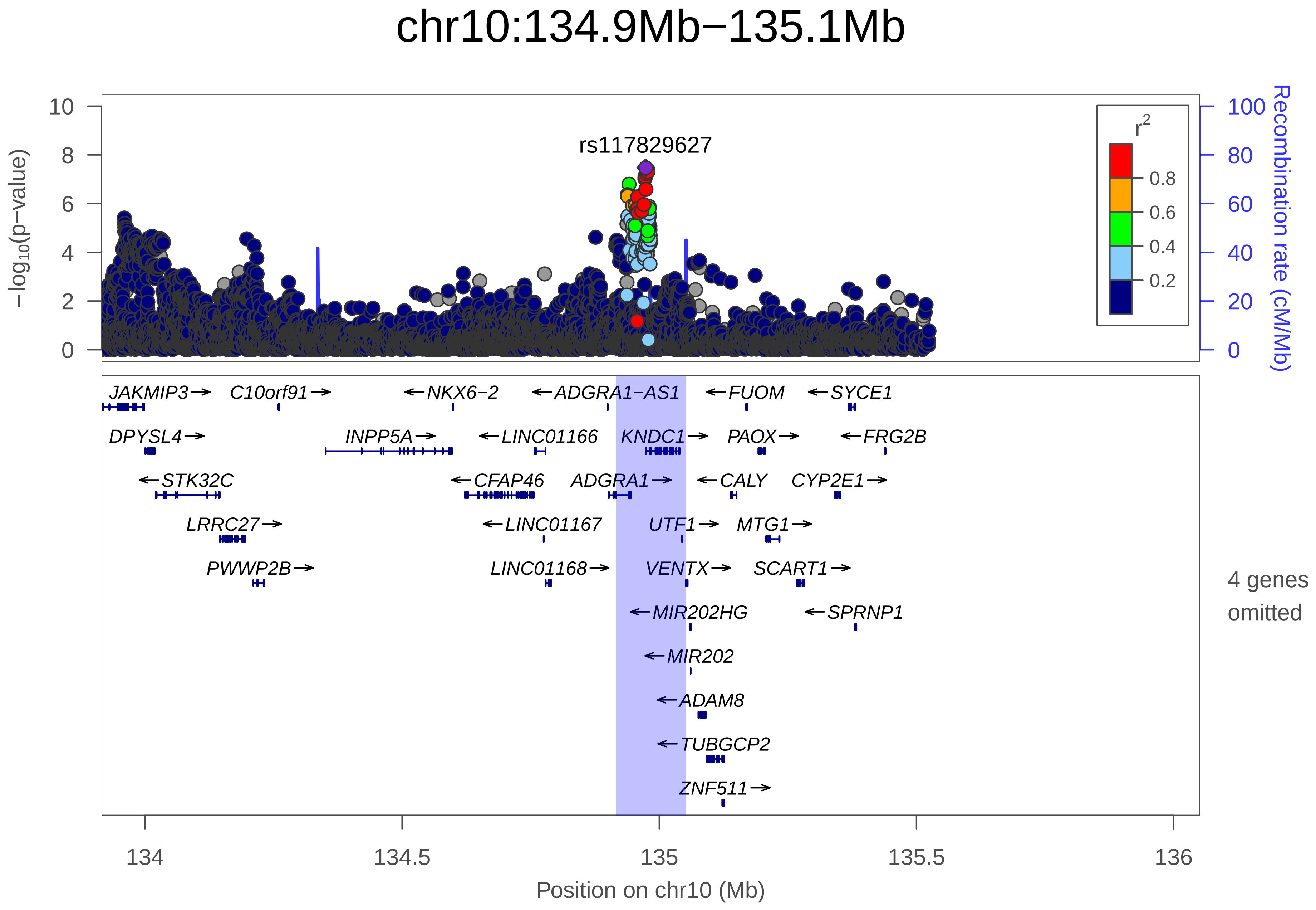
chr10:134.9Mb−135.1Mb
rs117829627
−log10(p−value)
Recombination rate (cM/Mb)
r2
0.8
0.6
0.4
0.2
10
8
6
4
2
0
100
80
60
40
20
0
JAKMIP3
C10orf91
NKX6−2
ADGRA1−AS1
FUOM
SYCE1
DPYSL4
INPP5A
LINC01166
KNDC1
PAOX
FRG2B
STK32C
CFAP46
ADGRA1
CALY
CYP2E1
LRRC27
LINC01167
UTF1
MTG1
PWWP2B
LINC01168
VENTX
SCART1
MIR202HG
SPRNP1
MIR202
ADAM8
TUBGCP2
ZNF511
4 genes omitted
134
134.5
135
135.5
136
Position on chr10 (Mb)

| | |
|---|---|
| date: | Mon Dec 10 07:02:40 2018 |
| build: | hg19 |
| display range: | chr10:133915611−136051562 [133915611−136051562] |
| hilite range: | 134.916Mb − 135.052Mb [ 134916000 − 135052000 ] |
| reference SNP: | chr10:134973637 |
| number of SNPs plotted: | 8012 |
| min P−value: | 3.4E−8 [chr10:134973637] |
| max P−value: | 1E0 [chr10:133928336] |
| omitted Genes: | PRAP1, ECHS1, MIR3944 |
| omitted Genes: | SPRN |

Make more plots at http://csg.sph.umich.edu/locuszoom/
