## Supplementary Figure 38 for "Genome-wide Association Study of Multisite Chronic Pain in UK Biobank"

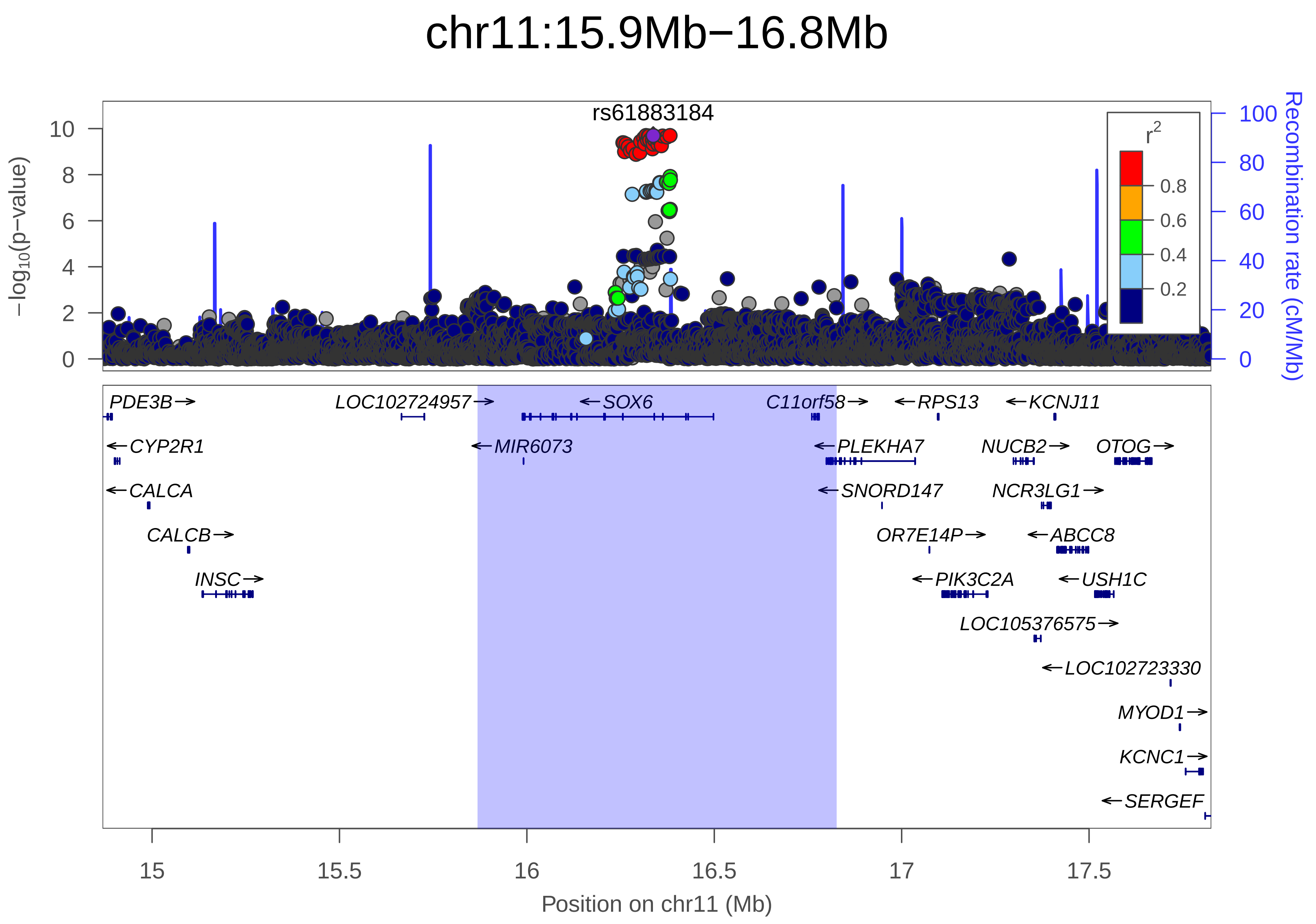

chr11:15.9Mb−16.8Mb
rs61883184
−log10(p−value)
Recombination rate (cM/Mb)
r2
0.8
0.6
0.4
0.2
PDE3B
CYP2R1
CALCA
CALCB
INSC
LOC102724957
MIR6073
SOX6
C11orf58
PLEKHA7
SNORD147
RPS13
OR7E14P
PIK3C2A
KCNJ11
NUCB2
NCR3LG1
ABCC8
USH1C
LOC105376575
LOC102723330
MYOD1
KCNC1
SERGEF
OTOG
15
15.5
16
16.5
17
17.5
Position on chr11 (Mb)

| | |
|---|---|
| date: | Mon Dec 10 07:20:26 2018 |
| build: | hg19 |
| display range: | chr11:14867726−17826201 [14867726−17826201] |
| hilite range: | 15.868Mb − 16.826Mb [ 15868000 − 16826000 ] |
| reference SNP: | chr11:16337092 |
| number of SNPs plotted: | 8855 |
| min P−value: | 2E−10 [chr11:16317779] |
| max P−value: | 1E0 [chr11:15175746] |

Make more plots at http://csg.sph.umich.edu/locuszoom/
