## Supplementary Figure 39 for "Genome-wide Association Study of Multisite Chronic Pain in UK Biobank"

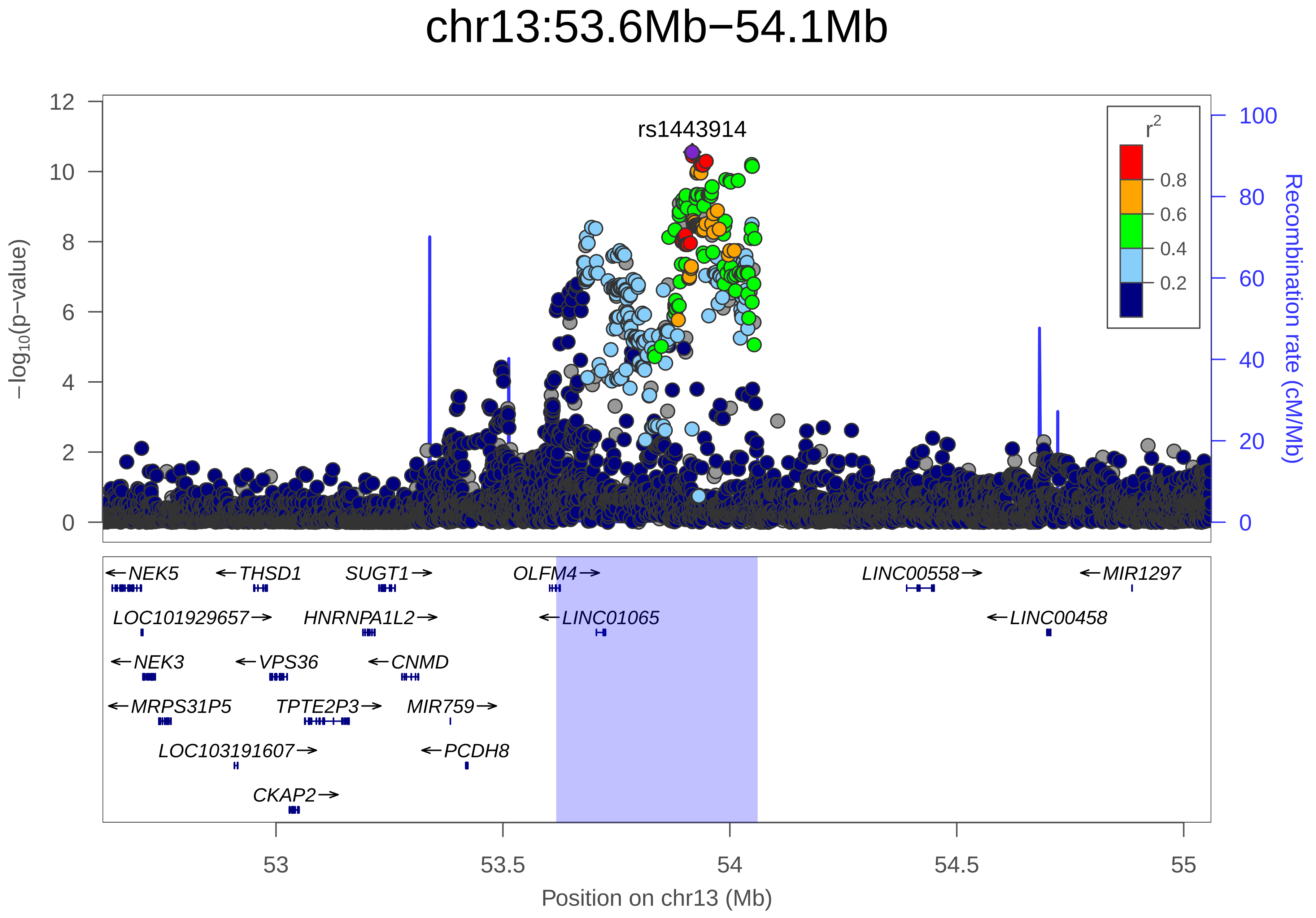

chr13:53.6Mb−54.1Mb
rs1443914
−log10(p−value)
12
10
8
6
4
2
0
r2
0.8
0.6
0.4
0.2
100
80
60
40
20
0
Recombination rate (cM/Mb)
←NEK5
←THSD1
SUGT1→
OLFM4→
LINC00558→
←MIR1297
LOC101929657→
HNRNPA1L2→
←LINC01065
←LINC00458
←NEK3
←VPS36
←CNMD
←MRPS31P5
TPTE2P3→
MIR759→
LOC103191607→
←PCDH8
CKAP2→
53
53.5
54
54.5
55
Position on chr13 (Mb)

| | |
|---|---|
| date: | Mon Dec 10 07:34:29 2018 |
| build: | hg19 |
| display range: | chr13:52617781−55060827 [52617781−55060827] |
| hilite range: | 53.618Mb − 54.061Mb [ 53618000 − 54061000 ] |
| reference SNP: | chr13:53917230 |
| number of SNPs plotted: | 7769 |
| min P−value: | 2.8E−11 [chr13:53917230] |
| max P−value: | 1E0 [chr13:52714964] |

Make more plots at http://csg.sph.umich.edu/locuszoom/
