## Supplementary Figure 40 for "Genome-wide Association Study of Multisite Chronic Pain in UK Biobank"

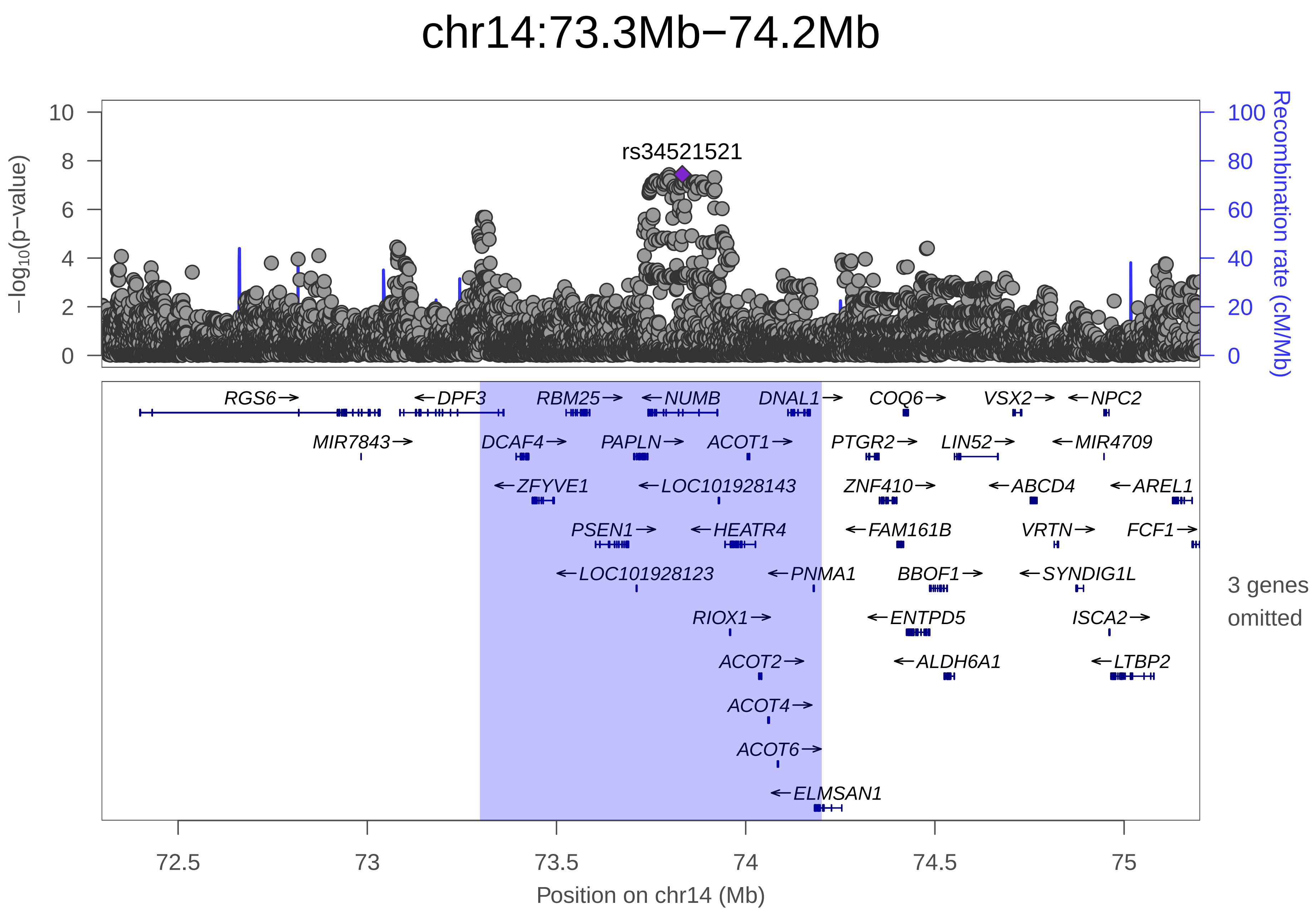

chr14:73.3Mb−74.2Mb
rs34521521
−log10(p−value)
Recombination rate (cM/Mb)
RGS6 DPF3 RBM25 NUMB DNAL1 COQ6 VSX2 NPC2
MIR7843 DCAF4 PAPLN ACOT1 PTGR2 LIN52 MIR4709
ZFYVE1 LOC101928143 ZNF410 ABCD4 AREL1
PSEN1 HEATR4 FAM161B VRTN FCF1
LOC101928123 PNMA1 BBOF1 SYNDIG1L
RIOX1 ENTPD5 ISCA2
ACOT2 ALDH6A1 LTBP2
ACOT4
ACOT6
ELMSAN1
3 genes omitted
Position on chr14 (Mb)

| | |
|---|---|
| date: | Mon Dec 10 08:03:18 2018 |
| build: | hg19 |
| display range: | chr14:72297741−75200876 [72297741−75200876] |
| hilite range: | 73.298Mb − 74.201Mb [ 73298000 − 74201000 ] |
| reference SNP: | chr14:73832318 |
| number of SNPs plotted: | 11036 |
| min P−value: | 3.6E−8 [chr14:73832318] |
| max P−value: | 1E0 [chr14:72375036] |
| omitted Genes: | MIR4505, LOC100506476, LOC100506498 |
| Warning: | No usable LD information for reference SNP. |

Make more plots at http://csg.sph.umich.edu/locuszoom/
