## Supplementary Figure 41 for "Genome-wide Association Study of Multisite Chronic Pain in UK Biobank"

### chr14:103.8Mb−104.5Mb

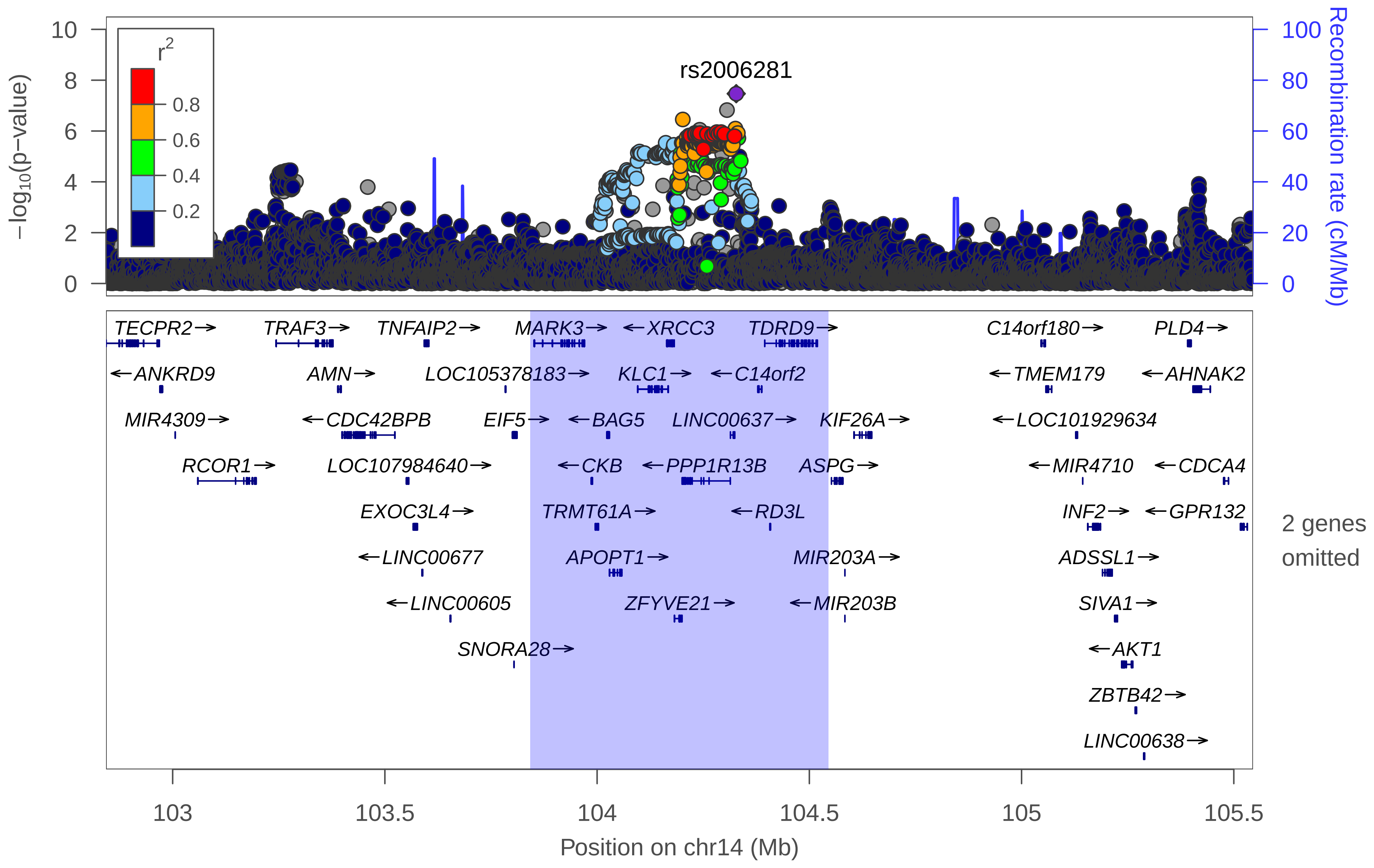

| | |
|---|---|
| date: | Mon Dec 10 07:08:40 2018 |
| build: | hg19 |
| display range: | chr14:102843045−105545285 [102843045−105545285] |
| hilite range: | 103.843Mb − 104.545Mb [ 103843000 − 104545000 ] |
| reference SNP: | chr14:104327732 |
| number of SNPs plotted: | 10657 |
| min P−value: | 3.4E−8 [chr14:104327732] |
| max P−value: | 1E0 [chr14:102904986] |
| omitted Genes: | CEP170B, C14orf79 |

Make more plots at http://csg.sph.umich.edu/locuszoom/
