## Supplementary Figure 42 for "Genome-wide Association Study of Multisite Chronic Pain in UK Biobank"

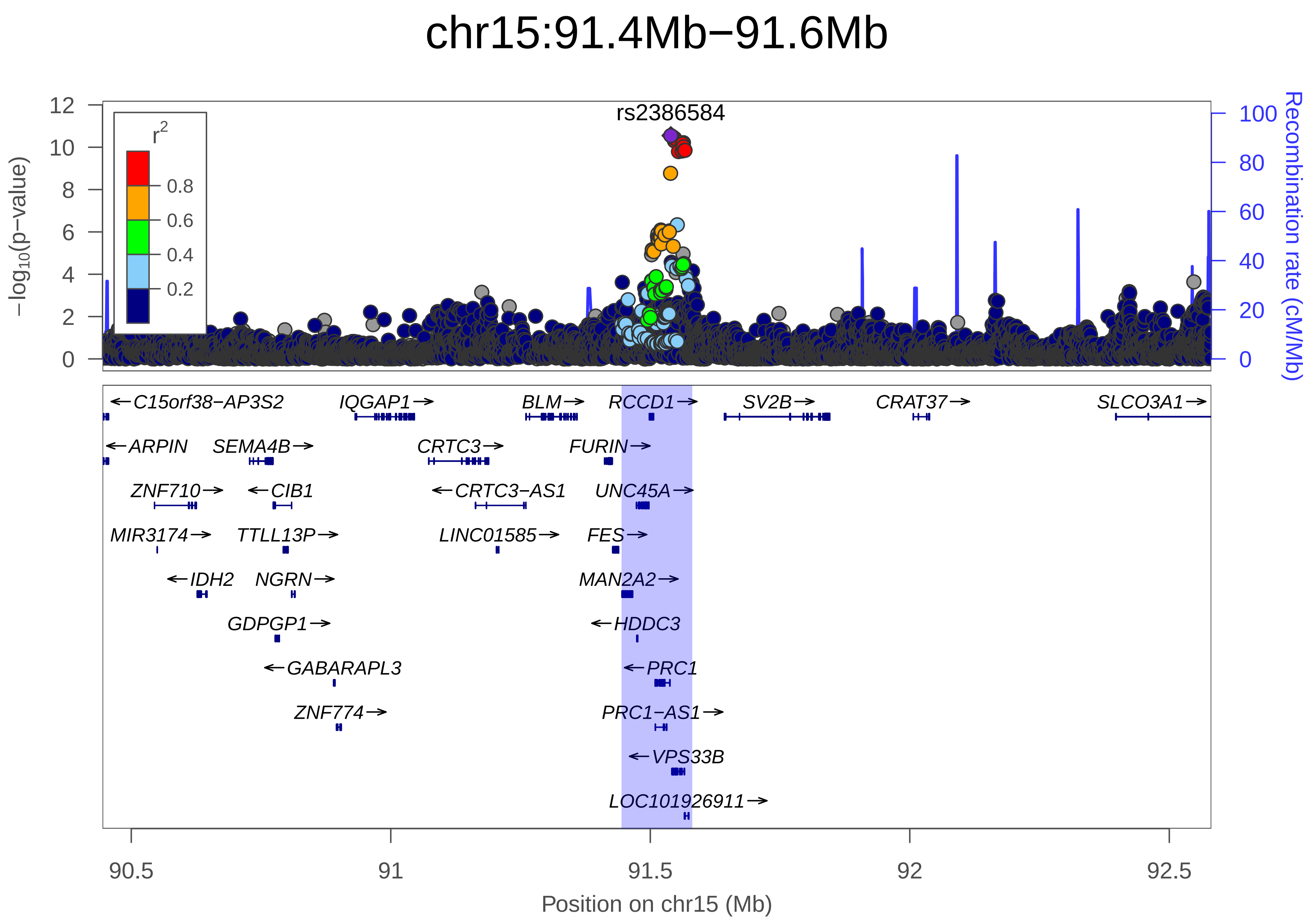
chr15:91.4Mb−91.6Mb
rs2386584
r2
0.8
0.6
0.4
0.2
−log10(p−value)
Recombination rate (cM/Mb)
0
2
4
6
8
10
12
0
20
40
60
80
100
←C15orf38−AP3S2
IQGAP1→
BLM→
RCCD1→
SV2B→
CRAT37→
SLCO3A1→
←ARPIN
SEMA4B→
CRTC3→
FURIN→
ZNF710→
←CIB1
←CRTC3−AS1
UNC45A→
MIR3174→
TTLL13P→
LINC01585→
FES→
←IDH2
NGRN→
MAN2A2→
GDPGP1→
←HDDC3
←GABARAPL3
←PRC1
ZNF774→
PRC1−AS1→
←VPS33B
LOC101926911→
90.5
91
91.5
92
92.5
Position on chr15 (Mb)

| | |
|---|---|
| date: | Mon Dec 10 07:16:54 2018 |
| build: | hg19 |
| display range: | chr15:90444573−92580961 [90444573−92580961] |
| hilite range: | 91.445Mb − 91.581Mb [ 91445000 − 91581000 ] |
| reference SNP: | chr15:91539572 |
| number of SNPs plotted: | 8241 |
| min P−value: | 2.8E−11 [chr15:91539572] |
| max P−value: | 1E0 [chr15:90488772] |

Make more plots at http://csg.sph.umich.edu/locuszoom/
