## Supplementary Figure 43 for "Genome-wide Association Study of Multisite Chronic Pain in UK Biobank"

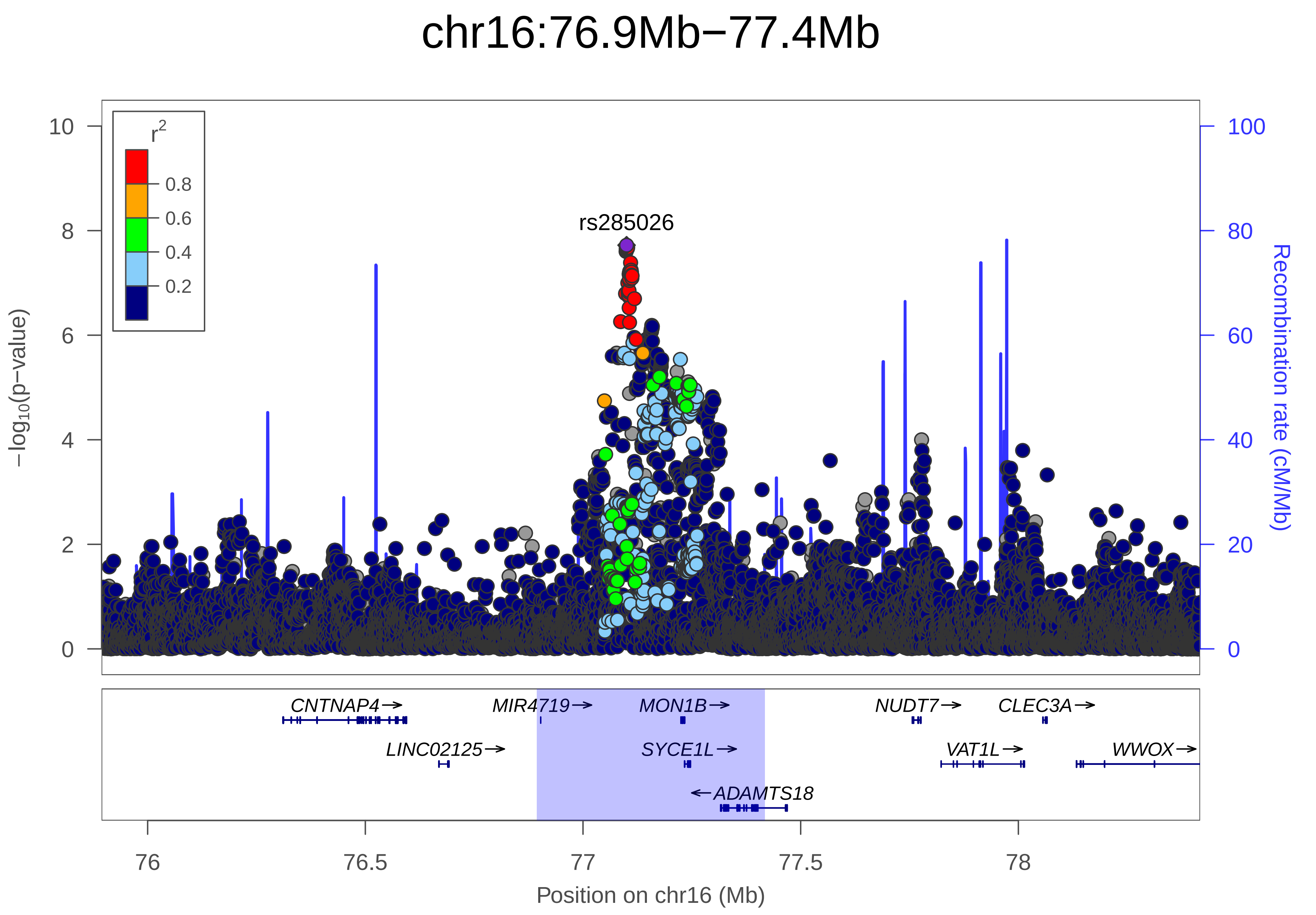
chr16:76.9Mb−77.4Mb
r2
0.8
0.6
0.4
0.2
rs285026
10
8
6
4
2
0
−log10(p−value)
100
80
60
40
20
0
Recombination rate (cM/Mb)
CNTNAP4→
MIR4719→
MON1B→
NUDT7→
CLEC3A→
LINC02125→
SYCE1L→
VAT1L→
WWOX→
←ADAMTS18
76
76.5
77
77.5
78
Position on chr16 (Mb)

| | |
|---|---|
| date: | Mon Dec 10 07:48:16 2018 |
| build: | hg19 |
| display range: | chr16:75894153−78417509 [75894153−78417509] |
| hilite range: | 76.894Mb − 77.418Mb [ 76894000 − 77418000 ] |
| reference SNP: | chr16:77100089 |
| number of SNPs plotted: | 13912 |
| min P−value: | 1.9E−8 [chr16:77100089] |
| max P−value: | 1E0 [chr16:75915873] |

Make more plots at http://csg.sph.umich.edu/locuszoom/
