## Supplementary Figure 44 for "Genome-wide Association Study of Multisite Chronic Pain in UK Biobank"

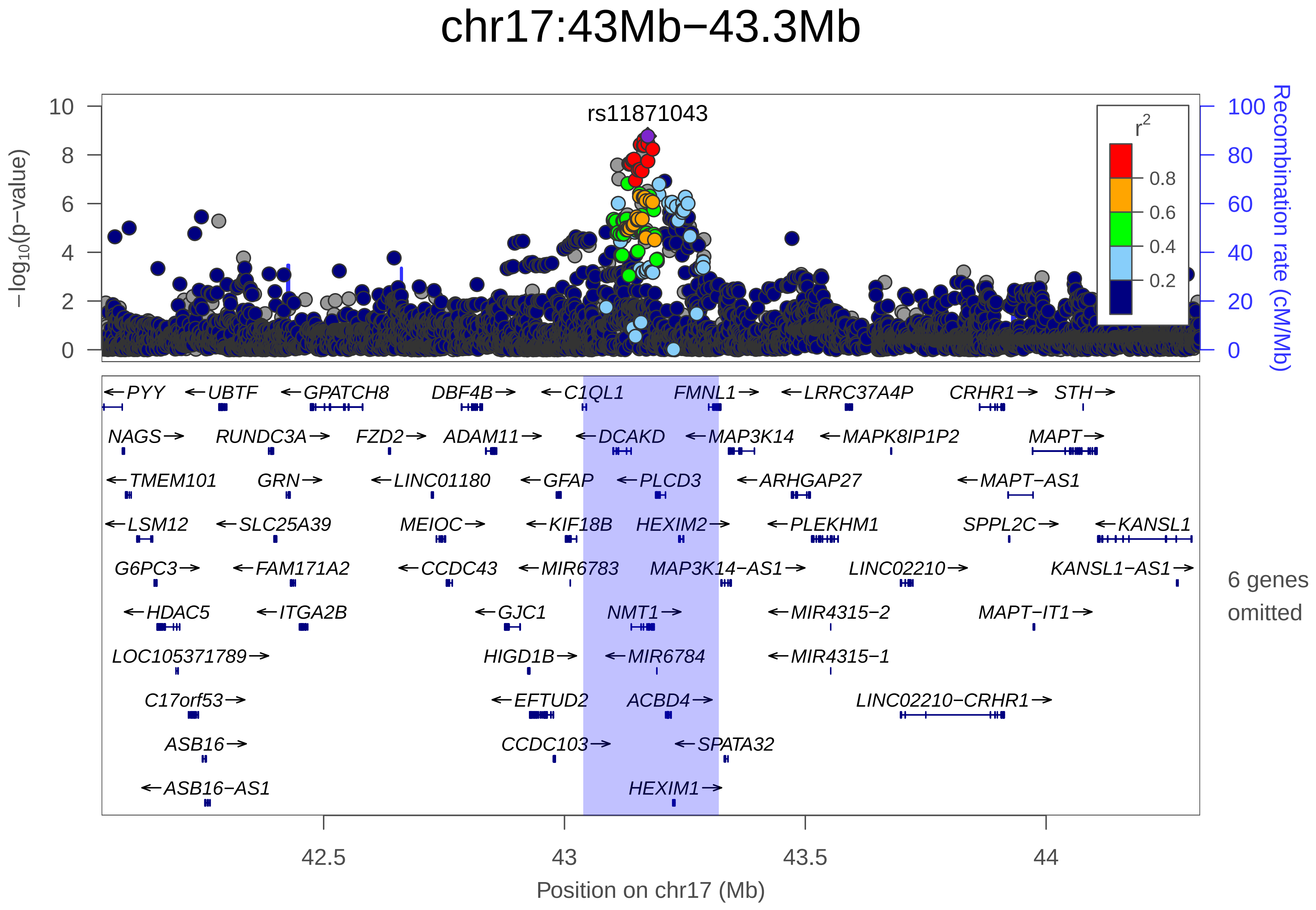

chr17:43Mb−43.3Mb
rs11871043
−log10(p−value)
Recombination rate (cM/Mb)
r2
0.8
0.6
0.4
0.2
PYY
UBTF
GPATCH8
DBF4B
C1QL1
FMNL1
LRRC37A4P
CRHR1
STH
NAGS
RUNDC3A
FZD2
ADAM11
DCAKD
MAP3K14
MAPK8IP1P2
MAPT
TMEM101
GRN
LINC01180
GFAP
PLCD3
ARHGAP27
MAPT−AS1
LSM12
SLC25A39
MEIOC
KIF18B
HEXIM2
PLEKHM1
SPPL2C
KANSL1
G6PC3
FAM171A2
CCDC43
MIR6783
MAP3K14−AS1
LINC02210
KANSL1−AS1
HDAC5
ITGA2B
GJC1
NMT1
MIR4315−2
MAPT−IT1
LOC105371789
HIGD1B
MIR6784
MIR4315−1
C17orf53
EFTUD2
ACBD4
LINC02210−CRHR1
ASB16
CCDC103
SPATA32
ASB16−AS1
HEXIM1
6 genes omitted
42.5
43
43.5
44
Position on chr17 (Mb)

| | |
|---|---|
| date: | Mon Dec 10 08:05:07 2018 |
| build: | hg19 |
| display range: | chr17:42038865−44319770 [42038865−44319770] |
| hilite range: | 43.039Mb − 43.32Mb [ 43039000 − 43320000 ] |
| reference SNP: | chr17:43172849 |
| number of SNPs plotted: | 8619 |
| min P−value: | 1.7E−9 [chr17:43172849] |
| max P−value: | 1E0 [chr17:42303305] |
| omitted Genes: | TMUB2, ATXN7L3, MIR6782 |
| omitted Genes: | SLC4A1, RUNDC3A−AS1, LOC105371795 |

Make more plots at http://csg.sph.umich.edu/locuszoom/
