## Supplementary Figure 45 for "Genome-wide Association Study of Multisite Chronic Pain in UK Biobank"

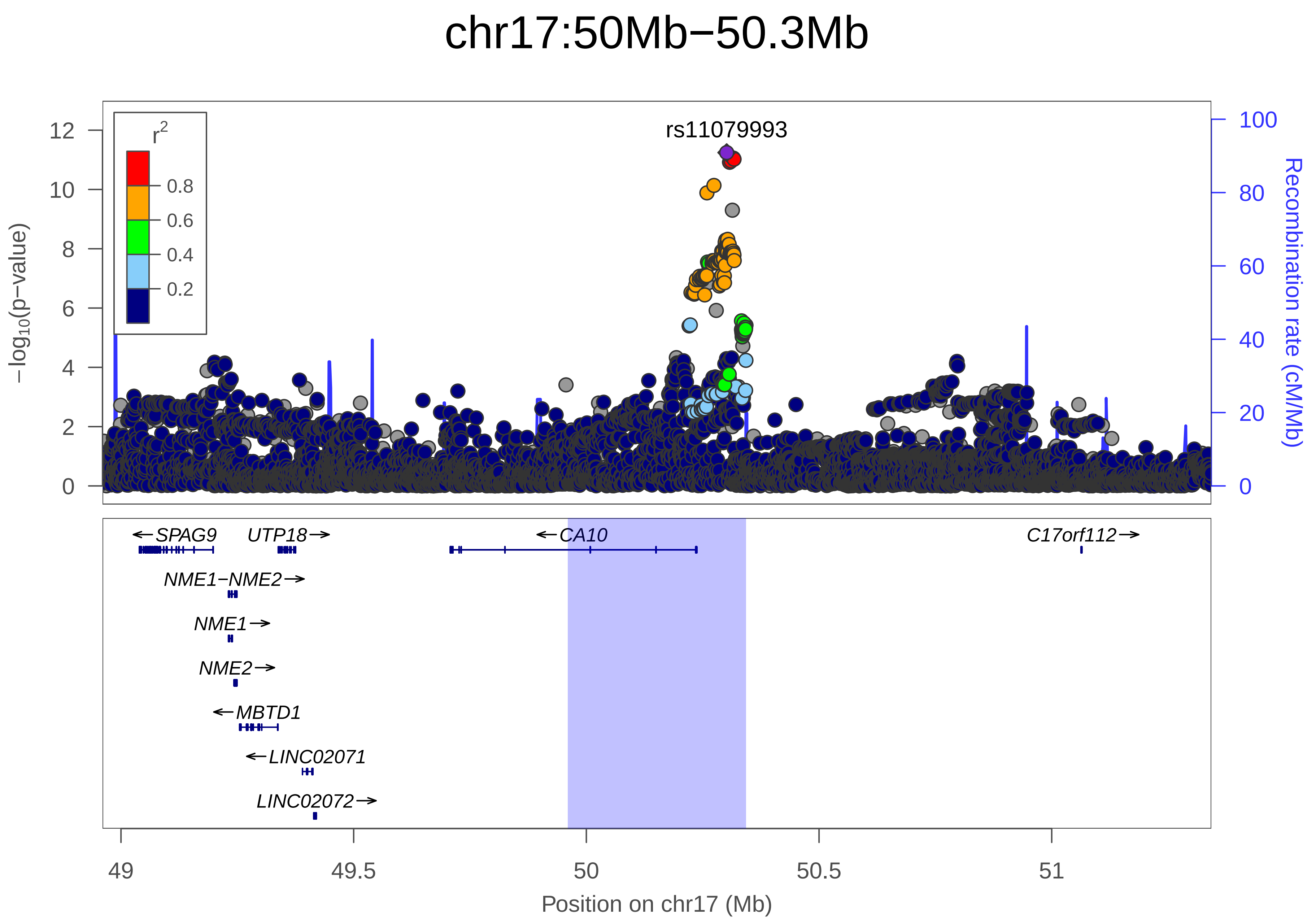
chr17:50Mb−50.3Mb
rs11079993
r²
0.8
0.6
0.4
0.2
−log10(p−value)
Recombination rate (cM/Mb)
SPAG9
UTP18
CA10
C17orf112
NME1−NME2
NME1
NME2
MBTD1
LINC02071
LINC02072
49
49.5
50
50.5
51
Position on chr17 (Mb)

| | |
|---|---|
| date: | Mon Dec 10 07:02:39 2018 |
| build: | hg19 |
| display range: | chr17:48960309−51342947 [48960309−51342947] |
| hilite range: | 49.96Mb − 50.343Mb [ 49960000 − 50343000 ] |
| reference SNP: | chr17:50301552 |
| number of SNPs plotted: | 8116 |
| min P−value: | 5.7E−12 [chr17:50301552] |
| max P−value: | 1E0 [chr17:49419123] |

Make more plots at http://csg.sph.umich.edu/locuszoom/
