## Supplementary Figure 46 for "Genome-wide Association Study of Multisite Chronic Pain in UK Biobank"

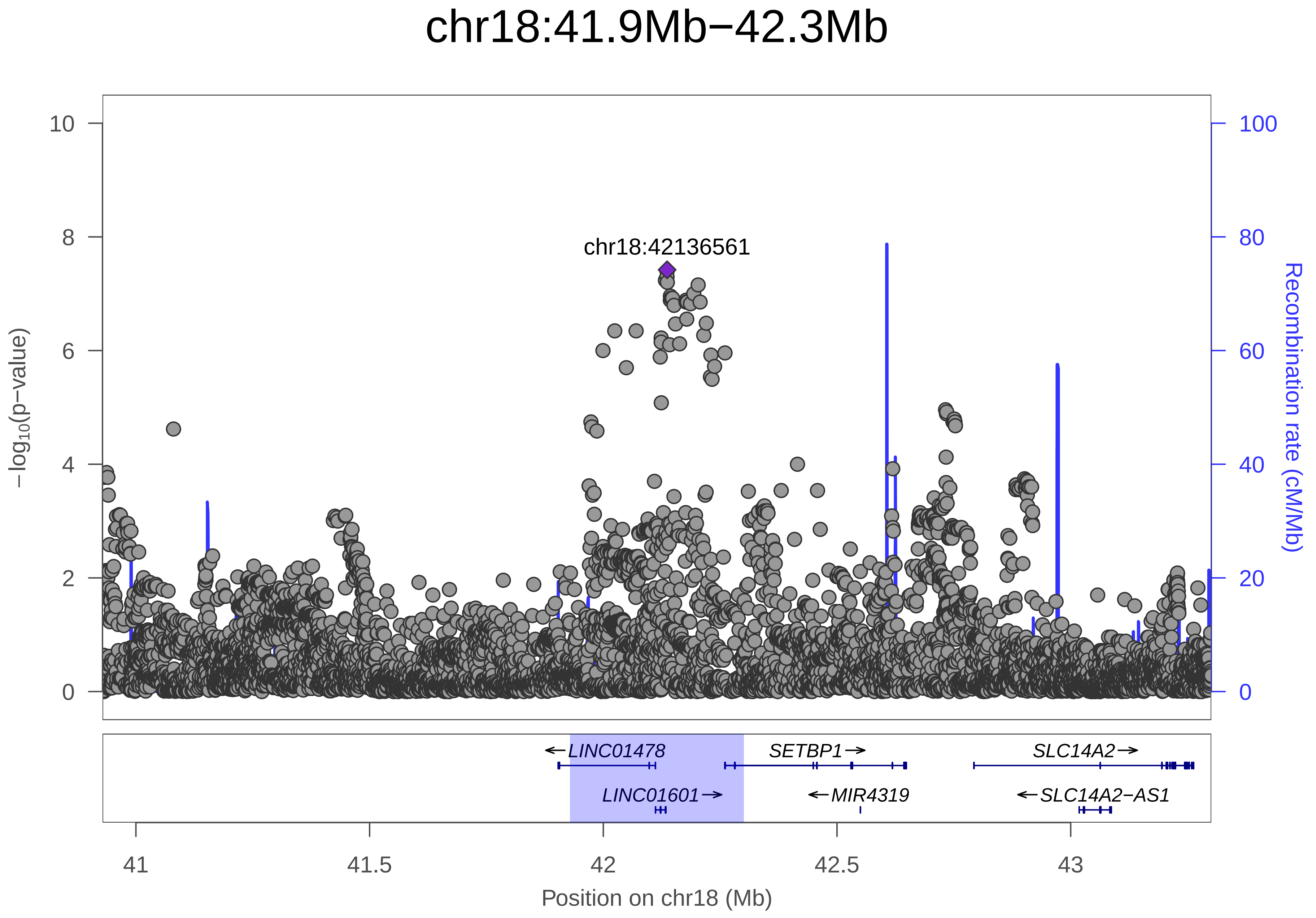

chr18:41.9Mb−42.3Mb
chr18:42136561
−log10(p−value)
Recombination rate (cM/Mb)
0
2
4
6
8
10
0
20
40
60
80
100
LINC01478
SETBP1
SLC14A2
LINC01601
MIR4319
SLC14A2−AS1
41
41.5
42
42.5
43
Position on chr18 (Mb)

| | |
|---|---|
| date: | Mon Dec 10 07:16:53 2018 |
| build: | hg19 |
| display range: | chr18:40928562−43300875 [40928562−43300875] |
| hilite range: | 41.929Mb − 42.301Mb [ 41929000 − 42301000 ] |
| reference SNP: | chr18:42136561 |
| number of SNPs plotted: | 7748 |
| min P−value: | 3.8E−8 [chr18:42136561] |
| max P−value: | 1E0 [chr18:40931364] |
| Warning: | No usable LD information for reference SNP. |

Make more plots at http://csg.sph.umich.edu/locuszoom/
