## Supplementary Figure 47 for "Genome-wide Association Study of Multisite Chronic Pain in UK Biobank"

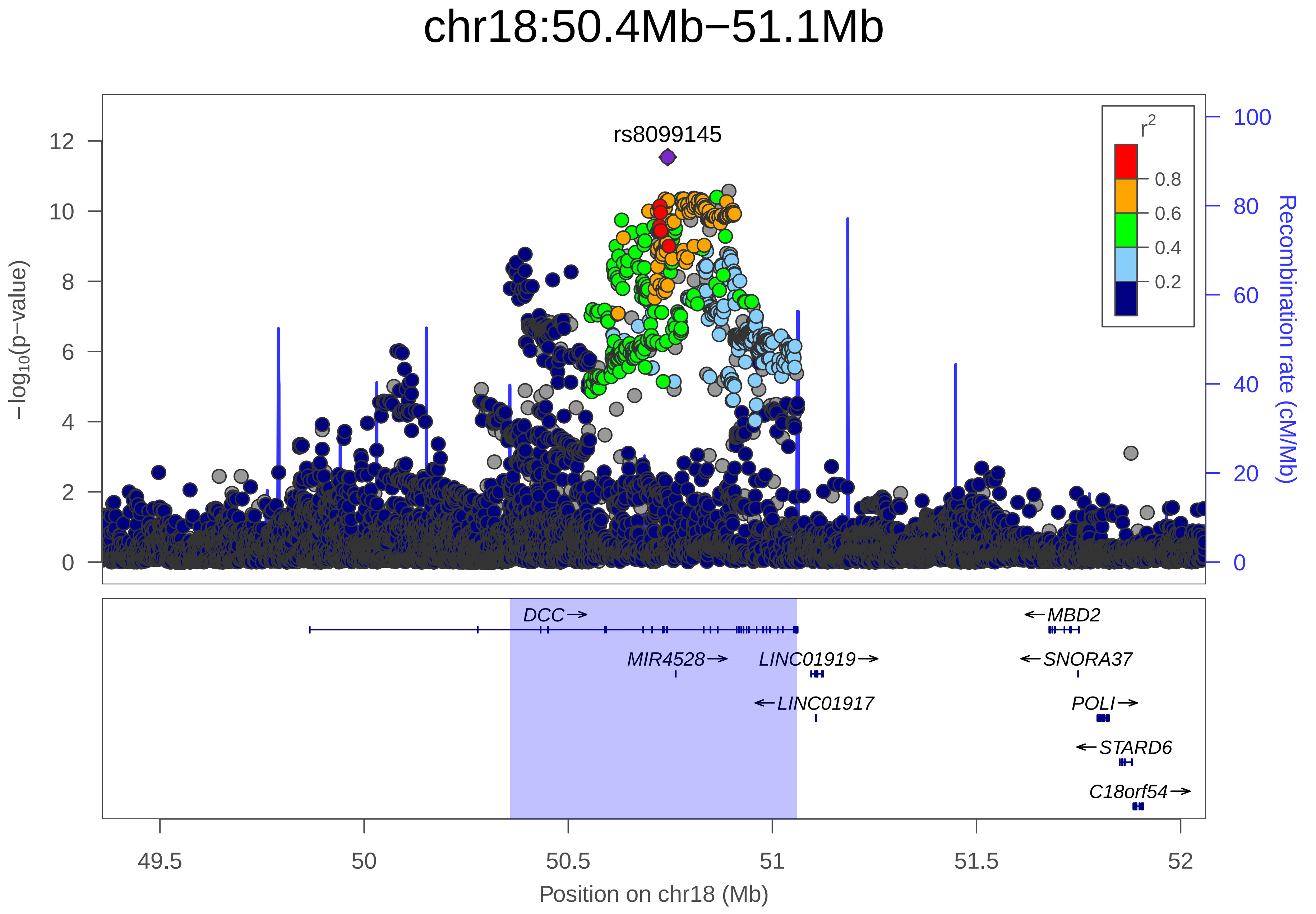

chr18:50.4Mb−51.1Mb
rs8099145
−log10(p−value)
Recombination rate (cM/Mb)
r2
0.8
0.6
0.4
0.2
12
10
8
6
4
2
0
100
80
60
40
20
0
DCC→
MIR4528→
LINC01919→
←LINC01917
←MBD2
←SNORA37
POLI→
←STARD6
C18orf54→
49.5
50
50.5
51
51.5
52
Position on chr18 (Mb)

| | |
|---|---|
| date: | Mon Dec 10 07:40:40 2018 |
| build: | hg19 |
| display range: | chr18:49358109−52061399 [49358109−52061399] |
| hilite range: | 50.358Mb − 51.061Mb [ 50358000 − 51061000 ] |
| reference SNP: | chr18:50743672 |
| number of SNPs plotted: | 11363 |
| min P−value: | 2.9E−12 [chr18:50743672] |
| max P−value: | 1E0 [chr18:49525912] |

Make more plots at http://csg.sph.umich.edu/locuszoom/
