## Supplementary Figure 48 for "Genome-wide Association Study of Multisite Chronic Pain in UK Biobank"

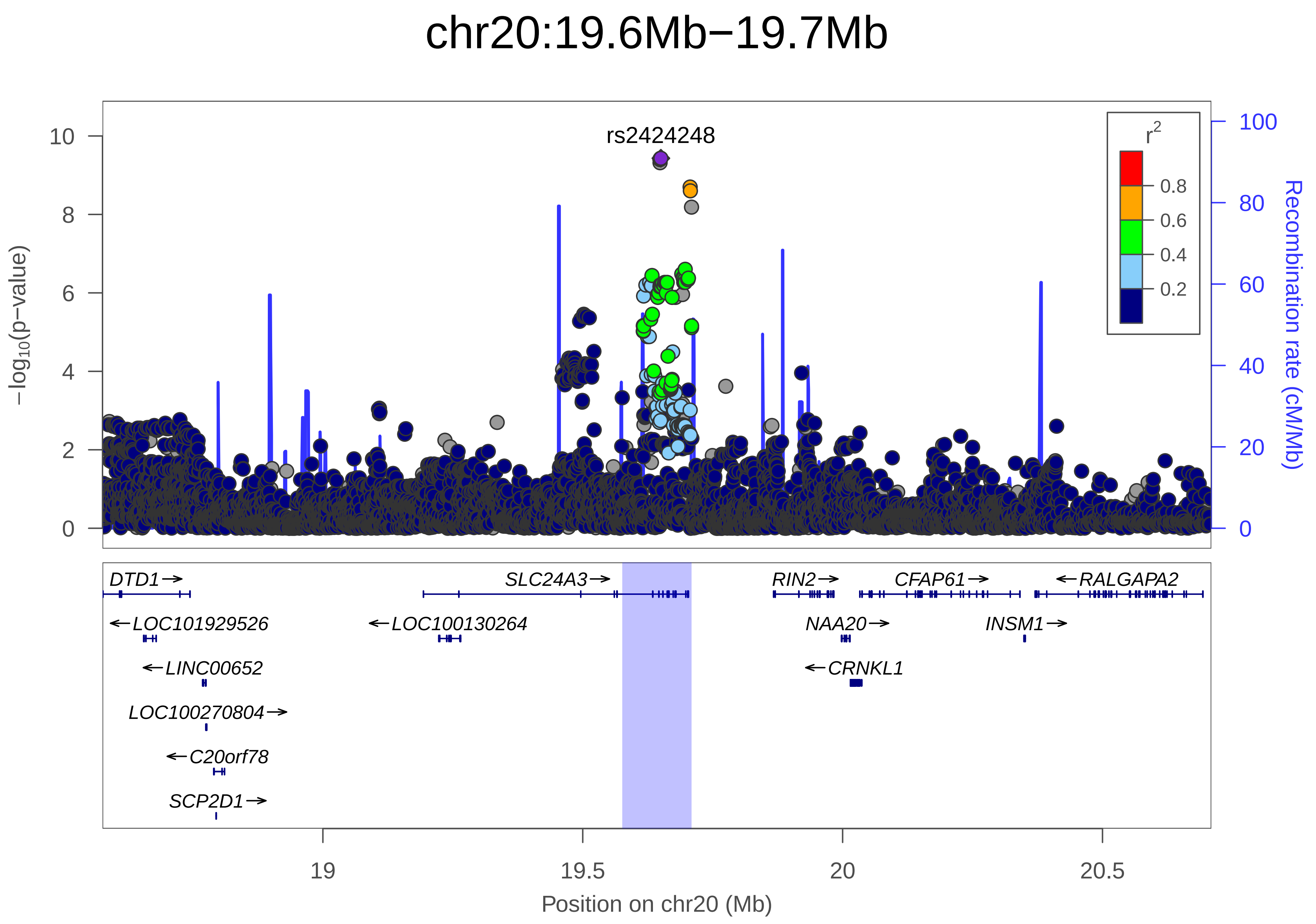

chr20:19.6Mb−19.7Mb
rs2424248
−log10(p−value)
0
2
4
6
8
10
Recombination rate (cM/Mb)
0
20
40
60
80
100
r2
0.2
0.4
0.6
0.8
DTD1→
SLC24A3→
RIN2→
CFAP61→
←RALGAPA2
←LOC101929526
←LOC100130264
NAA20→
INSM1→
←LINC00652
←CRNKL1
LOC100270804→
←C20orf78
SCP2D1→
19
19.5
20
20.5
Position on chr20 (Mb)

| | |
|---|---|
| date: | Mon Dec 10 08:03:18 2018 |
| build: | hg19 |
| display range: | chr20:18575990−20709392 [18575990−20709392] |
| hilite range: | 19.576Mb − 19.709Mb [ 19576000 − 19709000 ] |
| reference SNP: | chr20:19650324 |
| number of SNPs plotted: | 7522 |
| min P−value: | 3.7E−10 [chr20:19649716] |
| max P−value: | 1E0 [chr20:18934979] |

Make more plots at http://csg.sph.umich.edu/locuszoom/
