## Supplementary Figure 49 for "Genome-wide Association Study of Multisite Chronic Pain in UK Biobank"

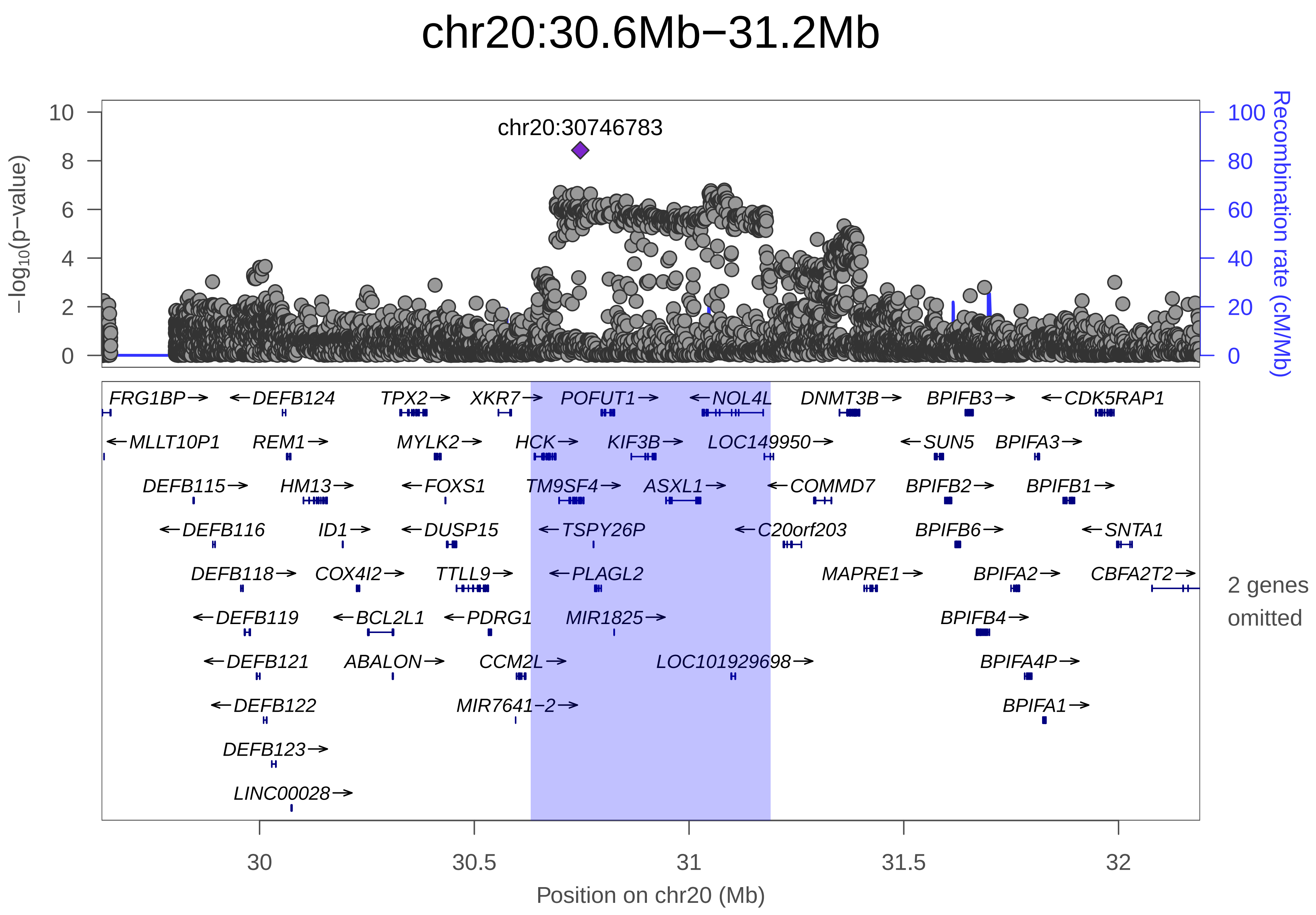

chr20:30.6Mb−31.2Mb
chr20:30746783
−log10(p−value)
Recombination rate (cM/Mb)
0
2
4
6
8
10
0
20
40
60
80
100
FRG1BP→
←DEFB124
TPX2→
XKR7→
POFUT1→
←NOL4L
DNMT3B→
BPIFB3→
←CDK5RAP1
←MLLT10P1
REM1→
MYLK2→
HCK→
KIF3B→
LOC149950→
←SUN5
BPIFA3→
DEFB115→
HM13→
←FOXS1
TM9SF4→
ASXL1→
←COMMD7
BPIFB2→
BPIFB1→
←DEFB116
ID1→
←DUSP15
←TSPY26P
←C20orf203
BPIFB6→
←SNTA1
DEFB118→
COX4I2→
TTLL9→
←PLAGL2
MAPRE1→
BPIFA2→
CBFA2T2→
2 genes omitted
←DEFB119
←BCL2L1
←PDRG1
MIR1825→
BPIFB4→
←DEFB121
ABALON→
CCM2L→
LOC101929698→
BPIFA4P→
←DEFB122
MIR7641−2→
BPIFA1→
DEFB123→
LINC00028→
30
30.5
31
31.5
32
Position on chr20 (Mb)

| | |
|---|---|
| date: | Mon Dec 10 08:33:54 2018 |
| build: | hg19 |
| display range: | chr20:29632082−32189993 [29632082−32189993] |
| hilite range: | 30.632Mb − 31.19Mb [ 30632000 − 31190000 ] |
| reference SNP: | chr20:30746783 |
| number of SNPs plotted: | 6631 |
| min P−value: | 3.7E−9 [chr20:30746783] |
| max P−value: | 1E0 [chr20:29634102] |
| omitted Genes: | HM13−AS1, MIR3193 |
| Warning: | No usable LD information for reference SNP. |

Make more plots at http://csg.sph.umich.edu/locuszoom/
