## Supplementary figures and images for "Genome-wide Association Study of Multisite Chronic Pain in UK Biobank"

### Supplementary Figure 1

Supplementary Figure 1: MDD Exposure QQ Plots

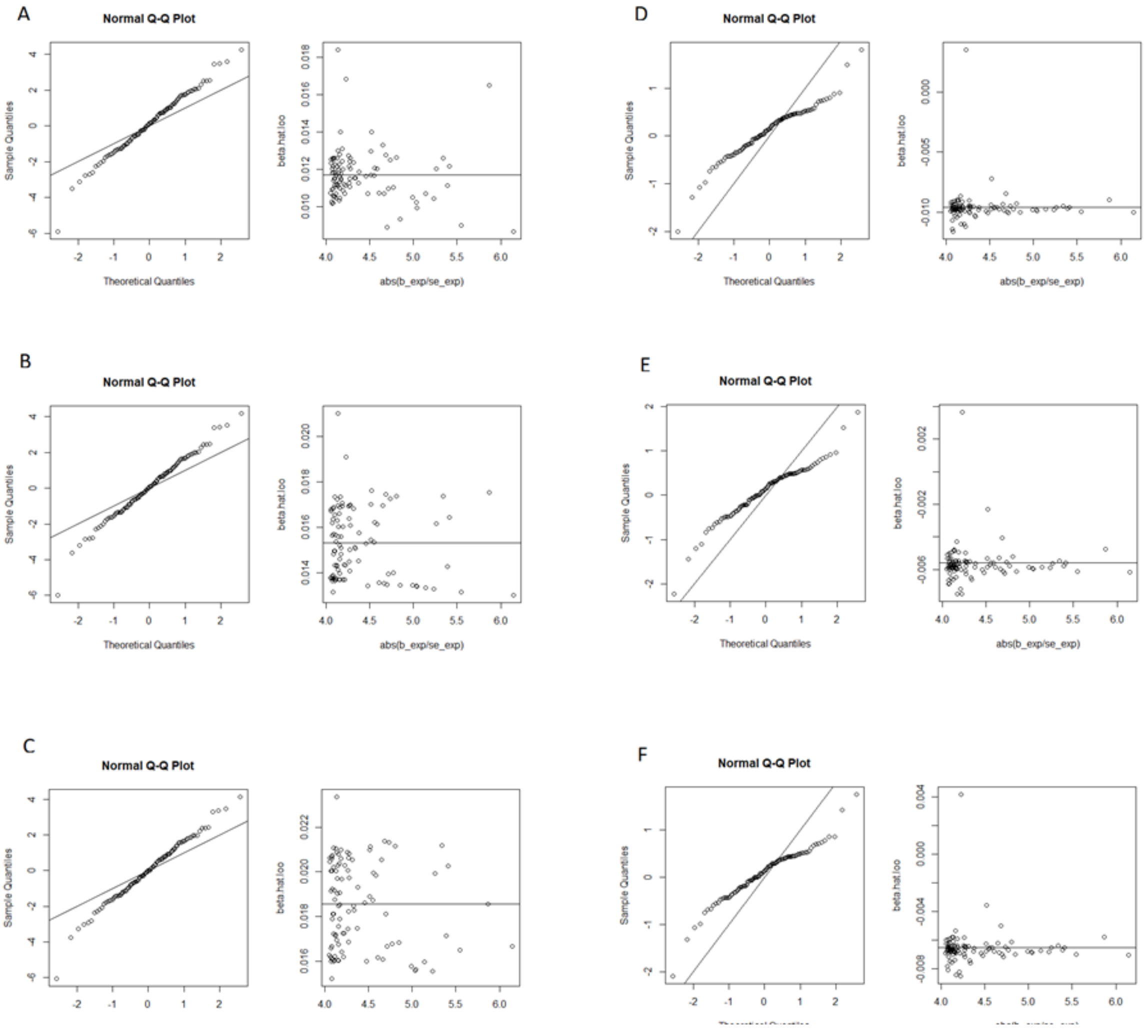

### Supplementary Figure 2

Supplementary Figure 2: MCP Exposure QQ Plots

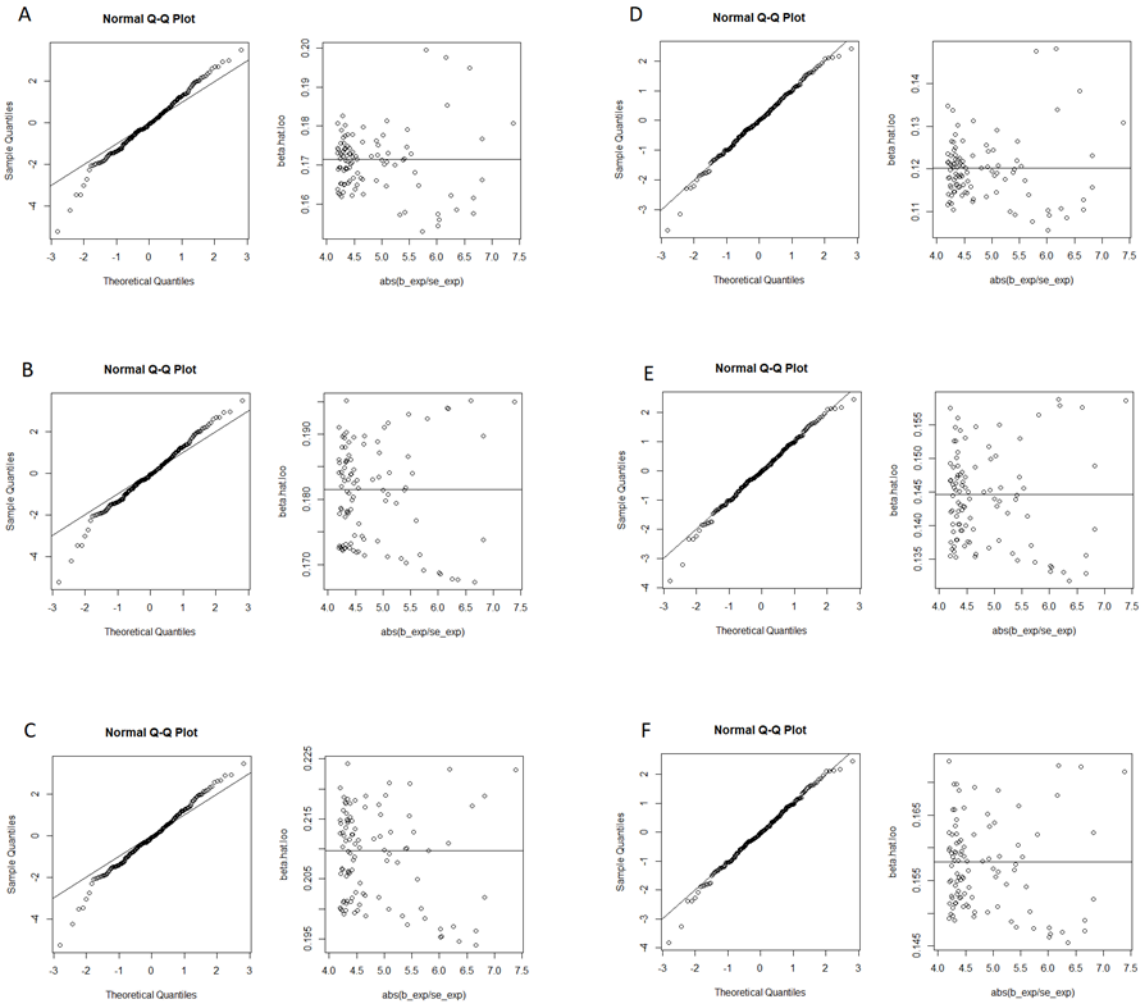
