## Supplementary Table 1 for "Genome-wide Association Study of Multisite Chronic Pain in UK Biobank"

**Supplementary Table 1: Model Types Fitted by MR-RAPS**

| Overdispersion | Loss Function | Underlying Model | Results-Section Figure Label |
|---|---|---|---|
| FALSE | l2 | No systematic pleiotropy, no idiosyncratic pleiotropy | A |
| FALSE | huber | No systematic pleiotropy, idiosyncratic pleiotropy | B |
| FALSE | tukey | No systematic pleiotropy, idiosyncratic pleiotropy | C |
| TRUE | l2 | systematic pleiotropy, no idiosyncratic pleiotropy | D |
| TRUE | huber | systematic pleiotropy, idiosyncratic pleiotropy | E |
| TRUE | tukey | systematic pleiotropy, idiosyncratic pleiotropy | F |
